# Using a reverse genetics system to generate recombinant SARS-CoV-2 expressing robust levels of reporter genes

**DOI:** 10.1101/2022.05.21.492922

**Authors:** Chengjin Ye, Luis Martinez-Sobrido

## Abstract

Reporter-expressing recombinant virus represents an excellent option and a powerful tool to investigate, among others, viral infection, pathogenicity, and transmission, as well as to identify therapeutic compounds that inhibit viral infection and prophylactic vaccines. To combat the still ongoing coronavirus disease 2019 (COVID-19) pandemic, we have established a robust bacterial artificial chromosome (BAC)-based reverse genetics (RG) system to rapidly generate recombinant severe acute respiratory syndrome coronavirus 2 (rSARS-CoV-2) to study the contribution of viral proteins in viral pathogenesis. In addition, we have also engineered reporter-expressing recombinant viruses in which we place the reporter genes upstream of the viral nucleocapsid (N) gene to promote high levels of reporter gene expression that facilitates the study of SARS-CoV-2 *in vitro* and *in vivo*. Although successful, the genetic manipulation of the BAC containing the entire SARS-CoV-2 genome of ∼30,000 nucleotides, is challenging. Herein, we depict the technical details to engineer rSARS-CoV-2 expressing reporter genes using the BAC-based RG approach. We describe i) assembly of the full-length (FL) SARS-CoV-2 genome sequences into the empty pBeloBAC, ii) verification of the pBeloBAC-FL, iii) cloning of a Venus reporter gene into the pBeloBAC-FL, and iv) recovery of the Venus-expressing rSARS-CoV-2. By following this protocol, researchers with basic molecular biology and gene engineering techniques knowledge will be able to generate wild-type and reporter-expressing rSARS-CoV-2.

## INTRODUCTION

Coronaviruses (CoVs) are enveloped, single-stranded, positive-sense RNA viruses, belonging to the Nidovirales order, that are responsible for causing seasonal mild-respiratory illness (e.g., 229E, NL63, OC43, and HKU1) or fatal disease (e.g., SARS-CoV and MERS-CoV) in humans ^1–3^. The emergence of severe acute respiratory syndrome coronavirus 2 (SARS-CoV-2) in December of 2019 has initiated a worldwide pandemic of CoV disease 2019 (COVID-19) that is still threatening global public health ^4–7^. To investigate this newly emerged virus, we have established a robust bacterial artificial chromosome (BAC)-based reverse genetics (RG) system for SARS-CoV-2 ^8^. By using this system, we were able to generate recombinant (r)SARS-CoV-2 where individual accessory open reading frame (ORF) proteins were deleted from the viral genome ^9^. Moreover, by replacing the viral ORF7a gene with a reporter gene, we have engineered rSARS-CoV-2 expressing fluorescent (mCherry or Venus) or bioluminescent (nanoluciferase, Nluc) reporter genes that can be used for the screening of compounds with antiviral activity *in vitro* ^10^. However, this approach based on the substitution of the viral ORF7a with the reporter gene resulted in low levels of reporter gene expression, making difficult to detect viral plaques with a ChemiDoc imaging system *in vitro* or the presence of the virus in K18 human angiotensin converting enzyme (hACE2) transgenic mice using *in vivo* imaging systems (IVIS). To overcome this limitation, we developed a novel strategy in which we place reporter genes upstream of the viral nucleocapsid (N) gene using the 2A autoproteolytic cleavage site of the porcine tescherovirus (PTV1) ^11^. Since the viral N gene is one of the highest expressed viral genes during SARS-CoV-2 infection, reporter gene expression using this new strategy resulted in ∼30-fold increase compared with the ORF7a-replacement strategy, allowing the detection of viral infection in cultured cells using a ChemiDoc imaging system or in K18 hACE2 mice using IVIS ^11^. Collectively, this approach offers researchers a BAC-based RG approach to generate rSARS-CoV-2 expressing robust levels of reporter genes from the locus of the N gene that does not require the deletion of any viral genes, and therefore, a useful tool to investigate SARS-CoV-2 in cultured cells or in validated animal models to assess the efficacy of vaccines or to identify prophylactic antivirals to combat SARS-CoV-2.

### Development of the protocol

RG represents a powerful tool to answer important questions about viral infection, transmission, and pathogenesis. In addition, RG approaches have been used to develop attenuated forms of viruses that can be used as live-attenuated vaccines (LAV) for the prevention of viral infections. Likewise, similar RG systems have been used to genetically manipulate the viral genome to express foreign genes, including those encoding reporter fluorescent or luciferase proteins, to establish viral inhibition assays to facilitate the identification of biologicals affecting viral infection, including antivirals and antibodies ^10^.

Two reports initially described the assembly of the entire SARS-CoV-2 genome by *in vitro* ligation ^12^ or homologous recombination in yeast ^13^. However, both approaches rely on the *in vitro* transcription of the viral genome using a bacteriophage T7 promoter, a process that requires careful control of experimental variables and extraordinary technical expertise, and is therefore restricted to only a few laboratories. To overcome this limitation, we developed a BAC-based RG system for SARS-CoV-2 ^8^. It applies the same principles as previously established to develop RG for other CoVs ^14–16^, and other single-stranded positive-sense RNA viruses ^17, 18^. After analyzing the complete genome sequences of SARS-CoV-2 and selecting appropriate restriction sites, we designed and chemically synthesized a set of five fragments (F1-F5) covering the entire viral genome ^8^. To assemble the entire SARS-CoV-2 genome, F1 was first assembled into the empty pBeloBAC and used as the backbone for assembling the other F2-F5 fragments ^8^. The F1 fragment includes the first 673 nucleotides of the SARS-CoV-2 genome under the control of a cytomegalovirus (CMV) immediate-early promoter, a multiple-cloning site containing the restriction sites selected for the sequential assembly of F2-F5, and the last 4,585 nucleotides of the SARS-CoV-2 genome, followed by the hepatitis delta virus ribozyme (Rz) and the bovine growth hormone (bGH) termination and polyadenylation sequences ^8^. This method has been described previously by our laboratory, showing the BAC RG approach gives rapid recombinant virus rescue by using a simple and conventional cell transfection technique ^8^. With this method, we have also been able to generate a stable rSARS-CoV-2 expressing robust levels of reporter genes ^11^. However, because the pBeloBAC that harbors all elements required for efficient viral rescue is ∼38,000 base pairs long, it is challenging to manipulate using conventional molecular biology methods and techniques. To assist other researchers to overcome challenges with the use of the SARS-CoV-2 BAC-based RG system, we describe here all the technical information and detailed protocols necessary for the assembly of the SARS-CoV-2 entire genome into the pBeloBAC, and its manipulation to generate rSARS-CoV-2 expressing robust levels of the reporter gene Venus. Importantly, this protocol could be employed by other researchers to generate wild-type (WT), reporter-expressing rSARS-CoV-2, and/or viral mutants of interest by following the similar experimental procedures and approaches.

### Applications of the method

The key features of this protocol are the assembly of large DNA fragments into the empty pBeloBAC and the genetic manipulation of the pBeloBAC and its derivates. In this protocol, we describe the assembly of the entire viral genome of SARS-CoV-2 WA-1 strain, one of the largest RNA viral genomes, into the empty pBeloBAC and the generation of rSARS-CoV-2 expressing robust levels of a fluorescent Venus reporter gene. Moreover, similar experimental procedures can be used to delete ^9^, replace ^10^, or mutate ^19^ SARS-CoV-2 genome sequences, or to insert additional foreign genes. In addition, this protocol can be also applied to establish RG systems to generate recombinant forms of other non-segmented positive stranded RNA viruses, as well as viral genome sequences that are either unstable or unable to be genetically assembled and manipulated in other DNA vector backbones.

### Comparison with other methods

Our SARS-CoV-2 BAC-based RG system has several advantages over alternative methods. First, our RG approach based on the use of a BAC DNA does not rely on *in vitro* RNA transcription using a bacteriophage T7 promoter and RNA electroporation. Second, our BAC DNA transfection protocol based on the use of Lipofectamine 2000 is less technically challenged than to electroporate cells with RNA molecules produced *in vitro* to recover rSARS-CoV-2. Third, once the entire SARS-CoV-2 genome sequences are successfully assembled into the pBeloBAC, viral fragments do not need to be prepared each time. Fourth, the BAC DNA is highly stable, which allows for sharing and transport between laboratories. Last, but not least, the levels of reporter gene expression using our innovative PTV1 2A approach from the locus of the viral N gene are dramatically higher than those obtained by expressing the reporter gene from the locus of the viral ORF7a ^10^. Importantly, our approach does not require substituting a viral gene by the reporter gene, contrary to the previous strategy to express foreign genes by substituting them for the viral ORF7a gene ^10, 11^.

However, assembling the entire viral genome into the empty pBeloBAC for successful rescue of SARS-CoV-2 is time consuming. To overcome this limitation, F2-F5 fragments could be assembled into the pBeloBAC-F1 simultaneously, rather than stepwise. Since *in vitro* ligation of seven viral fragments has been previously described in the literature ^20^, simultaneous assembly of F2-F5 into the pBeloBAC-F1 would be feasible. Alternatively, assembly of the F2-F5 fragments into the pBeloBAC-F1 could be accomplished by using a yeast recombination-based system, as previously described^13^. Another limitation of our RG system is that for successful results, is the yield of the pBeloBAC plasmid that is maintained as a single copy in each bacterium. To obtain sufficient pBeloBAC plasmid for a one-time rescue by transfection into Vero E6 cells using Lipofectamine 2000, we suggest to prepare the pBeloBAC plasmid from a minimum volume of 25 ml of overnight cultured bacteria.

### Experimental design

To clone the SARS-CoV-2 genome into the empty pBeloBAC, we analyzed the entire viral genome sequence of the SARS-CoV-2/human/USA/WA-CDC-02982586-001/2020 (GenBank accession no. MN985325) for unique restriction sites using the online NEBcutter program (http://nc2.neb.com/NEBcutter2/) and designed 5 viral fragments (F1-F5) with ∼5,000-8,000 base pairs in length (**Fig. 1a**). Some restriction sites that have dual cutters might need to be manipulated for cloning purposes by replacing them with silent mutations that can be used as genetic tags as well to distinguish between rSARS-CoV-2 generated with RG and the natural SARS-CoV-2 WA-1. In our case, we mutated a MluI restriction site in the Membrane (M) gene in the synthesized F1 and a BstBI restriction site in the Spike (S) gene in the synthesized F5 to facilitate the cloning work (**Supplementary Sequence Archive**).

**Figure 1.**
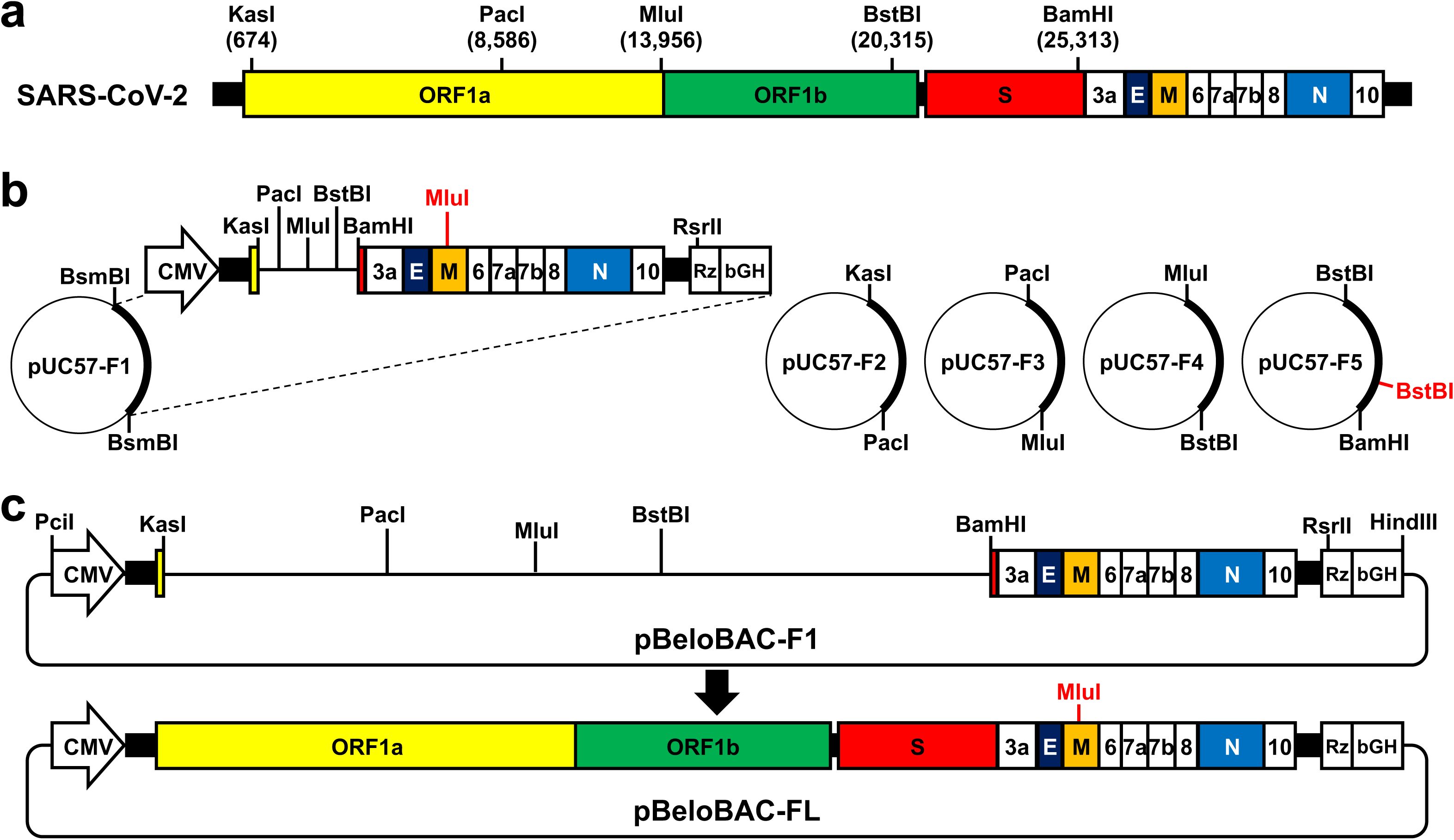
Overview of a BAC-based RG system for generation of rSARS-CoV-2. a. The schematic representation of the SARS-CoV-2 genome. Unique restriction sites used for vial genome assembly are indicated. b. Commercially synthesized five fragments (F1-F5) in pUC57 plasmids. Restriction sites used to release each of the viral fragments are indicated. The MluI and BstBI highlighted in red are the restriction sites that have been removed by silent mutation and used as genetics tags. c. Assembly of the entire SARS-CoV-2 genome into the empty pBeloBAC to generate the full-length rescue plasmid (pBeloBAC-FL).

It is also imperative to consider that the restriction sites used should be unique in the empty pBeloBAC. If a limited number of unique restriction sites are available for efficient and convenient cloning, one potential strategy is to incorporate type IIS restriction sites. The type IIS restriction enzymes recognize asymmetric DNA sequences and could generate compatible overhangs with unique restriction sites. We incorporated this strategy in our protocol: two BsmBI restriction sites were included in the 5’ and 3’ ends of the F1 to generate compatible overhangs with those of PciI and HindIII (**Supplementary Sequence Archive**).

After completing the design of the five fragments, they can be chemically synthesized using any commercial synthesis service and cloned in an appropriate shuttle vector (e.g., pUC57) (**Fig. 1b**) before being assembled into the empty pBeoBAC (**Fig. 1c**). Then, the fragments in the shuttle vectors were released and recovered (**Fig. 2a**). and the pBeloBAC-F1 positive colonies were selected by colony PCR (**Fig. 2b**) and confirmed by Sanger sequencing. Afterward, F2-F5 were sequentially cloned into the pBeloBAC-F1 using KasI and PacI (F2), PacI and MluI (F3), MluI and BstBI (F4), and finally, BstBI and BamHI (F5). Presence of all the F2-F5 was confirmed with restriction digestions (**Fig. 2c-f**). After confirming that the pBeloBAC contained all five fragments (pBeloBAC-FL) by restriction digestion, the pBeloBAC DNA plasmid was prepared using maxiprep for transfection, using Lipofectamine 2000, of Vero E6 cells (**Fig. 3a**). The ability of using the pBeloBAC-FL plasmid to generate rSARS-CoV-2 was confirmed by assessing cytopathic effect (CPE) in transfected Vero E6 cells (**Fig. 3b**), by immunofluorescent assay (IFA) (**Fig. 3c**), and by Sanger sequencing of reverse-transcription (RT)-PCR products (**Fig. 3d**).

**Figure 2.**
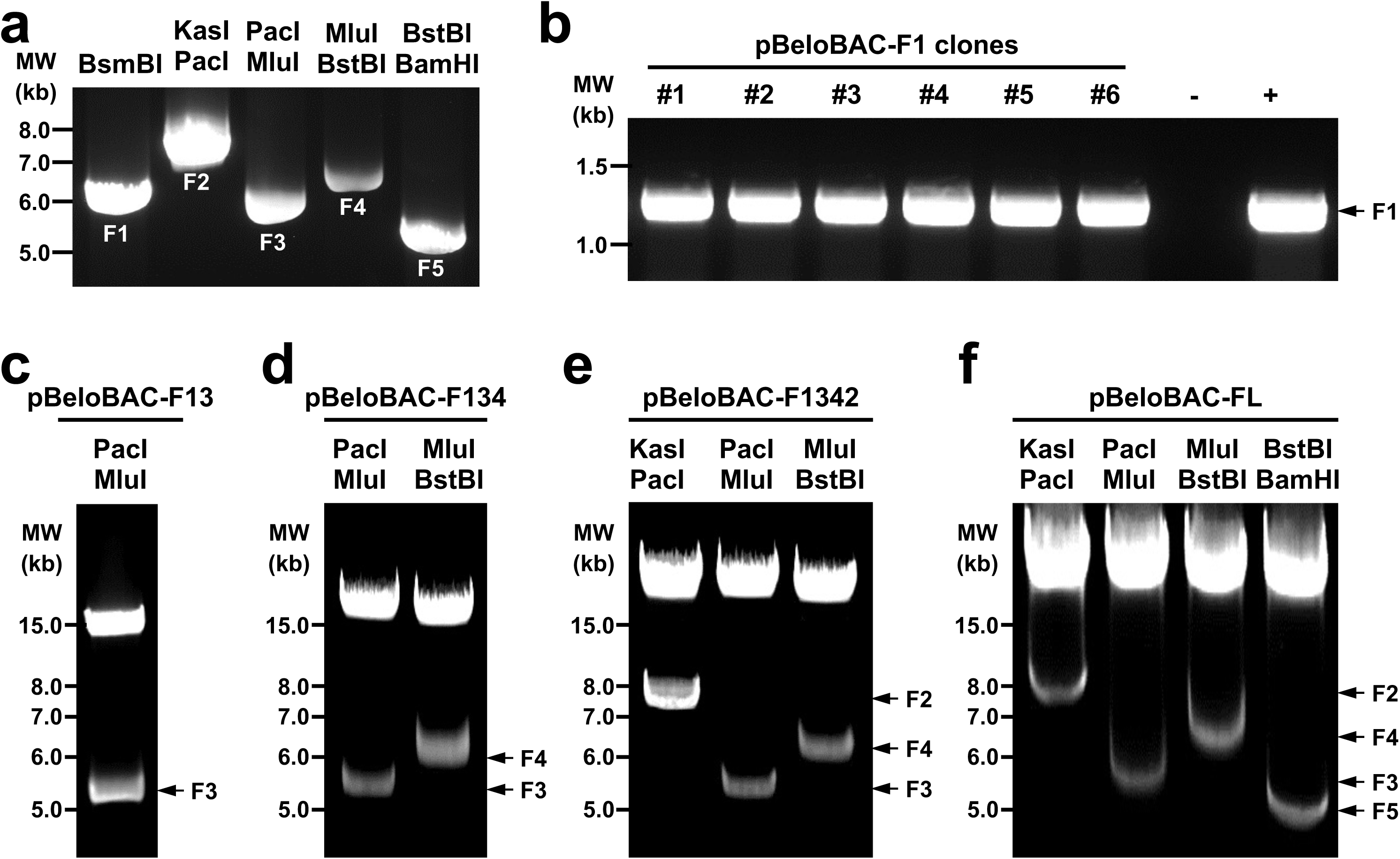
Assembly of entire SARS-CoV-2 genome into a BAC. a. The released five fragments (F1-F5) from pUC57 plasmids. b. Colony PCR screening for positive pBeloBAC-F1. c. Restriction digestion analysis of pBeloBAC-F13 using PacI and MluI. d. Restriction digestion analysis of pBeloBAC-F134 using PacI and MluI (F3); and MluI and BstBI (F4). e. Restriction digestion analysis of pBeloBAC-F1342 using KasI and PacI (F2); PacI and MluI (F3); and MluI and BstBI (F4). f. Restrict digestion confirmation of the pBeloBAC-FL using KasI and PacI (F2); PacI and MluI (F3); MluI and BstBI (F4); and BstBI and BamHI (F5).

**Figure 3.**
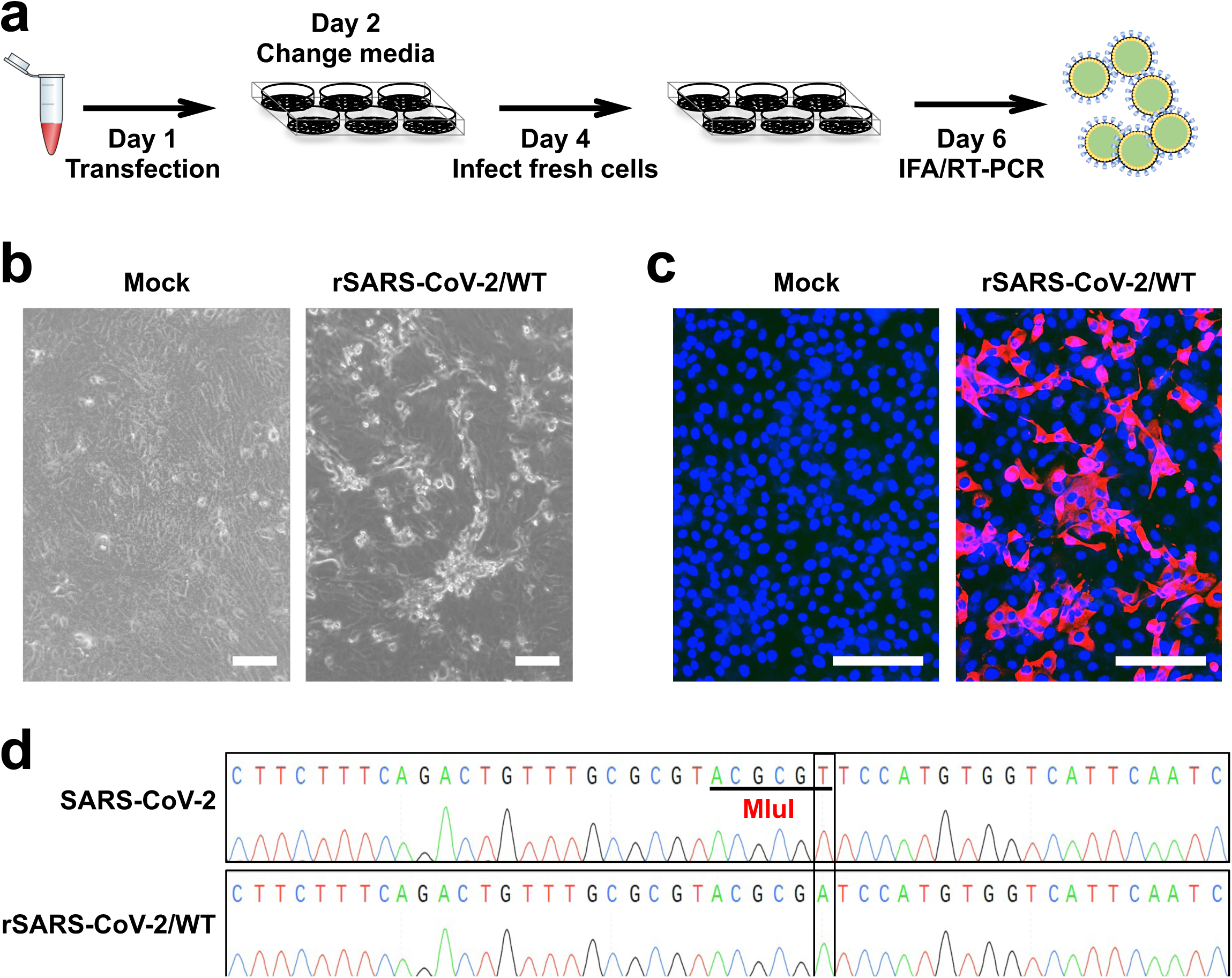
Recovery of rSARS-CoV-2 from the pBeloBAC-FL. a. Protocol used for recovery of rSARS-CoV-2 from the pBeloBAC-FL using Lipofectamine 2000 transfection in Vero E6 cells. b. CPE caused by rSARS-CoV-2/WT in Vero E6 cells. Scale bars, 100 µm. c. Confirmation of the rescued rSARS-CoV-2/WT by IFA using a mouse antibody against viral N protein and a TRITC labeled donkey anti-mouse IgG secondary antibody. The nucleus was stained by DAPI. d. Sanger sequencing of the M gene of the natural isolate SARS-CoV-2 (top) and the recombinant SARS-CoV-2(bottom) that the MluI was removed by silent mutation.

Next, the Venus reporter gene was introduced into the pUC57-F1 (**Fig. 4a**) before being subcloned into the pBeloBAC-FL (**Fig. 4b**). To that end, the Venus-2A and the entire pUC57-F1 were amplified by PCR and reverse PCR (**Fig. 5a**), respectively, and then they were ligated using compatible overhangs generated by BsaI restriction digestion to achieve a scarless pUC57-F1/Venus-2A without introducing any extra sequences. To generate the pBeloBAC-FL/Venus-2A, the F1 containing Venus-2A was released and subcloned into the pBeloBAC-FL using the restriction sites of BamHI and RsrII (**Fig. 5b**). Finally, positive pBeloBAC-FL/Venus-2A clones were selected by colony PCR (**Fig. 5c**), and maxiprep plasmids were prepared and transfected into Vero E6 cells using Lipofectamine 2000 to generate rSARS-CoV-2/Venus-2A. The expression of Venus in the Vero E6 cells that infected with the supernatant of the transfected cells confirmed the rescue of rSARS-CoV-2/Venus-2A (**Fig. 6a**). The successful detection of fluorescent plaques under a ChemiDoc imaging system further confirmed that the rescued rSARS-CoV-2/Venus-2A expressing a robust level of Venus fluorescent expression in all the viral plaques by comparing to those detected by immunostaining using an antibody against the viral N protein (**Fig. 6b**).

**Figure 4.**
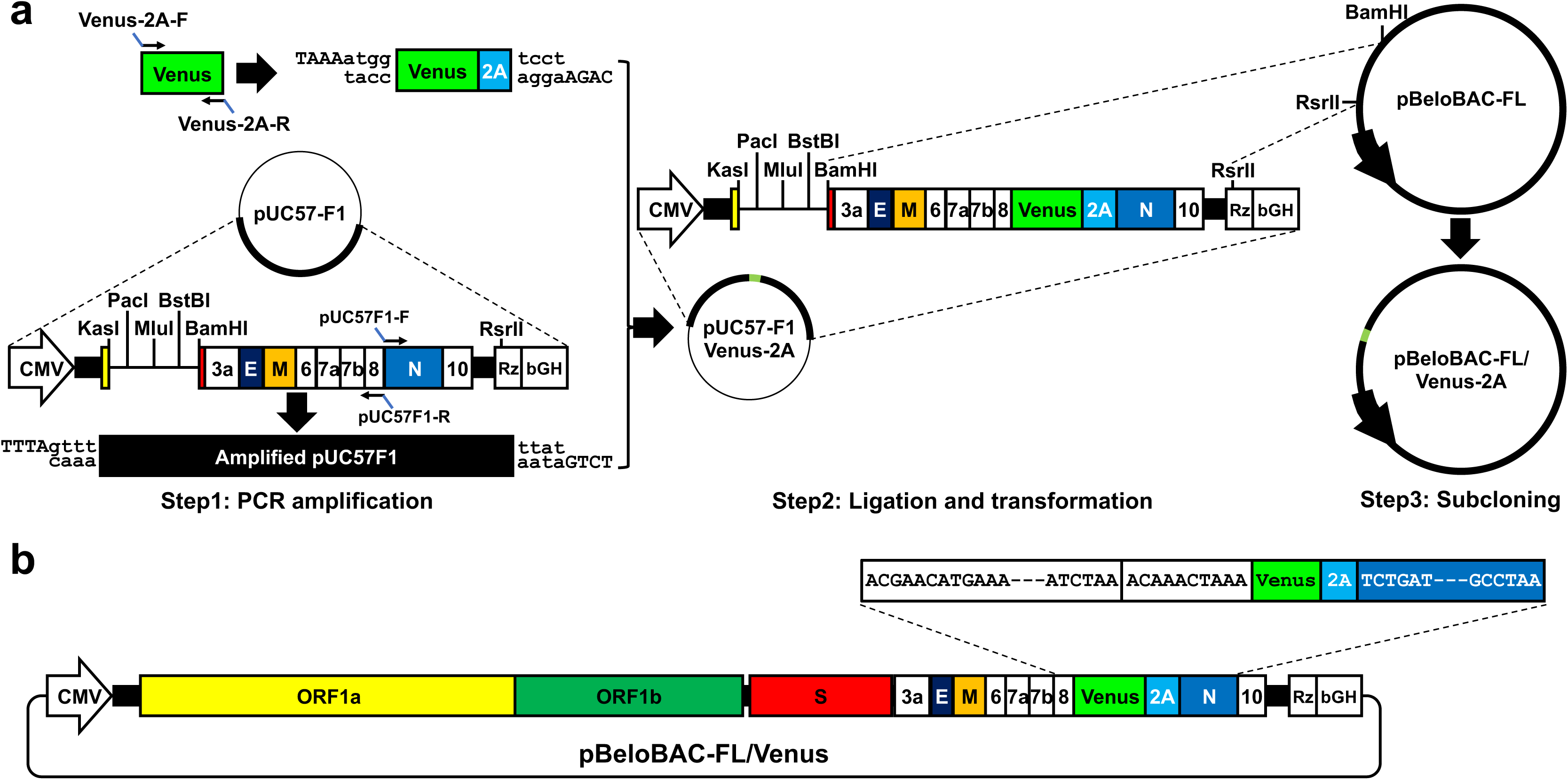
Overview of the BAC to generate rSARS-CoV-2 expressing Venus from the N protein locus. a. Schematic representation of the experimental flow to assemble the Venus into the pUC57-FA and subcloned into the pBeloBAC-FL. b. Schematic scheme of the pBeloBAC-FL/Venus-2A.

**Figure 5.**
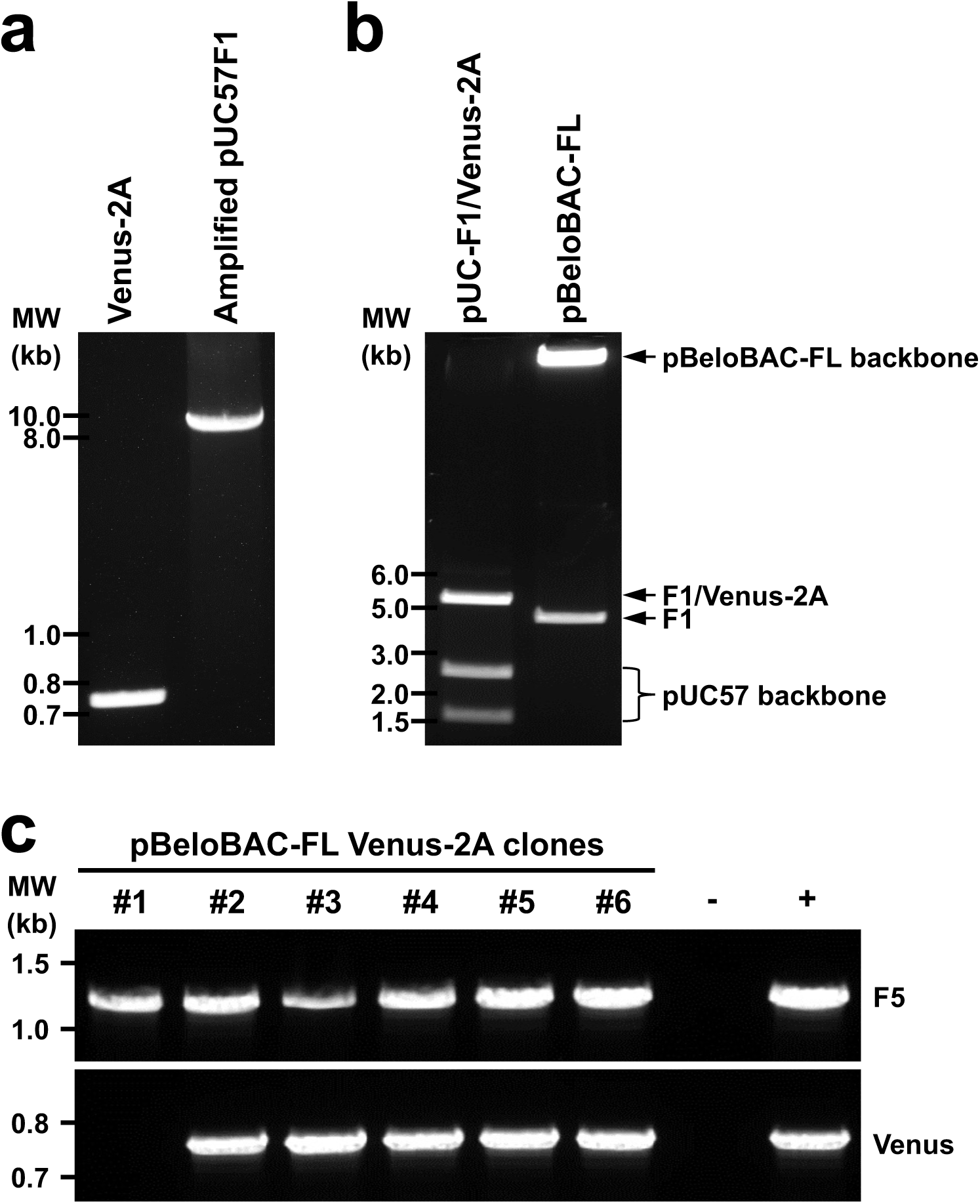
Assembly of F1 containing Venus-2A into the BAC for generation of rSARS-CoV-2/Venus-2A. a. PCR amplification of Venus-2A and pUC57F1. b. Purification of the F1/Venus-2A released from the pUC57-F1/Venus-2A; and the pBeloBAC-FL backbone after BamHI and RsrII digestion to remove the parental F1 segment for subcloning of the F1/Venus-2A. c. Colony PCR screening of pBeloBAC-FL/Venus-2A colonies.

**Figure 6.**
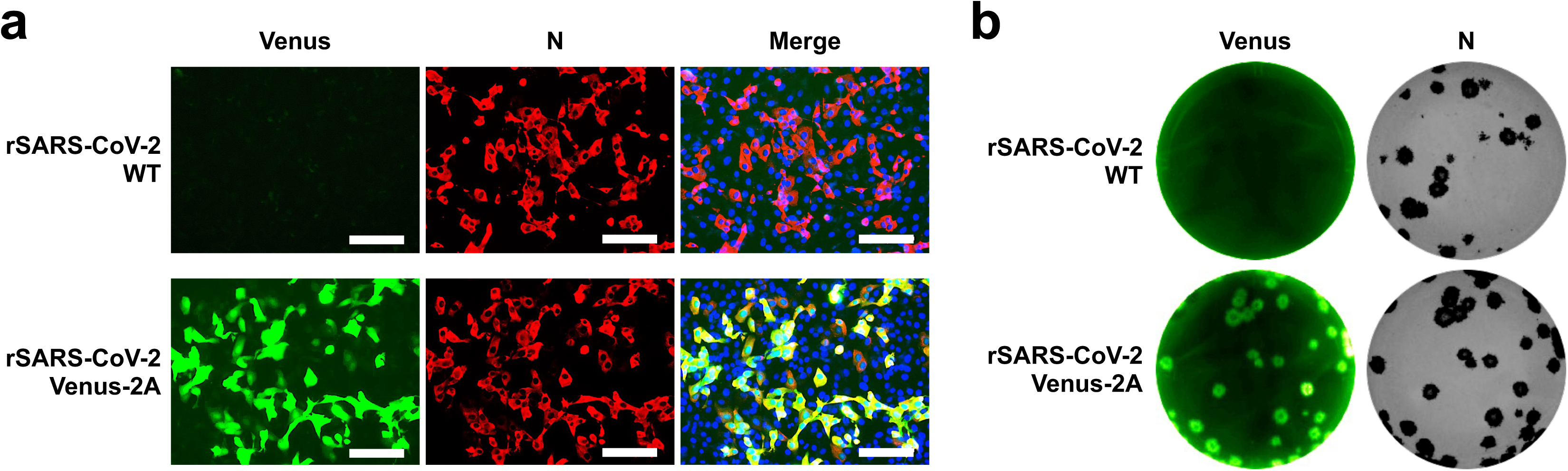
Characterization of rSARS-CoV-2/Venus-2A. a. Fluorescent (Venus) and IFA (N) confirmation of rSARS-CoV-2/Venus-2A rescue. rSARS-CoV-2/WT was included as an internal control. b. Detection of fluorescent plaques formed by rSARS-CoV-2/Venus-2A under a ChemiDoc imaging system. Immunostaining with the N protein 1C7C7 MAb of the plaques formed by rSARS-CoV-2/Venus-2A and rSARS-CoV-2/WT.

### Limitations

Our experimental design relies on the molecular genomic assemble of five fragments that span the entire SARS-CoV-2 genome into the empty pBeloBAC to establish an RG system for SARS-CoV-2. By using the pBeloBAC-FL, we introduce a Venus reporter gene upstream of the N gene to generate a rSARS-CoV-2 expressing robust levels of Venus (rSARS-CoV-2/Venus-2A). While the five viral fragments were commercially synthesized, it is also feasible they may be amplified from SARS-CoV-2 infected samples, or from purified SARS-CoV-2 RNA, using RT-PCR and then cloned into a plasmid vector. If so, it will be imperative to incubate the transformed bacteria and propagate the colonies at or below 25°C to avoid any potential deletions, insertions, or mutations arising from potential toxic effect of the viral sequences.

Also crucial for the successful assembly of all the five viral fragments into the pBeloBAC is the integrity of any DNA used. During DNA isolation using gel electrophoresis, only brief exposure the DNA to long-wavelength UV light (e.g., 360 nm) since DNA is very vulnerable to UV light, particularly short-wavelength UV light (e.g., 254, 302, or 312 nm). The integrity and degradation of the pBeloBAC plasmid must also be monitored closely since the pBeloBAC plasmid is prepared from a large volume of bacteria and bacterial enzymes are also isolated together with the pBeloBAC. We recommend that the pBeloBAC and its derivates to be stored at -80°C instead of classical plasmid storage of -20°C after being isolated and purified.

## MATERIALS

### Equipment

- -80°C Freezer (PHC Corporation, catalog #: MDF-DU900V).
- 0.2 mL PCR tube (Greiner bio-one, catalog #: 683201).
- 0.22 µm Syringe Filter Unit (Millipore, catalog #: SLGP033RS).
- 1.5 mL tube (Greiner bio-one, catalog #:616261).
- 10 mL Serological pipet (Greiner bio-one, catalog #: 607180).
- 10 μL Filtered tip (Thermo Fisher Scientific, catalog #: 94056980).
- 100 mm Petri dish (Fisher Scientific, catalog #: NC9463569).
- 1,000 μL Filtered tip (Thermo Fisher Scientific, catalog #: 94056710).
- 1,000 mL Conical flask (VWR, catalog #: 470332-218).
- 15 mL Tube (Greiner bio-one, catalog #: 188201).
- 2 mm Electroporation Cuvettes (Bio-Rad, catalog #: 1652092).
- 200 μL Filtered tip (Thermo Fisher Scientific, catalog #: 94056380).
- 25 mL Serological pipet (Greiner bio-one, catalog #: 760180).
- 250 mL Conical flask (VWR, catalog #: 470332-214).
- 250 mL Polycarbonate Centrifuge Bottles (Thermo Fisher Scientific, catalog#: 3122-0250).
- 250 mL Wheaton bottle (VWR, catalog #: 16159-856).
- 50 mL Tube (Greiner bio-one, catalog #: 227201)
- 500 mL Wheaton bottle (VWR, catalog #: 16159-889).
- 6-well plate (Greiner bio-one, catalog #: 657160).
- Bacterial culture tube (VWR, catalog #: 60818-689).
- Bacterial incubator (VWR, catalog #: 97025-630).
- Bunsen burner (VWR, catalog #: 89038-530).
- ChemiDoc imaging system (Bio-Rad, catalog #: 12003153).
- CO_2_ incubator (PHC Corporation, catalog #: MCO-170AICUVL-PA).
- Electroporator (Bio-Rad, catalog #: 1652100).
- EVOS Imaging System (Thermo Fisher Scientific, catalog #: AMF5000).
- Gel Electrophoresis System (Thermo Fisher Scientific, catalog #: A3-1).
- Large capacity refrigerated centrifuge (Thermo Fisher Scientific, catalog #: 12121680).
- Microcentrifuge (Thermo Fisher Scientific, catalog #: 75002440).
- Orbital shaker (Thermo Fisher Scientific, catalog #: SHKE4450).
- pH meter (Sartorius, catalog #: PYP102S).
- Pipette set (Gilson, catalog #: F167380).
- Power supply (Bio-Rad, catalog #: 1645050).
- Scale (Sartorius, catalog #: ENTRIS2202-1S).
- NanoDrop One Microvolume UV-Vis spectrophotometer (Thermo Fisher Scientific, catalog #: ND-ONE-W).
- Sterile single-use bottle top filters (Thermo Fisher Scientific, catalog #: 5954520).
- T225 flask (Thermo Fisher Scientific, catalog #: 159934).
- T75 flask (Thermo Fisher Scientific, catalog #: 156499).
- Thermal Cycler (G-Storm, catalog #: GS00482).
- Ultrapure Water System (Millipore, catalog #: ZIQ7000T0).
- Ultraviolet Transilluminator 365 nm (UVP, catalog #: 95-0459-01).
- Vacuum Manifold (Promega, catalog #: A7231).
- Water bath (VWR, catalog #: 76308-900).

### Cells

- DH10B electrocompetent cells (Thermo Fisher Scientific, catalog #: 18290015). CRITICAL Store at -80°C and avoid thaw-freeze cycles.
- NEB stable competent cells (New England Biolabs, catalog #: C3040H).
- Vero E6 cells (lab-maintained derivative from the American Type Culture Collection, ATCC, catalog #: CRL-1586). CRITICAL Ensure all the materials are exclusively used for maintaining this cell line.

### Reagents

- 0.05% Trypsin/0.53 mM EDTA (ethylenediaminetetraacetic acid) (Corning, catalog #: 25-051-CI).
- 0.4% Trypan blue (Corning, catalog #: 25-900-CI).
- 10% Formalin solution (Sigma-Aldrich, catalog #: HT501128-4L).
- 100x Penicillin-Streptomycin-L-Glutamine (PSG) (Corning, catalog #: 30-009-CI).
- 10,000x GelRed (Biotium, catalog #: 41003-1).
- 1 kb plus DNA ladder (Thermo Fisher Scientific, catalog #: 10787026).
- 35% Bovine Serum Albumin (BSA) solution (Sigma Aldrich, catalog #: A7409).
- 50x Tris-Acetate-EDTA (TAE) buffer (Fisher Scientific, catalog #: BP1332500).
- 6x Gel Loading Dye (New England Biolabs, catalog #: B7024S).
- ABC-HRP mouse IgG kit (Vector Laboratories, catalog #: PK-4002).
- Agar (OXOID, catalog #: LP0028).
- Agarose (Lonza, catalog #: 50004).
- Ampicillin (Corning, catalog #: 61-238-RM).
- ATP-Dependent DNase kit (Lucigen, catalog #: E3110K).
- BamHI-HF (New England Biolabs, catalog #: R3136L).
- BsaI (New England Biolabs, catalog #: R3733L).
- BsmBI (New England Biolabs, catalog #: R0739L).
- BstBI (New England Biolabs, catalog #: R0519L).
- Cell culture water (Thermo Fisher Scientific, catalog #: A1287301).
- Chloramphenicol (Corning, catalog #: 61-239-RI).
- Chloroform (Fisher Scientific, catalog #: AAJ67241AP).
- DAB substrate kit (Vector Laboratories, catalog #: SK-4100).
- DAPI (diamidino-2-phenylindole) Solution (1 mg/mL) (Thermo Fisher Scientific, catalog#: 62248)
- DEAE-Dextran (MP Biomedicals, catalog #: 195133).
- Dulbecco’s Modified Eagle Medium/Nutrient Mixture F-12 (DMEM/F-12) powder (Thermo Fisher Scientific, catalog #: 12400024).
- DMEM (Corning, catalog #: 15-013-CM).
- Ethanol (Sigma-Aldrich, catalog #: 459836-2L).
- Expand High Fidelity PCR System (Sigma-Aldrich, catalog #: 11759078001).
- Fetal Bovine Serum (FBS) (VWR, catalog #: 97068-085).
- First-Strand Synthesis System for RT-PCR (Thermo Fisher Scientific, catalog #: 11904018).
- Gel extraction kit (Qiagen, catalog #: 20021).
- Glacial acetic acid (Fisher Scientific, catalog #: 18-600-204).
- HindIII (New England Biolabs, catalog #: R0104L).
- Isopropanol (Fisher Scientific, catalog #: BP26181).
- KasI (New England Biolabs, catalog #: R0544L).
- Luria-Bertani (LB) agar (BD, catalog #: 214010).
- Lipofectamine 2000 Reagent (Thermo Fisher Scientific, catalog #: 11668019).
- MluI-HF (New England Biolabs, catalog #: R3198L).
- Mouse anti-N protein monoclonal antibody (kindly provide by Professor Thomas Moran from Icahn School of Medicine at Mount Sinai or commercially available from Sigma Aldrich, catalog #: ZMS1075).
- NEB buffer set (New England Biolabs, catalog #: B7200S).
- Nuclease-Free Water (Promega, catalog #: P1193).
- Opti-MEM (Thermo Fisher Scientific, catalog #: 31985-070).
- PacI (New England Biolabs, catalog #: R0547L).
- PciI (New England Biolabs, catalog #: R0655L) .
- PCR clean-up system (Promega, catalog #: A9285).
- Plasmid maxiprep kit (Omega, catalog #: D6922-04).
- Plasmid miniprep kit (Omega, catalog #: D6942-02).
- Phosphate Buffered Saline (PBS) (Corning, catalog #: 21-040-CM).
- RsrII (New England Biolabs, catalog #: R0501L).
- Shrimp Alkaline Phosphatase (rSAP) (New England Biolabs, catalog #: M0371L).
- SOC (Super Optimal broth with Catabolite repression) medium (Thermo Fisher Scientific, catalog #: 15544304).
- Sodium acetate (Fisher Scientific, catalog #: BP333-500).
- Sodium bicarbonate (Fisher Scientific, catalog #: S233-500).
- Sodium chloride (Fisher Scientific, catalog #: S271-1).
- T4 DNA ligase and buffer (New England Biolabs, catalog #: M0202L).
- TRITC (Tetramethylrhodamine) labeled donkey anti-mouse IgG secondary antibody (Thermo Fisher Scientific, catalog #: A16016).
- Triton X-100 (Sigma-Aldrich, catalog #: X100-500ML).
- TRIzol (Thermo Fisher Scientific, catalog #: 15596018).
- Tryptone (Thermo Fisher Scientific, catalog #: 211699).
- Yeast extract (BD, catalog #: 212720).

### Reagent setup

#### 0.5% Triton X-100 solution

Add 50 mL Triton X-100 to 950 mL PBS, mix thoroughly by stirring at room temperature for 1 h, and store at 4°C.

#### 1% DEAE-Dextran

Add 1 g of DEAE-Dextran powder to 100 mL cell culture water and sterilize by autoclaving at 121°C for 30 min.

#### 1x TAE buffer

Mix 100 mL of 50x TAE solution with 4,900 mL ultrapure deionized water in a sanitized carboy and store at room temperature.

#### 2x DMEM/F-12 solution

Dissolve 1 bag of DMEM/F-12 powder and 1.2 g sodium bicarbonate in 500 mL cell culture water, mix with 10 mL of PSG, 6 mL of 35% BSA solution, and sterilize by passing through a sterile single-use bottle top filter. CAUTION Keep sterile and store at 4°C.

#### 2% agar

Add 2 g of Oxoid agar to 100 mL cell culture water in a 250 mL Wheaton glass bottle. Autoclave at 121°C for 30 min and store at room temperature. CAUTION The 2% agar will solidify after cooling down. Microwave for 1-3 min to completely melt before using in the plaque assays.

#### 75% ethanol

Mix 150 mL of absolute ethanol and 50 mL of nuclease-free water in a 250 mL Wheaton glass bottle. Store at room temperature. CAUTION 75% ethanol is flammable, keep away from fire.

#### Ampicillin stock (150 mg/mL)

Add 1.5 g ampicillin to 10 mL of deionized water and mix thoroughly until the powder is fully dissolved in a 50 mL tube. Sterilize the solution by passing through a 0.22 μm Syringe Filter Unit and dispense into 0.5 mL aliquots in 1.5 mL tubes. Store the aliquots at -20°C.

#### Blocking solution

Add 77 mL of 35% BSA solution to 1 L of PBS and mix completely. Store at 4°C. The blocking solution is also used to dilute primary and secondary antibodies.

#### Chloramphenicol stock (12.5 mg/mL)

Add 0.25 g chloramphenicol to 20 mL absolute ethanol in a 50 mL tube, vortex until the powder is completely dissolved. Aliquot the solution into 1.5 mL tubes containing 0.5 mL per tube and store at -20°C. CAUTION Ethanol is flammable, keep away from fire.

#### Complete cell culture media

Add 110 mL FBS and 11 mL 100x PSG to 1 L DMEM and mix completely. Keep the complete cell culture media at 4°C and use within 1 month. This media is used for supporting cell propagation. CAUTION The complete cell culture media should be prepared in a biosafety cabinet to avoid contamination.

#### LB agar plates containing ampicillin or chloramphenicol

Add 4 g sodium chloride, 2 g yeast extract, 4 g tryptone and 6 g LB Agar to 400 mL deionized water in a 500 mL Wheaton glass bottle and autoclave at 121°C for 30 min. To prepare LB agar plates containing ampicillin or chloramphenicol, add 0.4 mL ampicillin stock (final working concentration: 150 mg/mL) or chloramphenicol stock (final working concentration: 12.5 mg/mL) to 400 mL LB agar solution after equilibrating the solution in a 55°C water bath for 30 min. In a bench area with a Bunsen burner on, pour ∼25 mL LB agar solution into each 100 mm Petri dish. Return the lids to the plates and leave the plates at room temperature overnight to solidify. The plates should be wrapped using plastic film, stored at 4°C, and used within 1 month. CAUTION Keep the chloramphenicol stock a certain distance away from the burner.

#### LB broth containing ampicillin or chloramphenicol

Dissolve 4 g sodium chloride, 2 g yeast extract and 4 g of tryptone into 400 mL deionized water in a 500 mL Wheaton glass bottle and autoclave at 121°C for 30 min. After cooling, LB broth could be stored at 4°C or room temperature. To prepare the LB broth containing ampicillin or chloramphenicol for bacteria growth, add 0.4 mL ampicillin stock or chloramphenicol stock to 400 mL LB medium, respectively in a biosafety cabinet. The final working concentration are 150 μg/mL for ampicillin and 12.5 μg/mL for chloramphenicol. Store the LB medium containing antibiotics at 4°C. CAUTION Use the prepared LB broth containing ampicillin or chloramphenicol within 1 month.

#### Sodium acetate solution (3.0 M, pH 5.2)

Dissolve 12.3 g sodium acetate into 30 mL nuclease-free water in a 250 mL Wheaton glass bottle. Adjust the pH to 5.2 using glacial acetic acid. Adjust the final volume to 50 mL using nuclease-free water. Store at room temperature.

#### Post-infection media

Add 22 mL FBS and 11 mL 100x PSG to 1 L DMEM and mix completely. Keep the post-infection media at 4°C and use it within 1 month. This media is used to maintain cells after being infected with a virus. CAUTION The post-infection media should be prepared in a biosafety cabinet to avoid contamination.

## PROCEDURES

### Section 1. Assembly of the full-length (FL) SARS-CoV-2 genome sequences into the empty pBeloBAC to generate pBeloBAC-FL

#### Preparation of the five fragments

1. Set up a 100 µL digestion reaction in a 0.2 mL PCR tube with appropriate restriction enzymes for each plasmid according to the information provided below. CAUTION F1 is released by a single-restriction digestion of BsmBI and generates compatible overhangs with those of PciI and HindIII.

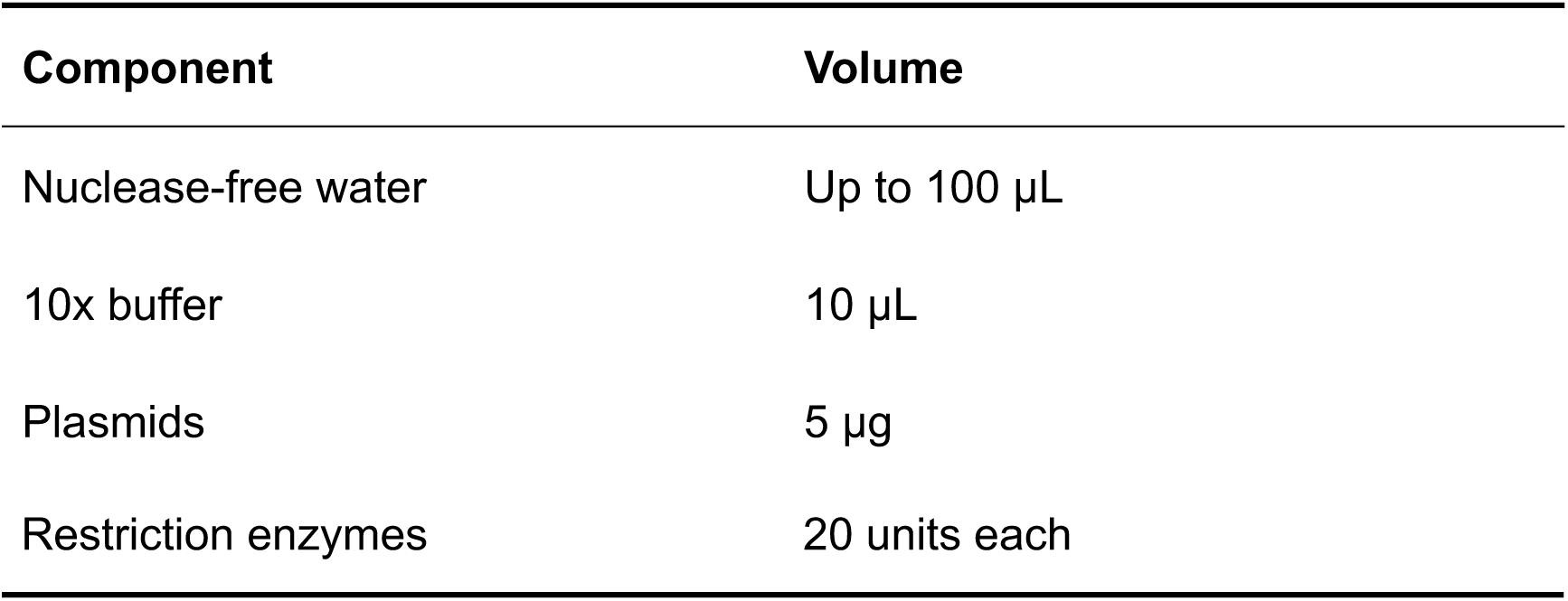

2. Incubate the mixture at 37°C in a thermal cycler for 3 h. CRITICAL BsmBI and BstBI work at 55°C and 65°C, respectively.

(i) Prepare five 0.6% agarose gels that are 10-cm long and with 10-mm-wide loading wells.

(ii) Weigh 0.6 g agarose powder and dispense into a conical 250 mL glass flask. Add 100 mL 1x TAE buffer.

(iii) After gently mixing, microwave for 3 min to completely melt the agarose.

(iv) Add 10 μL of 10,000x GelRed to the 0.6% agarose solution after the gel’s temperature has cooled down to 50°C. Mix thoroughly.

(v) Pour the mixture into a gel casting tray which has been fitted with a gel comb. CAUTION The agarose gel should be used after solidifying.

(vi) Place the solidified agarose gel into an electrophoresis tank and add enough 1x TAE buffer to ensure the gel is fully submerged.

2. Stop restriction digestion reaction by adding 20 µL of 6x Gel Loading Dye to each tube and mix thoroughly.

3. Load each DNA sample to 3 wells (40 µL/well) using a P100 pipette and load 10 µL of 1 kb plus DNA ladder in a separate well.

4. Separate the DNA fragments with electrophoresis at 60 V until bands are adequately separated.

5. Recover the DNA fragments from the gel using a gel extraction kit.

(i) Excise the target DNA bands in the gel by visualizing with a UV transilluminator and place each gel slice into a 2 mL tube. CAUTION Always wear a UV protection mask when the UV transilluminator is on.

(ii) Weigh the gel slice, add 3 volumes of Buffer QX1 and 2 volumes of nuclease-free water. CAUTION The weight of a 100 mg gel slice corresponds to a volume of 100 µL.

(iii) Add 30 µL of vortexed QIAEX II to each tube.

(iv) Incubate the tubes at 50°C for 10 min to solubilize the agarose gel slices and to bind the DNA. Mix by flicking and inverting the tube 5 times every 2 min until completely solubilized.

(v) Centrifuge the samples for 30 s and carefully remove supernatant with a P1000 pipette.

(vi) Add 500 µL Buffer QX1, resuspend the pellet by flicking and inverting the tube 5 times.

(vii) Centrifuge the sample for 30 s and remove the supernatant with a P1000 pipette.

(viii) Add 500 µL Buffer PE, resuspend the pellet by flicking and inverting the tube 5 times.

(ix) Centrifuge the sample for 30 s and remove the supernatant with a P1000 pipette.

(x) Add 500 µL Buffer PE, resuspend the pellet by flicking and inverting the tube 5 times.

(xi) Centrifuge the sample for 30 s, remove the supernatant with a P1000 pipette, and carefully remove all traces of supernatant with a P10 pipette.

(xii) Air-dry the sample until a white pellet is visualized. CAUTION It takes ∼20 min, it may lead to decreased elution efficiency if it is overdried.

(xiii) Add 30 µL nuclease-free water to resuspend the pellet by flicking 5 times. Incubate the tube at 50°C for 10 min.

(xiv) Centrifuge for 30 s, carefully pipet the supernatant into a clean 1.5 mL tube.

6. Measure the concentration using a NanoDrop, and run 1 μL of each fragment in a 0.6% agarose gel to check the size of DNA fragments. Image the gel using a ChemiDoc imaging system. The expected bands for F1-F5 are shown in **Fig. 2a**.

#### Preparation of the empty pBeloBAC

8. Electroporate the empty pBeloBAC plasmid into DH10B electrocompetent cells.

(i) Thaw a vial of DH10B electrocompetent cells on ice, dispense 40 µL to the tube containing 10 ng of the empty pBeloBAC DNA, and add deionized water to a final volume of 100 µL.

(ii) Transfer the 100 µL DH10B-DNA mixture into a pre-chilled 2 mm electroporation cuvette.

(iii) Pulse the DH10B-DNA mixture once with a MicroPulser electroporator using the EC2 program.

(iv) Immediately add 900 µL SOC medium to the electroporation cuvette, pipette up and down gently 3 times and move all the cells into a bacterial culture tube.

(v) Incubate the bacterial culture tube at 37°C with shaking at 250 rpm in an orbital shaker for 1 h.

(vi) Spread all bacteria (200 µL/plate) on pre-warmed LB plates containing chloramphenicol.

(vii) Incubate the plates inversely at 37°C for 16 h.

9. Pick a single colony into 200 mL LB broth that contains chloramphenicol in a 1,000 mL conical flask. Keep the flask at 37°C with 250 rpm shaking in an orbital shaker for 16 h.

10. Maxiprep the empty pBeloBAC.

(i) Pellet bacteria in a 250 mL bottle by centrifugating at 12,000 g for 5 min at 4°C.

(ii) Resuspend cells in 48 mL Solution I/RNase A.

(iii) Add 48 mL Solution II and invert gently to obtain a clear lysate.

(iv) Add 64 mL ice-cold N3 buffer and mix gently by inverting tube 10 times until flocculent white precipitate forms. Incubate on ice for 10 min.

(v) Spin down the precipitate by centrifugating at 15,000 g for 15 min at 4°C.

(vi) Clarify the supernatant by passing through a syringe filter.

(vii) Pass the clarified supernatant through a maxiprep column which is inserted in a vacuum manifold. CAUTION The maximum vacuum (negative) pressure is 150 mm Hg.

(viii) Add 10 mL HBC buffer to the column, switch on the vacuum source to draw the solution through the column, and then switch off the vacuum source.

(ix) Add 10 mL Wash buffer to the column, switch on the vacuum source to draw the solution through the column, and then switch off the vacuum source.

(x) Add another 10 mL Wash buffer to the column, switch on the vacuum source to draw the solution through the column, and then switch off the vacuum source.

(xi) Put the column back into a collection 50 ml tube and centrifugate at 12,000 g for 5 min at room temperature.

(xii) Elute DNA into a new 50 mL tube by adding 1 mL nuclease-free water into the column and centrifugating at 12,000 g for 5 min at room temperature.

(xiii) Using a NanoDrop, determine the concentration, which is usually ∼25 ng/µL.

11. Purify the empty pBeloBAC using an ATP-dependent DNase kit to remove the contaminated bacterial genome fragments. CRITICAL The pBeloBAC plasmids that are used for cloning must be purified.

(i) Set up a purification reaction in a 2.0 mL tube according to the table below.

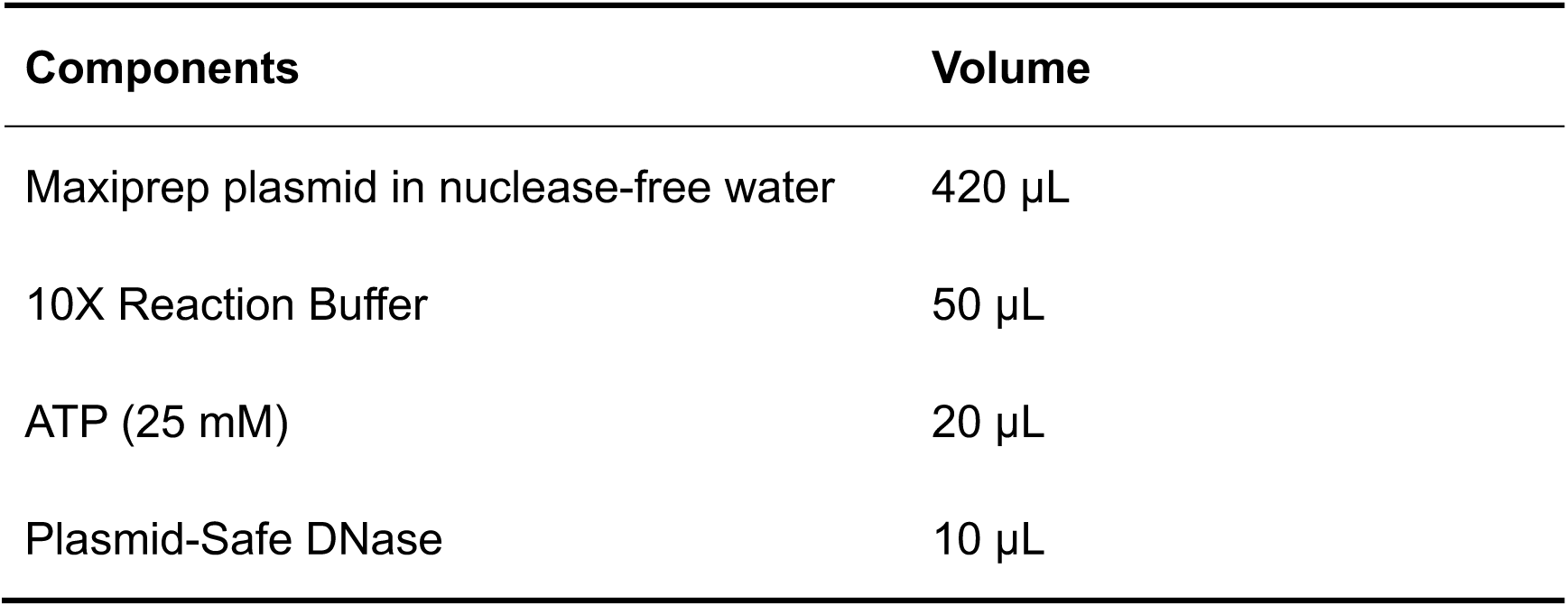

(ii) Incubate the tube at 37°C for 3 h.

(iii) Inactivate Plasmid-Safe DNase by incubating at 70°C for 30 min.

(iv) Add 50 µL of sodium acetate (3M, pH 5.2) and 1.25 mL of absolute ethanol to the reaction.

(v) After mixing thoroughly, freeze the solution at -80°C for 30 min.

(vi) Centrifugate at 15,000 g for 10 min at 4°C.

(vii) Decant supernatant, and resuspend the pellet in 1 mL of 75% ethanol.

(viii) Centrifugate at 15,000 g for 5 min at 4°C.

(ix) Discard supernatant and remove trace ethanol with a P100 pipette.

(x) Air-dry pellet for 5 min, dissolve the pellet into the desired volume of nuclease-free water, and measure concentration using a NanoDrop.

12. Aliquot the purified empty pBeloBAC and store at -80°C.

#### Preparation of the linearized empty pBeloBAC

13. Set up a digestion reaction of the purified empty pBeloBAC in a 0.2 mL tube according to the list below.

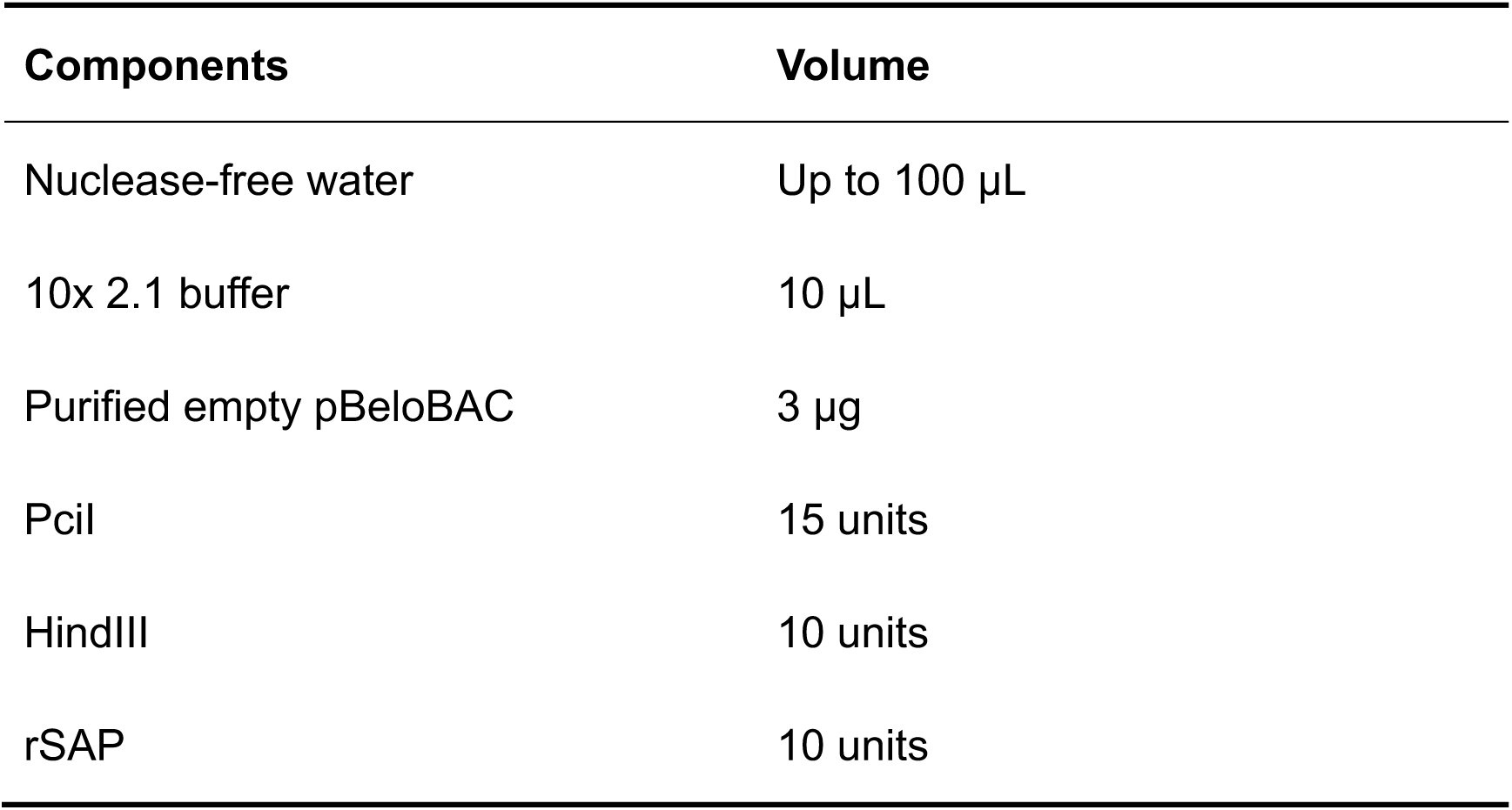

14. Incubate the mixture at 37°C for 3 h.

15. Prepare a 0.6% agarose gel as described in step 2.

16. Stop the digestion reaction by adding 20 µL of 6x Gel Loading Dye to each tube, mixing thoroughly.

17. Load the sample (40 µL/well) into a solidified agarose gel using a P100 pipette and load 10 µL of 1 kb plus DNA ladder in a separate well.

18. Separate the DNA fragments by electrophoresis at 60 V until bands are adequately separated.

19. Recover the linearized pBeloBAC from the gel as described in step 6.

20. After determining the concentration, aliquot the linearized empty pBeloBAC and store at -80°C.

#### Assembly of the F1 into the linearized pBeloBAC

21. Set up a ligation reaction in a 0.2 mL tube for assembling the F1 into the linearized empty pBeloBAC according to the table below.

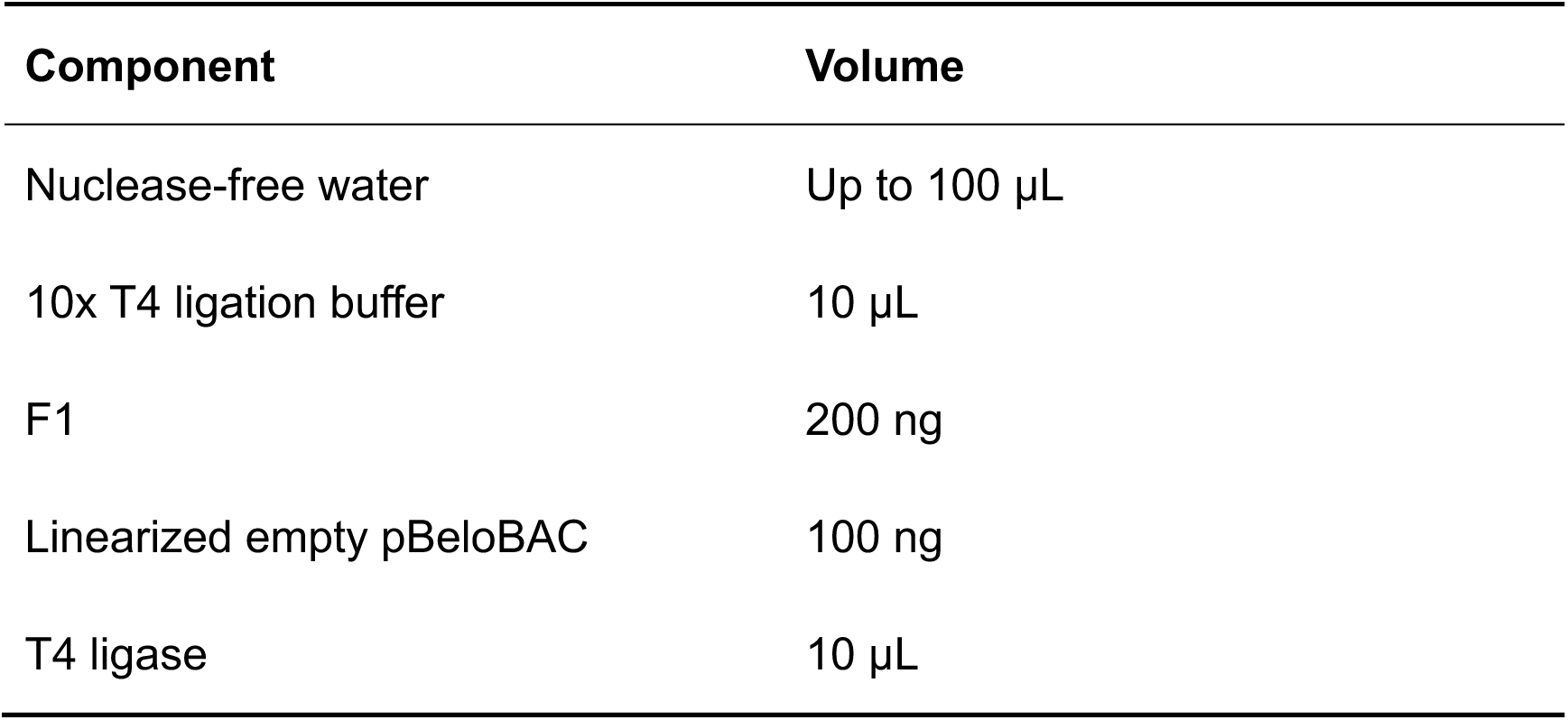

22. Incubate ligation at room temperature for 3 h.

23. Precipitate DNA from ligation reaction.

(i) Add 10 µL of sodium acetate solution (3.0 M, pH 5.2) and 250 µL of absolute ethanol to the tube, and incubate at -80°C for 30 min.

(ii) Centrifugate at 15,000 g for 10 min at 4°C.

(iii) Discard the supernatant, and resuspend the pellet with 1 mL of 75% ethanol.

(iv) Centrifugate at 15,000 g for 5 min at 4°C.

(v) Decant the supernatant, remove the trace fluid by using a P100 pipette and air-dry for 5 min.

(vi) Dissolve the pellet in 60 µL nuclease-free water and immerse the tube in ice.

24. Electroporate the ligated DNA into DH10B electrocompetent cells.

(i) Thaw a vial of DH10B electrocompetent cells, dispense 40 µL to the tube containing 60 µL of dissolved ligated DNA.

(ii) Transfer the 100 µL DH10B-DNA mixture into a pre-chilled 2 mm electroporation cuvette.

(iii) Pulse the DH10B-DNA mixture once with a MicroPulser electroporator using the EC2 program.

(iv) Immediately add 900 µL SOC medium to the electroporation cuvette, pipette up and down gently 3 times and move all the cells into a bacterial culture tube.

(v) Incubate the bacterial culture tube at 37°C with shaking at 250 rpm in an orbital shaker for 1 h.

(vi) Spread all bacteria on four pre-warmed LB plates containing chloramphenicol (250 µL/plate).

(vii) Incubate the plates inversely at 37°C for 16 h.

25. Pick 6 colonies and place each colony into a separate culture tube with 1 mL LB medium containing chloramphenicol.

26. Incubate the tubes at 37°C with shaking at 250 rpm in an orbital shaker for 3 h.

27. Colony PCR screen for positive clones using the primers of F1-F and F1-R (**Table 1**). An example result for colony PCR for cloning of F1 into pBeloBAC is shown in **Fig. 2b**. The PCR-positive clone is further confirmed by sanger sequencing using the CMV/F and bGH/R primers (**Table 2**) and designated as pBeloBAC-F1.

**Table 1.**
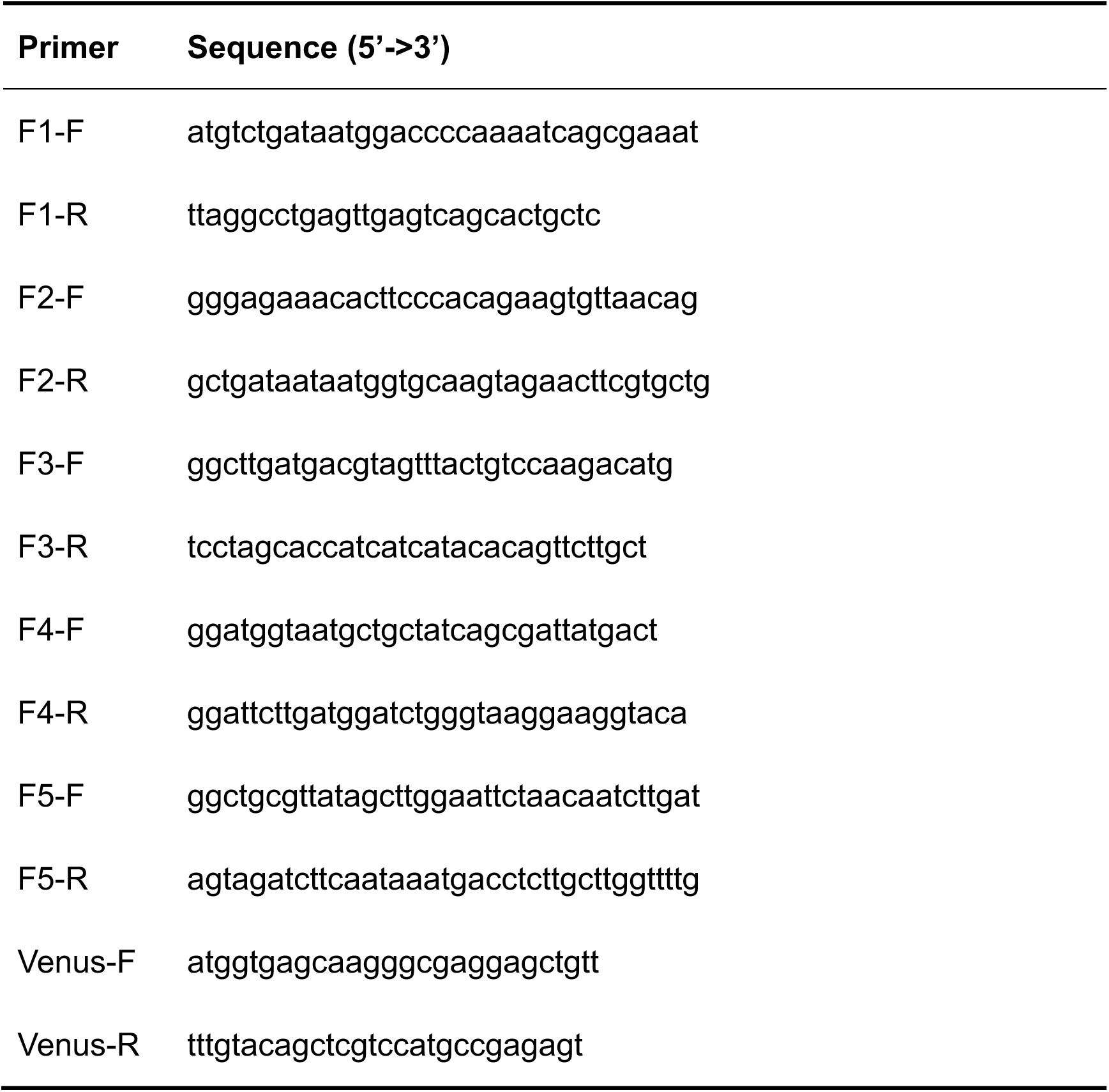
Primers and primer sequences for colony PCR screening

**Table 2.**
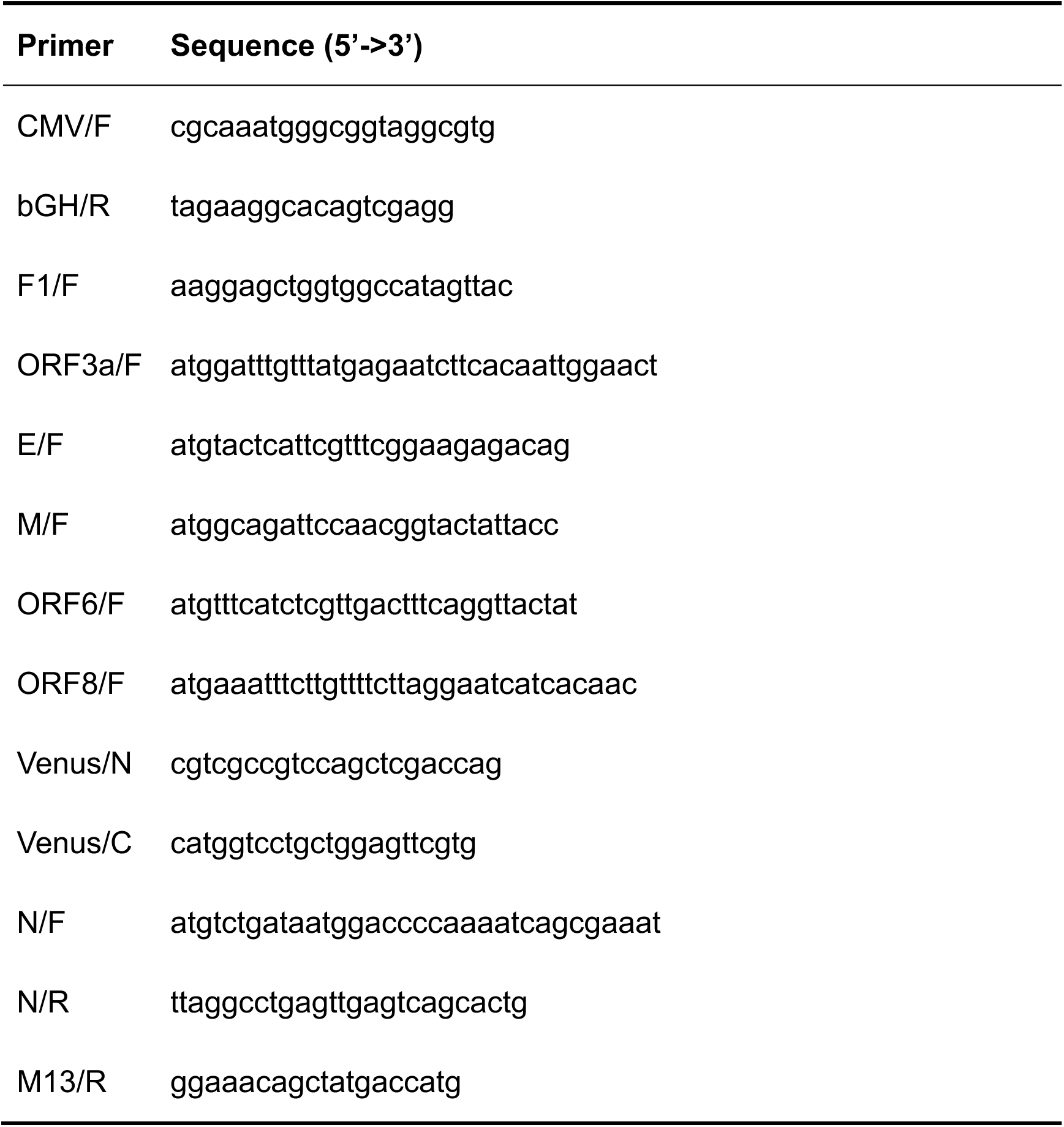
Primers and primer sequences for Sanger sequencing

#### Sequential assembly of the F2-F5 into the pBeloBAC-F1

28. Maxiprep and purify the pBeloBAC-F1 similar as described in steps 10 and 11.

29. Set up a restriction digestion of PacI and MluI to linearize the purified pBAC-F1, similar to step 13. Recover the linearized pBeloBAC-F1 from a 0.6% gel as described in step 6.

30. Assemble F3 into the linearized pBeloBAC-F1, similar to steps 21-26. Colony PCR screen for positive clones using the primers of F3-F and F3-R (**Table 1**).

31. Confirm insertion of F3 in PCR positive clones by digesting with PacI and MluI, then name the positive clone as pBeloBAC-F13, which contains F1 and F3. An example of the expected pBeloBAC-F13 digestion with PaI and MluI is shown in **Fig. 2c**.

32. Next, assemble F4 (using MluI and BstBI), F2 (using KasI and PacI), and F5 (using BstBI and BamHI) into pBeloBAC-F13 one by one as previously described for F3 into pBeloBAC-F1. Confirm by using both colony PCR and restriction digestion. Expected restriction analysis results of the sequential cloning of F4, F2, and F5 are shown in **Fig. 2d-f**. The resultant plasmids are named, respectively, pBeloBAC-F134, pBeloBAC-F1342, and finally pBeloBAC-FL, which contains F1, F2, F3, F4 and F5. CAUTION The insertion order is flexible, and we assemble them based on the order of the plasmids we receive from the company we used for chemical synthesis.

### Section 2. Rescue of rSARS-CoV-2 using the pBeloBAC-FL

#### Transfection of the pBeloBAC-FL into Vero E6 cells

33. Inoculate 200 µL of the DH10B that harbors pBeloBAC-FL into 200 mL LB broth containing chloramphenicol in a 1 L flask. Culture at 37°C with 250 rpm shaking in an orbital shaker for 16 h.

34. Maxiprep pBeloBAC-FL as described in step 10.

35. Transfect pBeloBAC-FL into Vero E6 cells in a 6-well plate. CAUTION Transfection to generate rSARS-CoV-2 using the pBeloBAC-FL must be performed in a biosafety level (BSL) 3 laboratory following institutional biosafety guidelines.

(i) Dilute 10 µL of Lipofectamine 2000 in 250 µL Opti-MEM in a 1.5 mL tube, and incubate at room temperature for 5 min.

(ii) Dilute 4 µg of pBeloBAC-FL in 250 µL Opti-MEM in a 1.5 mL tube. CAUTION We recommend transfecting 3 different clones of the plasmid to ensure a successful virus rescue.

(iii) Add the diluted Lipofectamine 2000 to the diluted pBeloBAC-FL.

(iv) Incubate the pBeloBAC-FL-Lipofectamine 2000 mixture at room temperature for 20-30 min.

(v) During the incubation, prepare a Vero E6 cell suspension and seed cells in a 6 well plate (∼5x10^5^ cells/well) in complete cell culture media after counting with Trypan blue staining. Add the pBeloBAC-FL-Lipofectamine 2000 mixture into the wells and gently shake back and forth 5 times.

(vi) Incubate the cell plate at 37°C in a CO_2_ incubator.

36. Replace the media with post-infection medium (4 mL/well) 24 h after transfection.

37. At 72 h post-transfection, collect the supernatant of transfected Vero E6 cells and dispense 1 mL/well to infect 2 different wells of fresh Vero E6 cells monolayers in a 6-well plate for confirmation of virus rescue using IFA and RT-PCR.

#### Confirmation of rSARS-CoV-2/WT rescue

38. At 48 h post-infection (hpi), check the infected cells using an EVOS imaging system for CPE. An example of expected CPE is shown in **Fig. 3b**.

39. Add 1 mL of TRIzol to one well of the infected cells for RNA extraction. Fix the other well of infected cells with 10% formalin solution for IFA. CAUTION The TRIzol-treated sample and the 10% formalin-fixed plate could now be moved to BSL2 by complying with proper biosafety guidelines and institutional policy.

40. Analyze the 10% formalin-fixed cells by IFA.

(i) Permeabilize the cells using 0.5% Triton X-100 (2 mL/well) for 10 min at room temperature.

(ii) Block the cells using blocking solution (1 mL/well) for 30 min at 37°C.

(iii) Dilute the primary antibody against viral N protein (1C7C7) in blocking solution at a final concentration of 1 µg/mL, and incubate with the cells (1 mL/well) at 37°C for 1 h.

(iv) Wash the cells 3 times with PBS.

(v) Dilute the TRITC-labeled donkey anti-mouse IgG secondary antibody in blocking solution to a final concentration of 1 µg/mL, and incubate with the cells (1 mL/well) at 37°C for 1 h.

(vi) Wash the cells 3 times with PBS.

(vii) Dilute the DAPI solution in PBS to a final concentration of 1 µg/mL, and incubate with the cells (1 mL/well) at 37°C for 10 min.

(viii) Wash the cells 3 times with PBS.

(ix) Observe and photograph the cells using an EVOS imaging system. An example of an IFA result is shown in **Fig. 3c**.

41. During the immunostaining, extract RNA from the TRIzol-treated cells.

(i) Add 200 µL of chloroform into 1 mL of TRIzol-treated sample and mix thoroughly by vigorous shaking. CAUTION Do not vortex.

(ii) Centrifuge the sample for 15 min at 15,000 g at 4°C. The mixture should separate into a lower red phenol-chloroform, interphase, and a colorless upper aqueous phase.

(iii) Collect the aqueous phase containing the RNA into a new 1.5 mL tube.

(iv) Add 500 µL of isopropanol, mix thoroughly by inverting the tube 10 times, and incubate for 5 min at room temperature.

(v) Centrifuge for 10 min at 15,000 g at 4°C.

(vi) The total RNA precipitate should form a white gel-like pellet at the bottom of the tube.

(vii) Decant the supernatant, and resuspend the pellet in 1 mL of 75% ethanol. (viii)Centrifuge at 15,000 g for 5 min at 4°C.

(viii) Decant the supernatant, and remove trace ethanol using a P100 pipette.

(ix) Air-dry the RNA pellet for 5 min at room temperature. CRITICAL Overdried RNA is very difficult to dissolve.

(x) Resuspend the pellet in 100 µL nuclease-free water and determine the concentration using a NanoDrop. CAUTION Proceed to downstream applications or store the RNA at -80°C for less than 1 month.

42. Set up a RT reaction to convert RNA to cDNA.

(i) Combine the following components in a 0.2 ml tube.

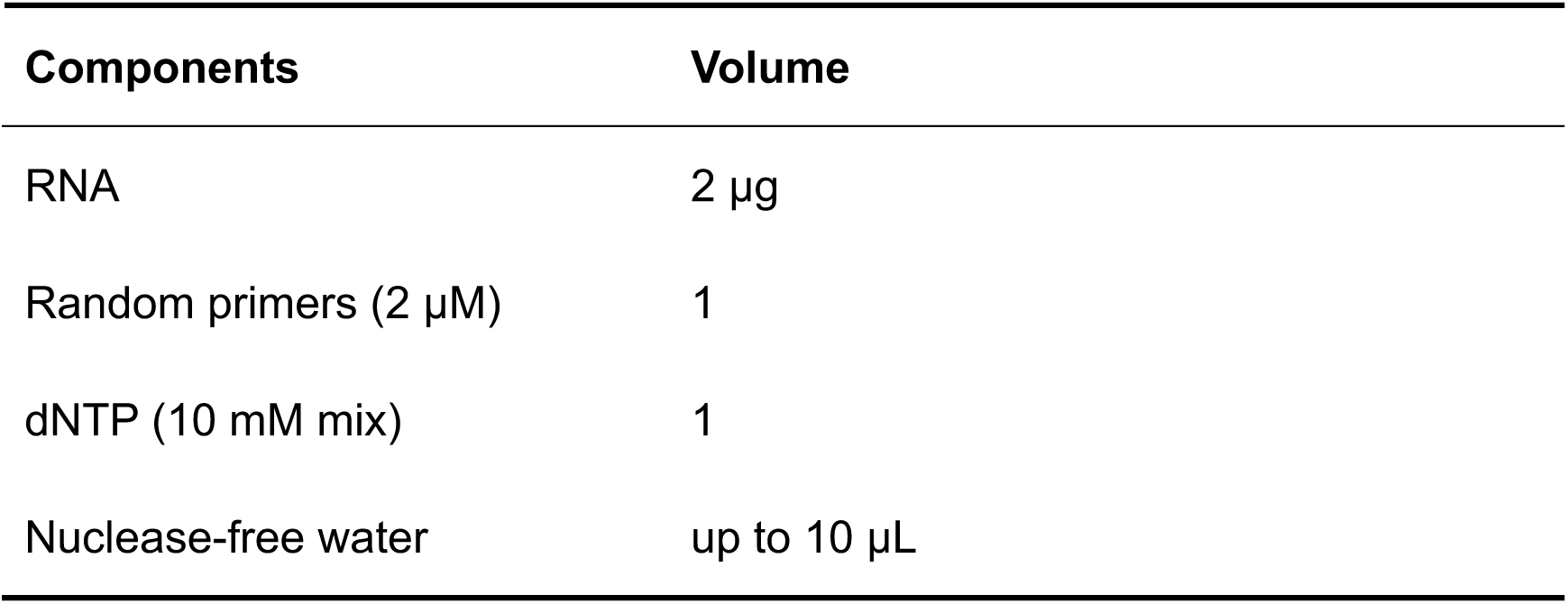

(ii) Incubate the mixture at 65°C for 5 min, then immediately place on ice for 1 min.

(iii) Add the following components to each RNA/primer mixture, mix gently, and collect by brief centrifugation.

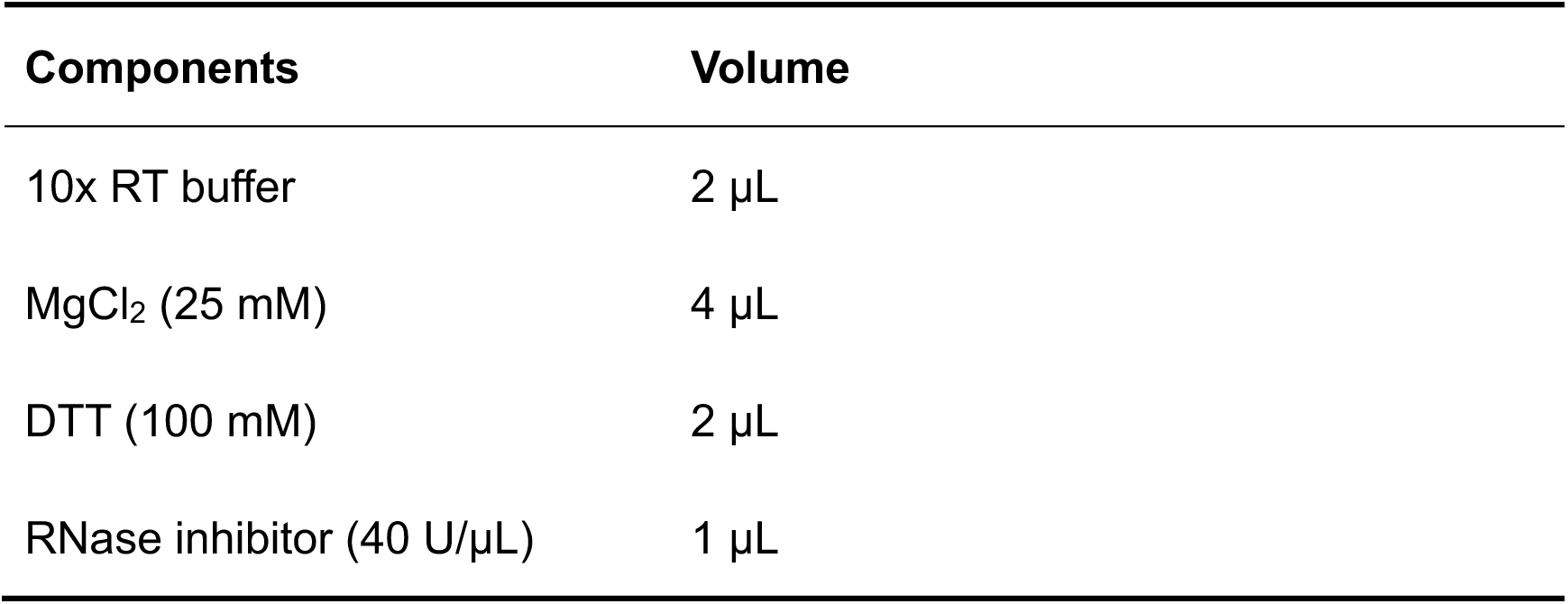

(iv) Incubate at 25°C for 2 min.

(v) Add 1 µl of the SuperScript II reverse transcriptase to each tube and incubate at 42°C for 50 min.

(xii) Terminate the reaction by incubating at 70°C for 15 min. CAUTION Proceed to downstream applications or store the cDNA at -20°C.

43. Set up a PCR reaction in a 0.2 mL tube according to the table below. The sequences of the primers are listed in **Table 3**.

**Table 3.**
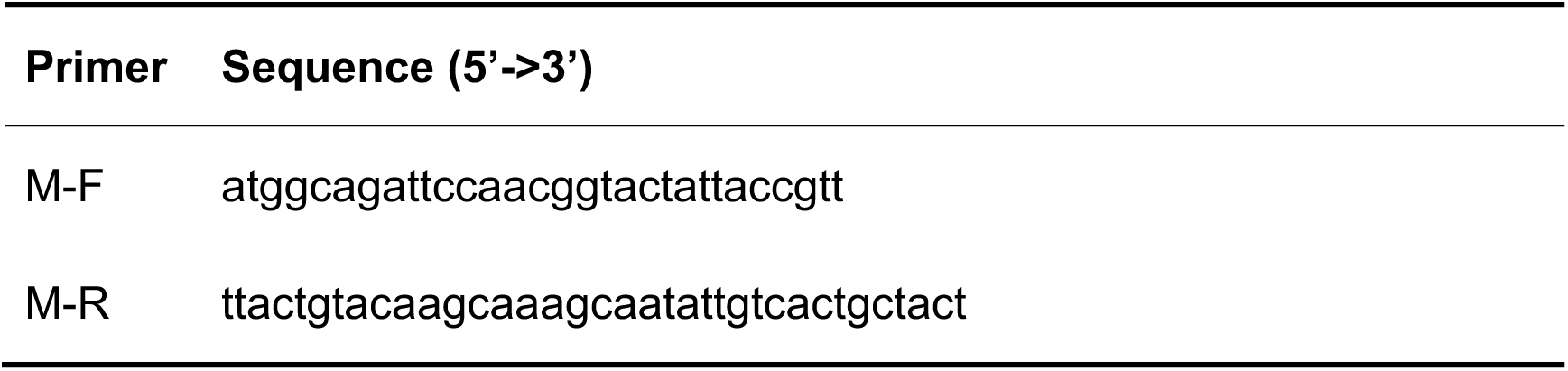
Primers and primer sequences for RT-PCR

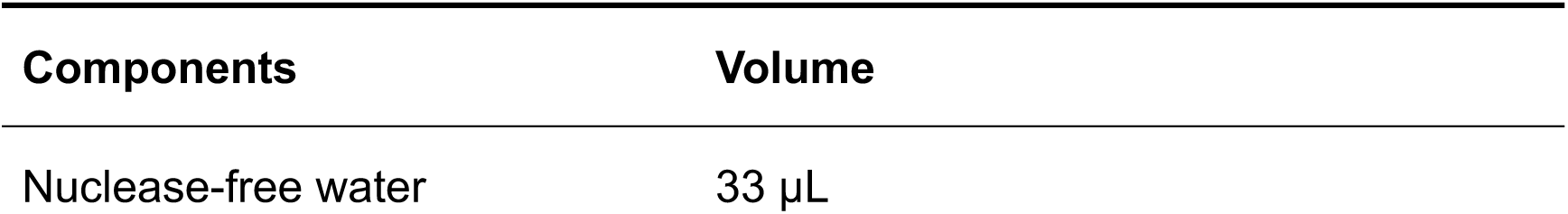
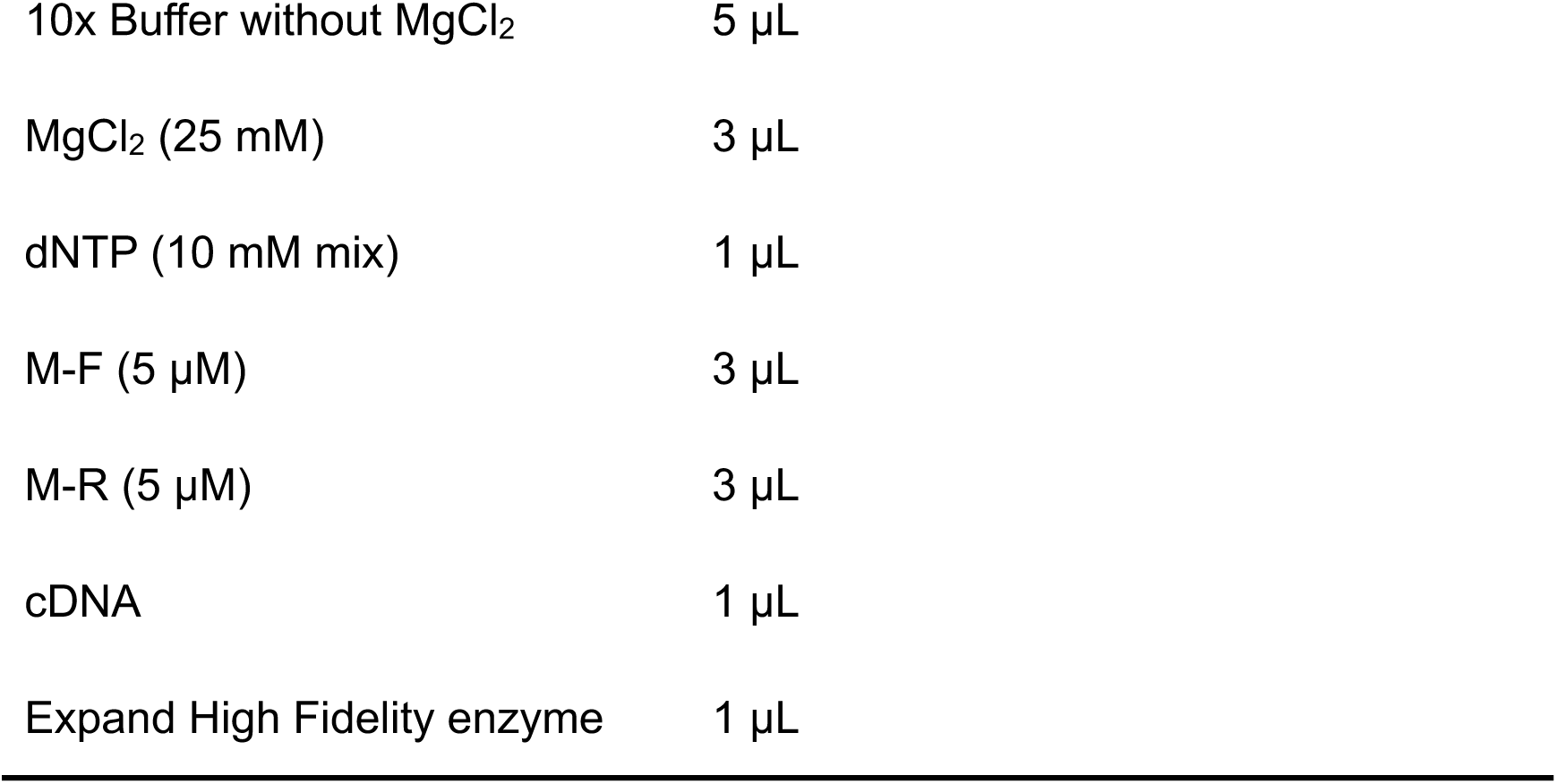

44. Place the tube in a thermal cycler and start the cycling using the conditions listed below.

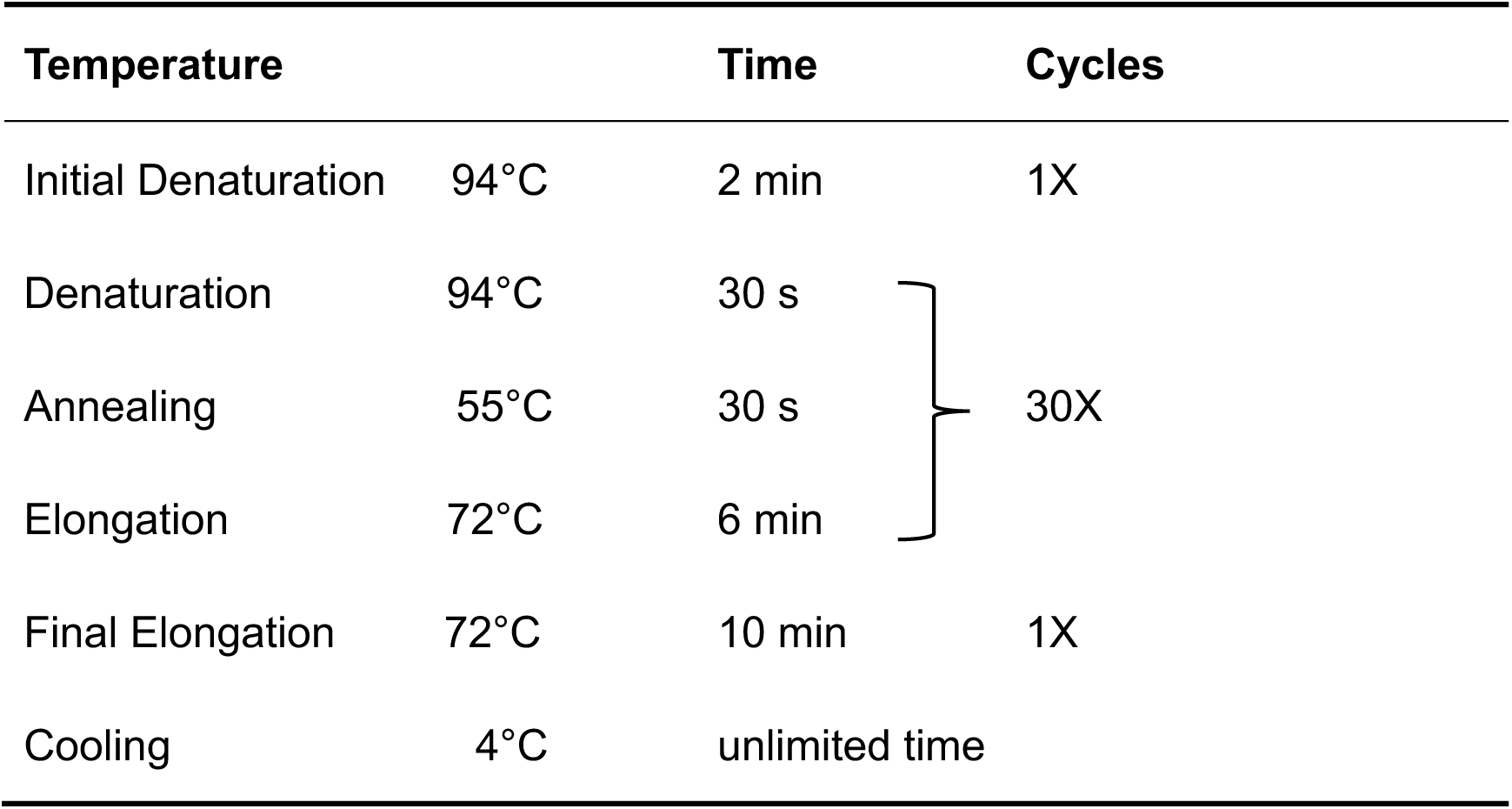

45. Confirm PCR amplification in a 0.6% agarose gel and recover target DNA band from the gel as described in step 6.

46. Sequence PCR product using primer of M/F (**Table 2**). The expected results of sequencing the M gene from rSARS-CoV-2/WT are shown in **Fig. 3d**.

### Section 3. Cloning of the Venus reporter gene into the pBeloBAC-FL

#### Cloning of Venus into the pUC57-F1

47. Set up a PCR reaction in a 0.2 mL tube according to the list below to amplify Venus-2A. The sequences of the primers are listed in **Table 4**.

**Table 4.**
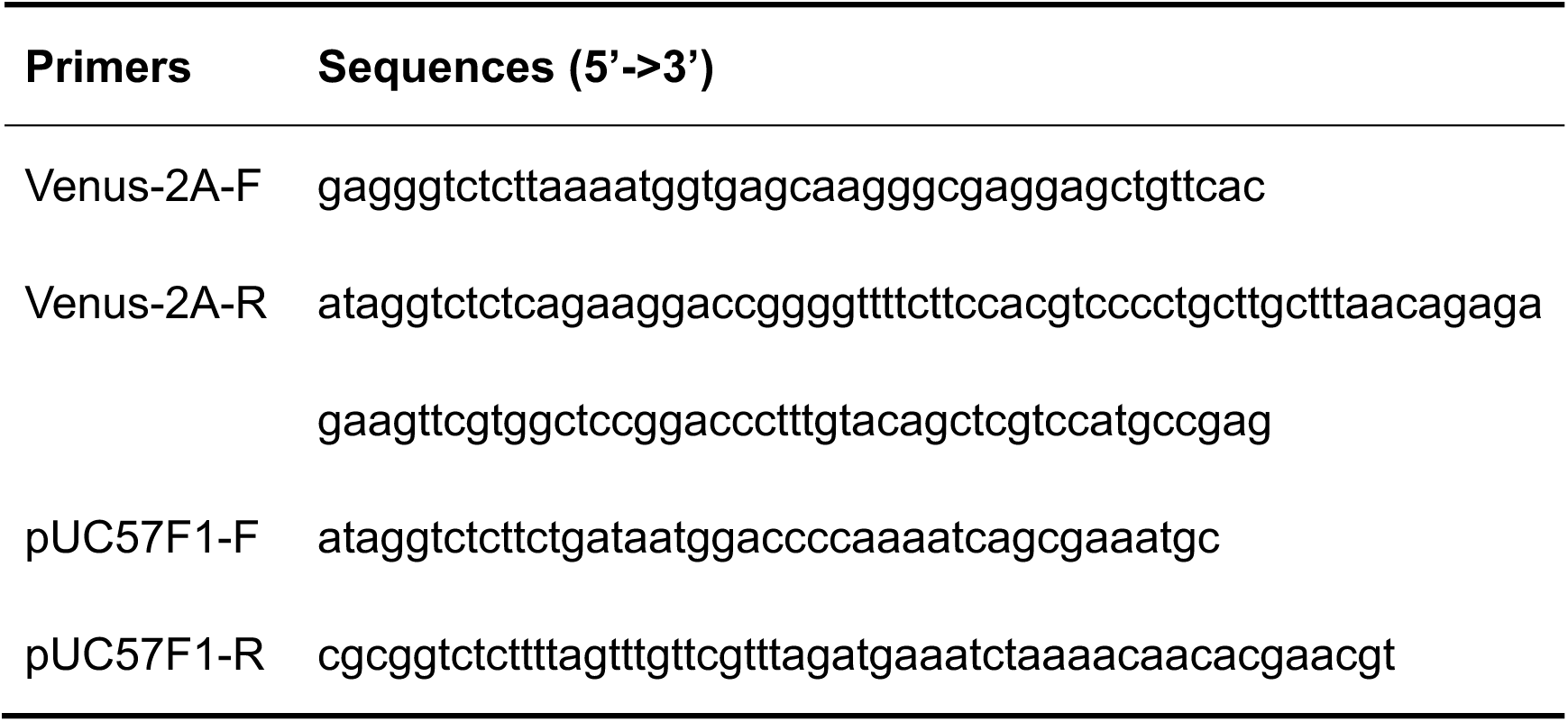
Primers and primer sequences for generation of rSARS-CoV-2/Venus-2A

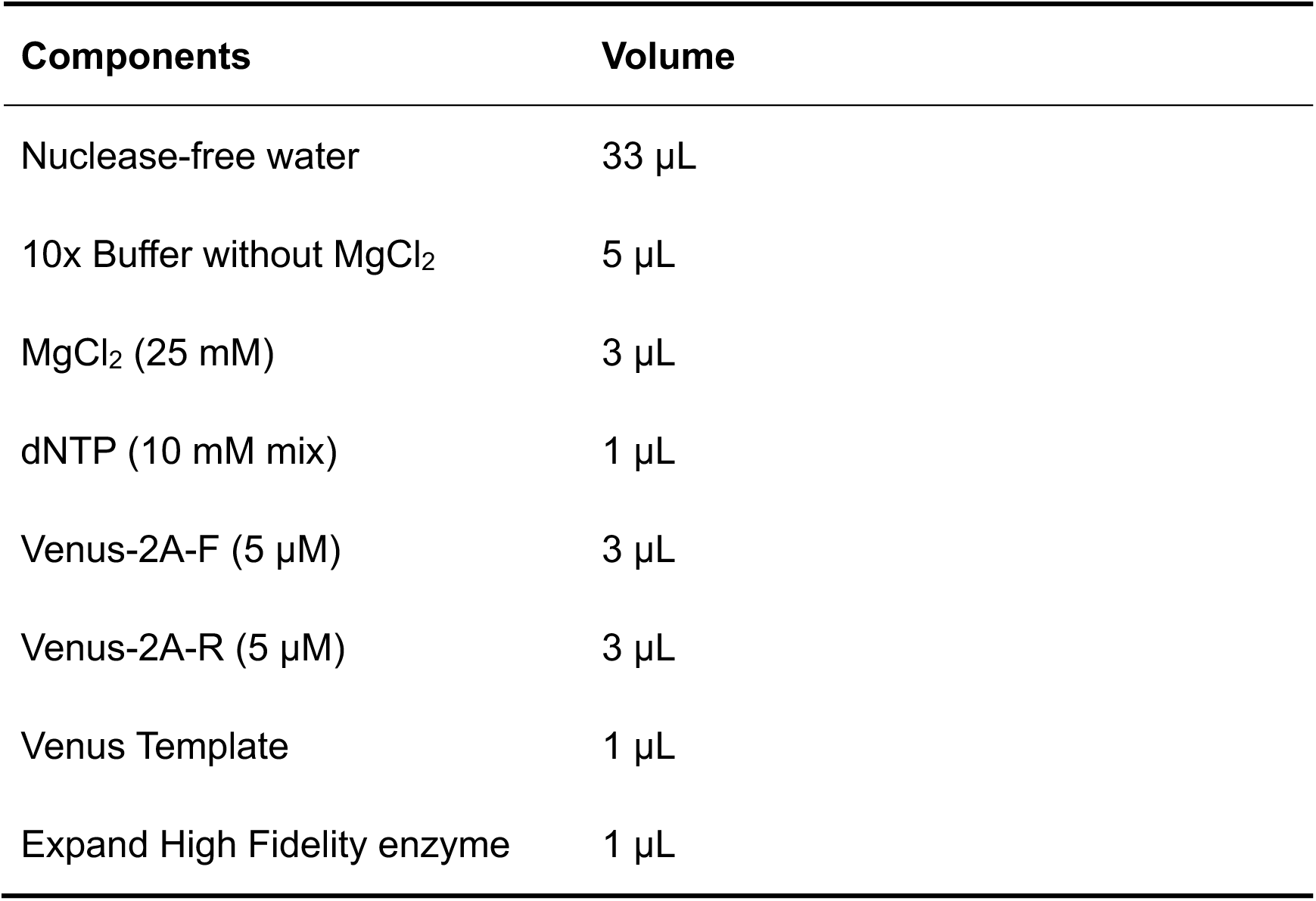

48. Place the tube in a thermal block cycler and start the cycling using the thermal profile as described in step 44.

49. Recover the amplified Venus-2A from a gel as described in step 6.

50. Set up a PCR reaction in a 0.2 mL tube according to the list below to amplify pUC57F1. The sequences of the primers are listed in **Table 4**.

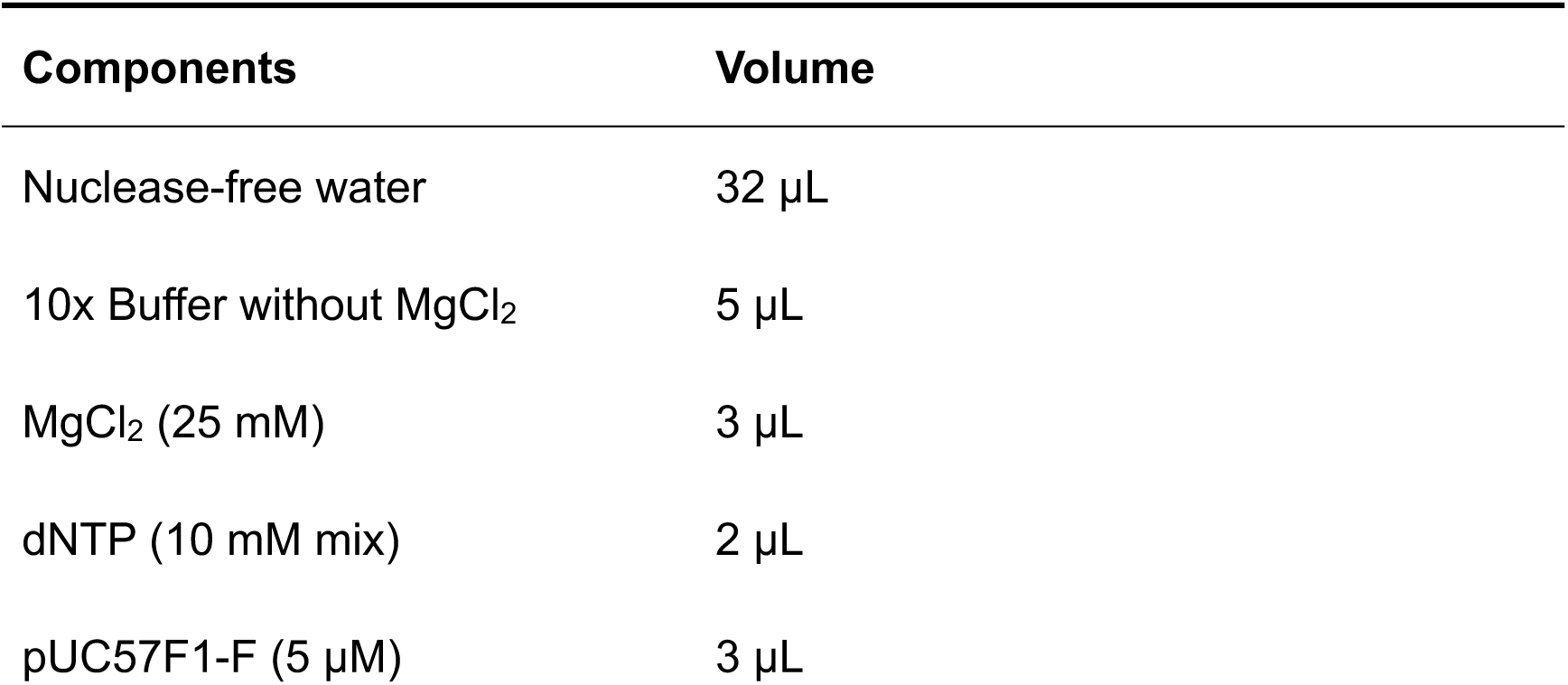
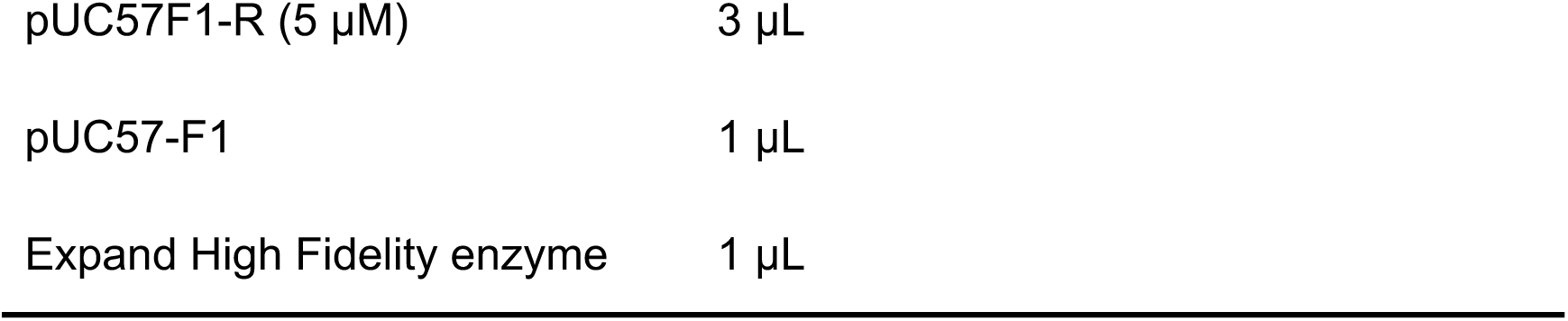

51. Place the tube in a thermal block cycler and start the cycling using the thermal profile below as described in step 44.

52. Recover the amplified pUC57F1 from a gel as described in step 6.

53. Run 1 μL of each DNA product in a 0.6% agarose gel to check the size. Image the gel using a ChemiDoc imaging system. The expected bands for the PCR products are shown in **Fig. 5a**.

54. Set up digestion reactions for the purified PCR products of Venus-2A and pUC57F1 according to the table below.

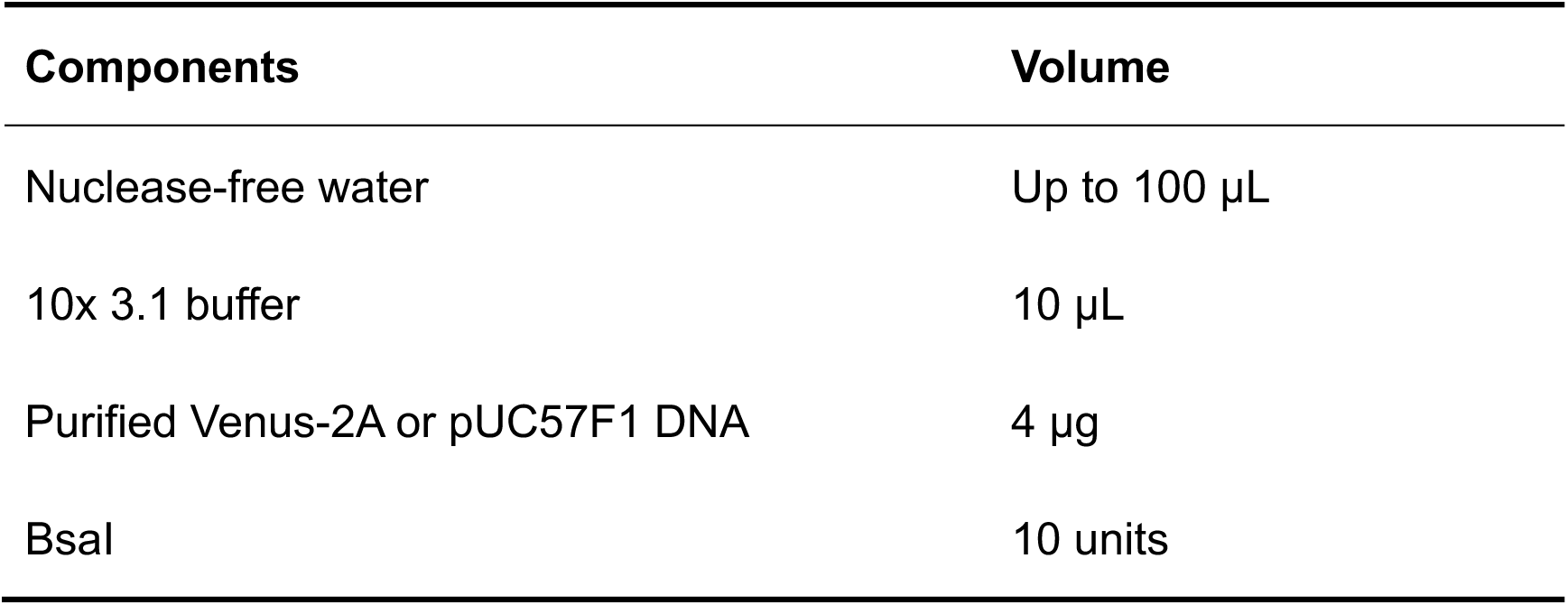

55. Incubate at 37°C for 3 h.

56. Recover the DNA using a PCR clean-up kit. CAUTION There are 2 bands after BsaI digestion of the amplified pUC57F1, cut and combine these 2 bands into one tube if using gel purification procedure.

(i) Add 3 volumes membrane binding solution of PCR reaction, mix thoroughly.

(ii) Pass the mixture through a column by centrifugating at 15,000 g for 10 s.

(iii) Wash the column twice with 75% ethanol solution.

(iv) Centrifugate at 15,000 g for 1 min to remove trace ethanol.

(v) Elute DNA by adding 30 µL nuclease-free water.

57. Set up a ligation reaction according to the table below to assemble Venus-2A into pUC57F1.

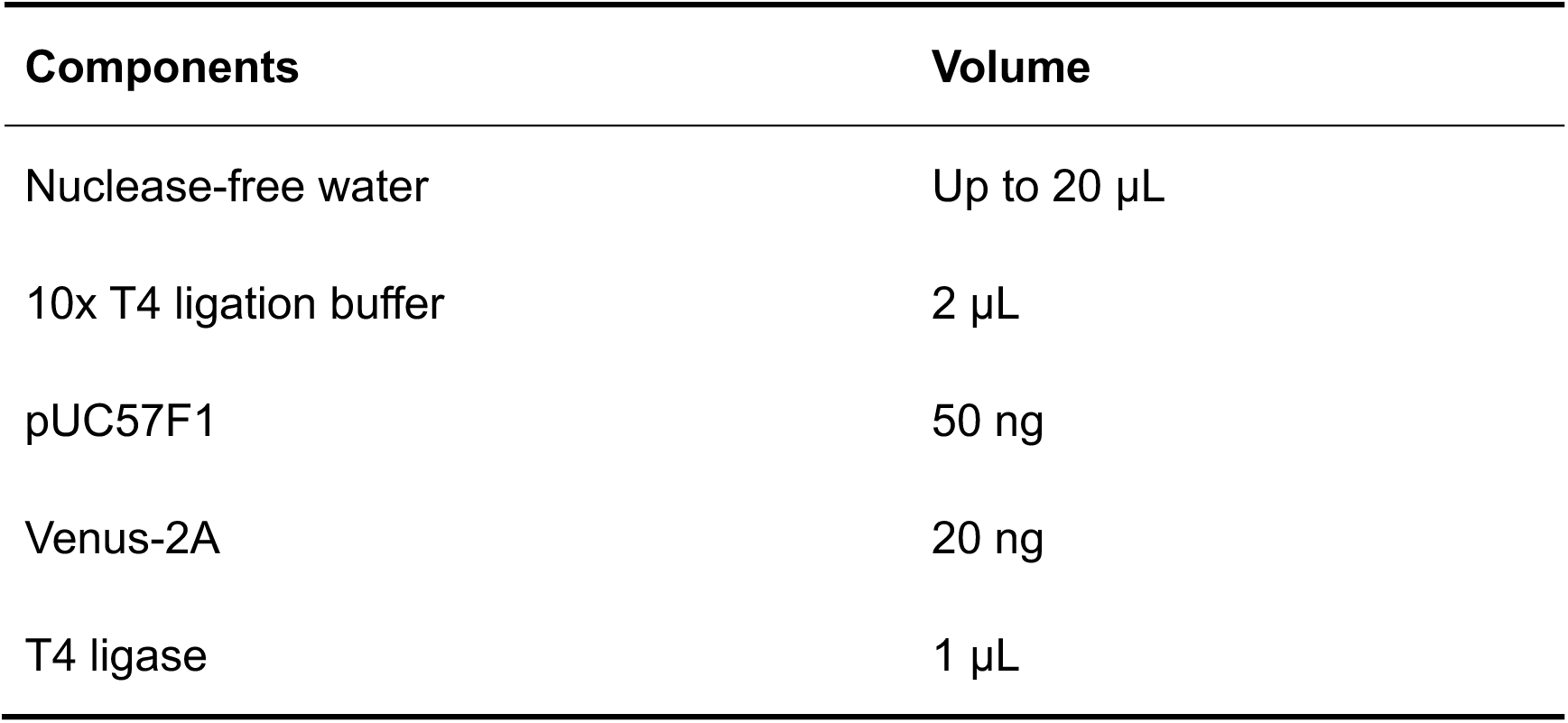

58. Incubate the ligation reaction at room temperature for 1 h.

59. Transform the ligated DNA into NEB Stable Competent cells.

(i) Thaw 1 tube of NEB Stable Competent cells on ice.

(ii) Add 10 µL of ligation solution to the competent cells, and carefully flick the tube 3 times to mix cells and DNA.

(iii) Place the mixture on ice for 30 min.

(iv) Heat shock the cells in the mixture at exactly 42°C for exactly 30 seconds.

(v) Immediately put the tubes back on ice for 5 min.

(vi) Pipette 950 µL of room temperature SOC medium into the mixture and place the mixture at 37°C with shaking at 250 rpm in an orbital shaker for 60 min.

(vii) Spread 100 µL of cells onto prewarmed LB agar plates containing ampicillin.

(viii) Incubate plates invertedly at 37°C for 16-24 h.

60. Pick three colonies and place each colony in 5 mL of LB broth containing ampicillin in a bacterial culture tube and culture at 37°C with shaking at 250 rpm in an orbital shaker for 16-24 h.

61. Miniprep plasmids using an Omega miniprep kit.

(i) Pellet the bacterial culture in a 2.0 mL tube at 15,000 g at 4°C for 1 min.

(ii) Resuspend cells in 500 µL Solution I/RNase A.

(iii) Add 500 µL Solution II and invert gently to obtain a clear lysate.

(iv) Add 700 µL ice-cold N3 buffer and mix gently by inverting tube 10 times until a white precipitate forms, and incubate the tube on ice for 10 min.

(v) Spin down the precipitate by centrifugating at 15,000 g at 4°C for 10 min.

(vi) Pass the clarified supernatant through a miniprep column which is inserted in a vacuum manifold. CAUTION The maximum vacuum (negative) pressure is 150 mm Hg.

(vii) Add 500 µL HBC buffer to the column, switch on the vacuum source to draw the solution through the column, and then switch off the vacuum source.

(xiv) Add 700 µL Wash buffer to the column, switch on the vacuum source to draw the solution through the column, and then switch off the vacuum source.

(xv) Add another 700 µL Wash buffer to the column, switch on the vacuum source to draw the solution through the column, and then switch off the vacuum source.

(viii) Put the column back into a collection tube and centrifugate at 15,000 g at room temperature for 5 min.

(ix) Elute DNA into a new 1.5 mL tube by adding 100 µL nuclease-free water to the column and centrifugate at 15,000 g at room temperature for 5 min. Using a NanoDrop, determine the concentrations of the plasmids, which are usually 100-200 ng/µL.

62. Sequence the entire plasmid using the primers listed in **Table 2**. The positive plasmid is designated as pUC57-F1/Venus-2A.

#### Cloning of Venus into the pBeloBAC-FL

63. Set up a digestion reaction for pUC57-F1/Venus-2A according to the recipe listed below.

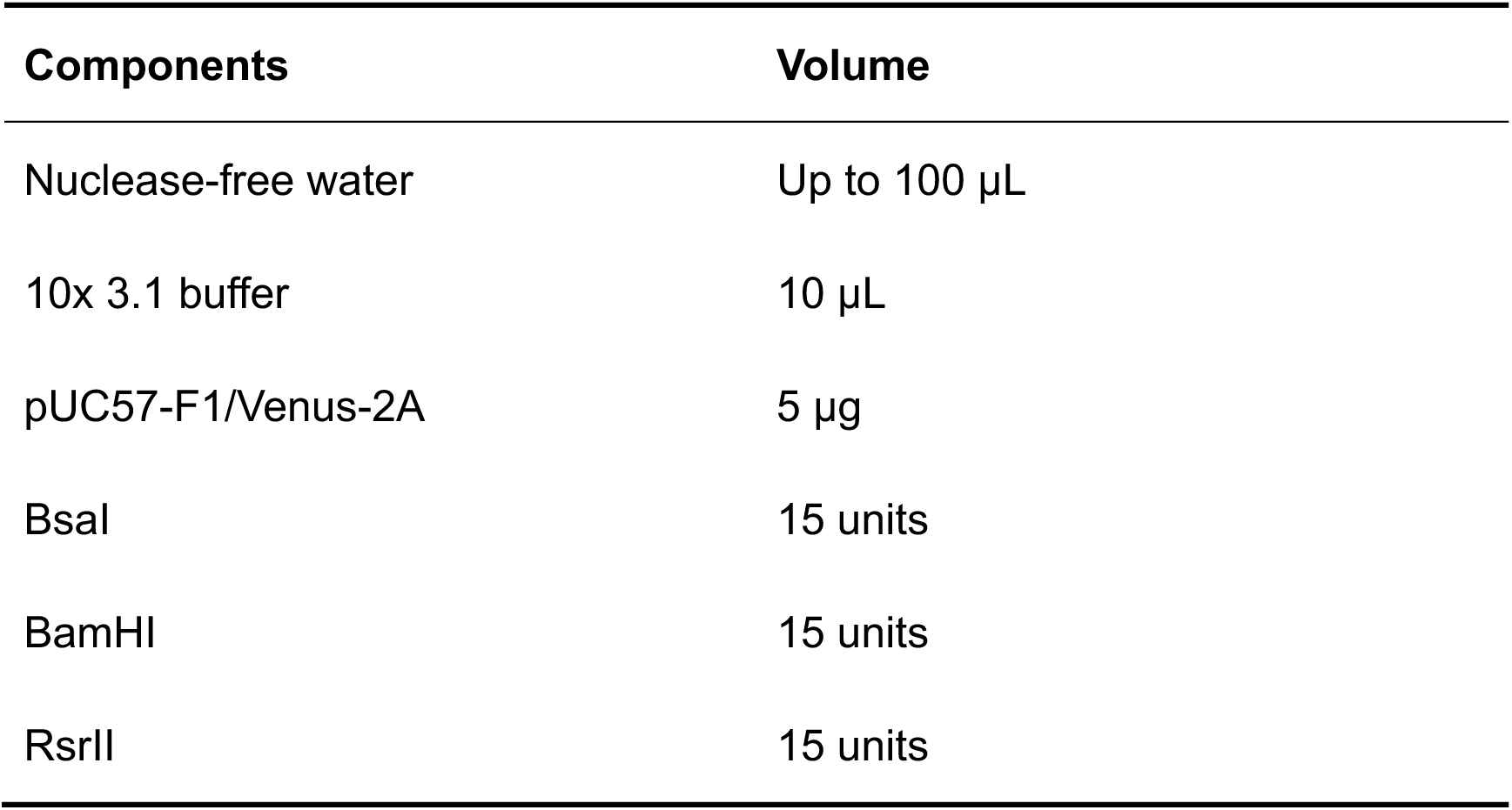

64. Incubate at 37°C for 3 h.

65. Separate the digested DNA on a 0.6% agarose by electrophoresis at 60 V until bands are adequately separated (**Fig. 5b, left lane**) and recover the top DNA (F1/Venus-2A) as described in step 6.

66. Elute target DNA in 30 µL nuclease-free water and measure concentration using a NanoDrop.

67. Maxiprep the pBeloBAC-FL as described in step 10.

68. Purify the pBeloBAC-FL as described in step 11.

69. Set up a digestion reaction in a 0.2 mL tube to linearize the purified pBeloBAC-FL according to the recipe listed below.

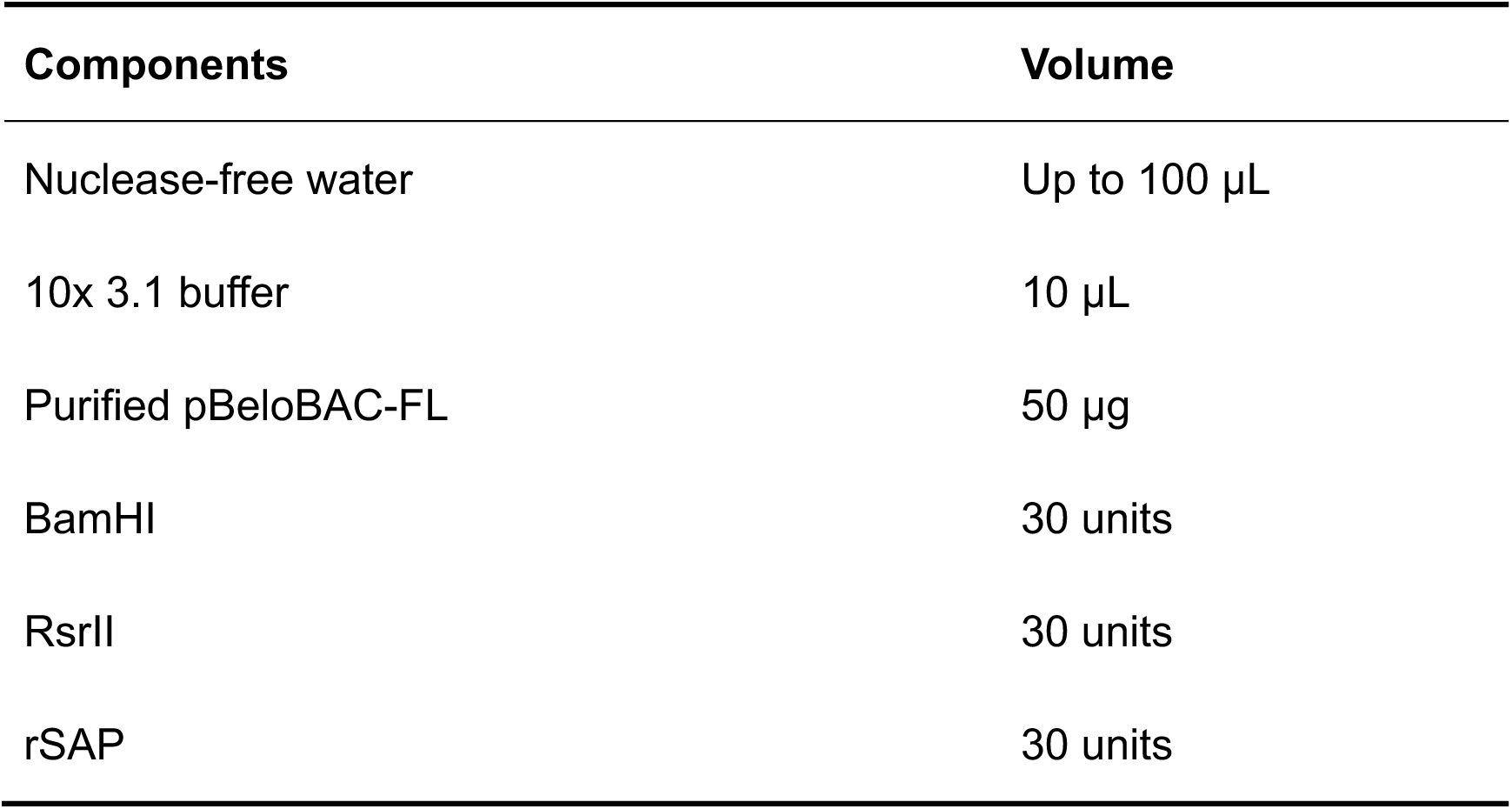

70. Incubate the reaction at 37°C for 3 h.

71. Stop the reaction by adding 20 µL of 6x Gel Loading Dye and mix thoroughly by pipetting gently up and down 3 times.

72. Load the mixture on a 0.6% agarose gel, separate the bands by electrophoresis at 60 V until bands are adequately separated (**Fig. 5b, right lane**). CRITICAL Do not load more than 30 µL per well, otherwise the linearized pBeloBAC-FL band will stack in the well.

73. Recover the pBeloBAC-FL backbone band as described in step 6.

74. Elute the linearized pBeloBAC-FL in 100 µL nuclease-free water and measure the concentration using a NanoDrop.

75. Aliquot the linearized pBeloBAC-FL and store at -80°C.

76. Set up a ligation reaction for cloning F1/Venus-2A into the digested pBeloBAC-FL in a 1.5 mL tube according to the recipe listed below.

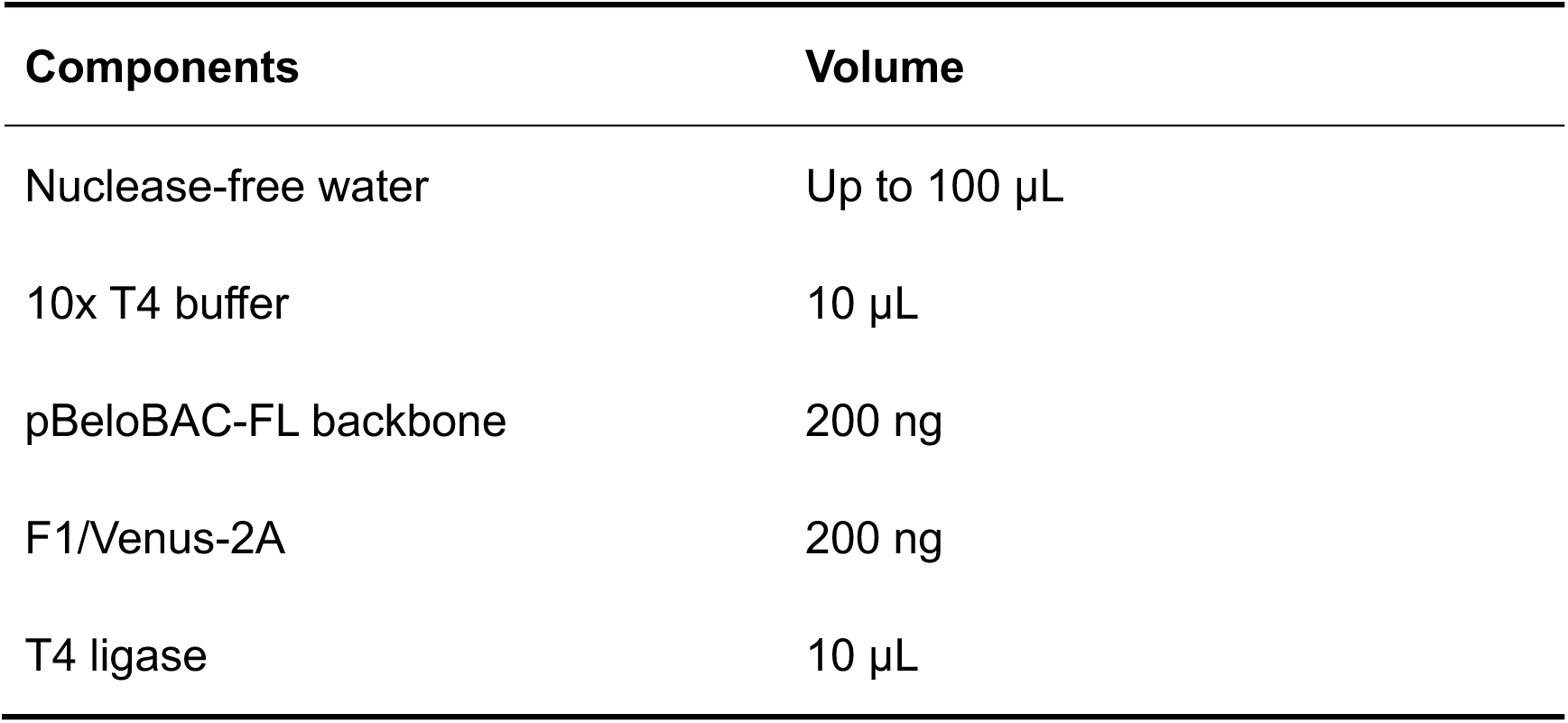

77. Incubate at room temperature for 6 h.

78. Recover the ligated DNA as described in step 23.

79. Electroporate the ligated DNA into DH10B cells as described in step 25.

80. Pick 6 colonies and place each in a bacteria culture tube that contains 1 mL LB broth containing chloramphenicol.

81. Culture at 37°C for 3 h.

82. Colony PCR screen for positive clones as described above using the primers binding to F5 (F5-F and F5-R) and Venus (Venus-F and Venus-R) (**Table 1**). The positive clone is named as pBeloBAC-FL/Venus-2A. An example of PCR screening of positive pBeloBAC-FL/Venus-2A clones is shown in **Fig. 5c**.

### Section 4. Recovery of the Venus-expressing rSARS-CoV-2 (rSARS-CoV-2/Venus-2A)

#### Rescue of rSARS-CoV-2/Venus-2A

83. Inoculate 100 µL of PCR-positive bacteria into 100 mL LB broth containing chloramphenicol in a 1 L flask. Culture at 37°C with shaking at 250 rpm in an orbital shaker for 16 h. CRITICAL We usually select 3 PCR-positive clones to expand.

84. Maxiprep the pBeloBAC-FL/Venus-2A as described in step 10.

85. Transfect Vero E6 cells with the purified pBeloBAC-FL/Venus-2A as described in step 35 for pBeloBAC-FL.

86. Replace the media with post-infection medium (4 mL/well) at 24 h post-transfection.

87. Return the plates to the 37°C CO_2_ incubator.

#### Characterization of the rSARS-CoV-2/Venus-2A

88. Check the transfected cells under an EVOS imaging system for fluorescence Venus expression and collect the supernatant at 72 h post-transfection.

89. Infect fresh confluent Vero E6 cells in a 6-well plate by adding 1 mL/well of the supernatant of the transfected cells for 72 h.

90. Analyze the supernatant of the transfected cells by plaque assay.

(i) Dilute the supernatant in 10-fold serial using post-infection medium.

(ii) Infect monolayers of Vero E6 cells in a 6-well plate with the diluted supernatant for 1 h at 37°C.

(iii) During viral infection, prepare the semi-solid medium according to the list below. CRITICAL Melt the agar completely in a microwave oven and mix thoroughly before adding to the medium.

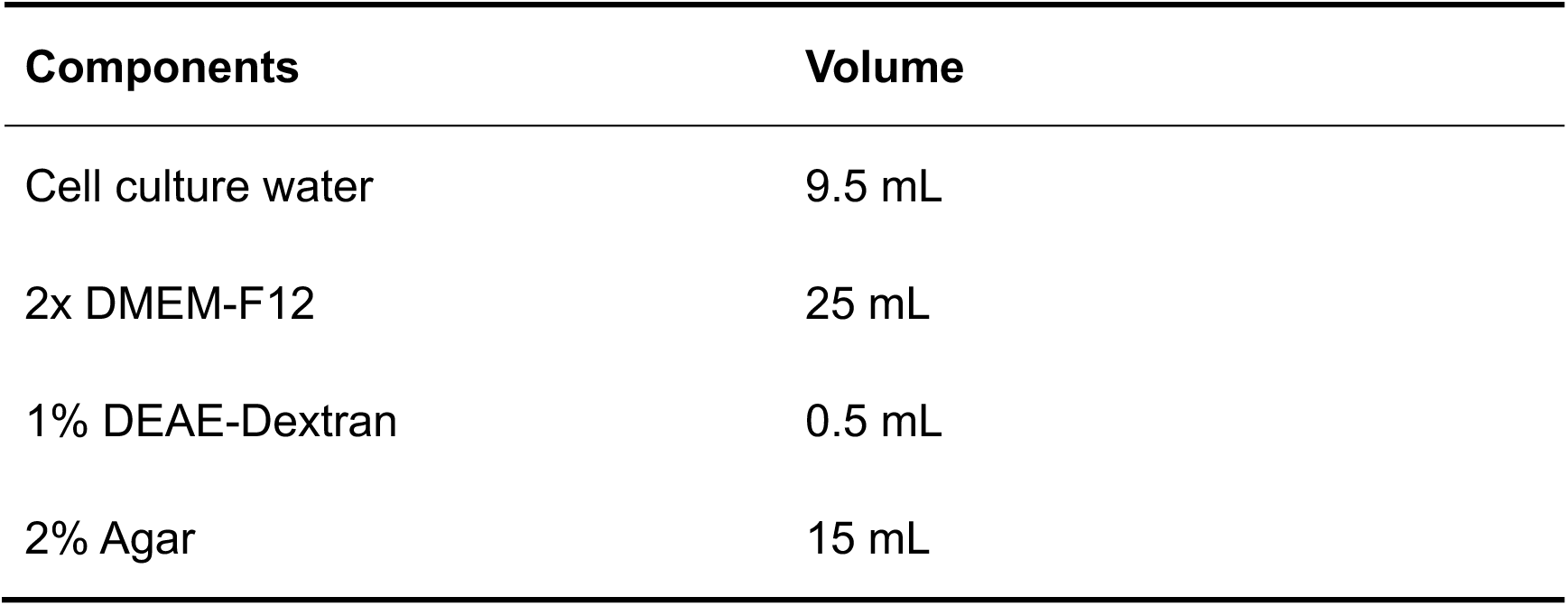

(iv) Discard the infection supernatant and wash cells 3 times with PBS.

(v) Overlay the cells with 4 mL of semi-solid medium. CRITICAL Ensure the temperature of the semi-solid medium does not exceed 42°C before overlaying the cells.

(vi) Solidify the medium by leaving the plates at room temperature for 10 min.

(vii) Incubate the plates invertedly at 37°C for 72 h in a 5% CO_2_ incubator.

91. Fix the plates in 10% formalin. The 10% formalin-fixed plates could now be moved to BSL2 by complying with institutional biosafety policy.

92. Analyze the fresh infected cells by IFA as described in step 39. The expected results of Vero E6 cells infected with rSARS-CoV-2/Venus-2A stained with the MAb 1C7C7 against the viral N protein are shown in **Fig. 6a**.

93. Remove the agar from the plate used for plaque assay, and image the cells under a ChemiDoc. Expected results of Venus expression in the Chemidoc from Vero E6 cells infected with rSARS-CoV-2/WT and rSARS-CoV-2/Venus-2A are shown in **Fig. 6b (left column)**.

94. Visualize the plaques by immunostaining.

(i) Permeabilize cells with 2 mL of 0.5% TritonX-100 at room temperature for 10 min.

(ii) Block the cells by incubating in blocking solution (1 mL/well) at 37°C for 30 min.

(iii) Dilute the primary antibody against viral N protein (1C7C7) in blocking solution to a final concentration of 1 µg/mL, and incubate with the cells (1 mL/well) at 37°C for 1 h.

(iv) Wash the cells 3 times with PBS.

(v) Dilute the biotinylated horse anti-mouse secondary antibody in blocking solution, and incubate with the cells (1 mL/well) at 37°C for 1 h.

(vi) Prepare the ABC reagent by mixing 100 µL Reagent A and 100 µL Reagent B in 10 mL of PBS and incubate at 37°C for 30 min.

(vii) Wash the cells 3 times with PBS.

(viii) Incubate the cells with the ABC reagent (1 mL/well) at 37°C for 30 min.

(ix) Wah the cells 3 times with PBS.

(x) Prepare DAB substrate by mixing 2 drops of Reagent 1, 4 drops of Reagent 2, 2 drops of Reagent 3, and 2 drops of Reagent 4 in 5 mL deionized water.

(xi) Visualize the plaques by overlaying 1 mL of the prepared DAB substrate onto the cells immediately.

(xii) Stop the staining by washing the cells with PBS after discarding the substrate solution.

95. Image the immunostained wells under a ChemiDoc. Expected results from viral plaques immunostained with the 1C7C7 N protein antibody are shown in **Fig. 6b (right column)**.

## TIMING

### Section 1. Assembly of the full-length (FL) SARS-CoV-2 genome sequences into the empty pBeloBAC to generate pBeloBAC-FL

Steps 1-7, Preparation of the five fragments: 1 day.

Steps 8-12, Preparation of the empty pBeloBAC: 1 day.

Steps 13-20, Preparation of the linearized empty pBeloBAC: 1 day.

Steps 21-27, Assembly of the F1 into the linearized pBeloBAC: 3 days.

Steps 29-32, Sequential assembly of the F2-F5 into the pBeloBAC-F1: 12 days.

### Section 2. Rescue of rSARS-CoV-2 using the pBeloBAC-FL

Steps 33-37, Transfection of the pBeloBAC-FL into Vero E6 cells: 4 days.

Steps 38-46, Confirmation of rSARS-CoV-2/WT rescue: 4 days.

### Section 3. Cloning of the Venus reporter gene into the pBeloBAC-FL

Steps 47-62, Cloning of Venus into the pUC57-F1: 2 days.

Steps 63-82, Cloning of Venus into the pBeloBAC-FL: 3 days.

### Section 4. Recovery of the Venus-expressing rSARS-CoV-2 (rSARS-CoV-2/Venus-2A)

Steps 83-87, Rescue of rSARS-CoV-2/Venus-2A: 4 days.

Steps 88-95, Characterization of rSARS-CoV-2/Venus-2A: 4 days.

## TROUBLESHOOTING

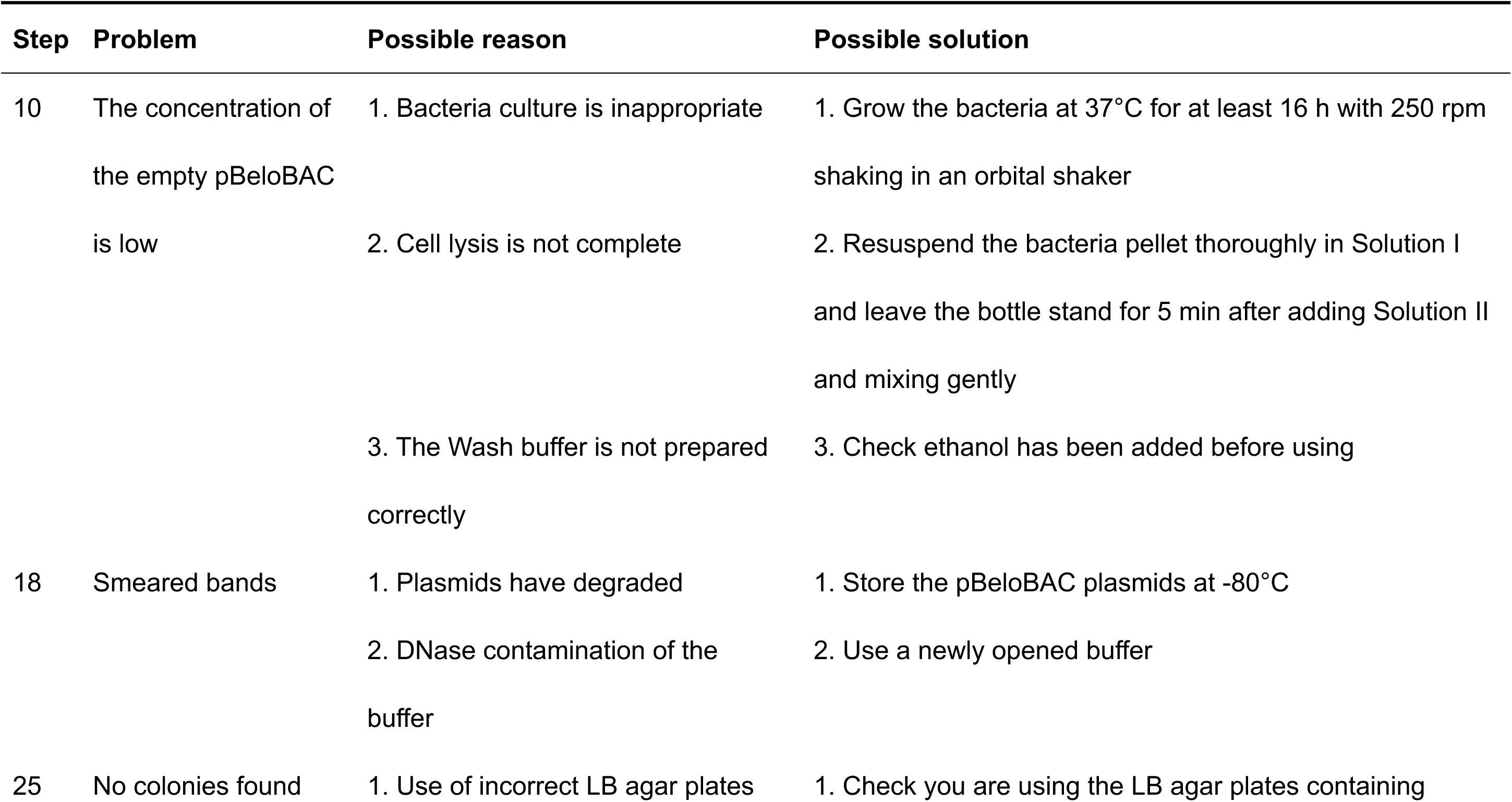
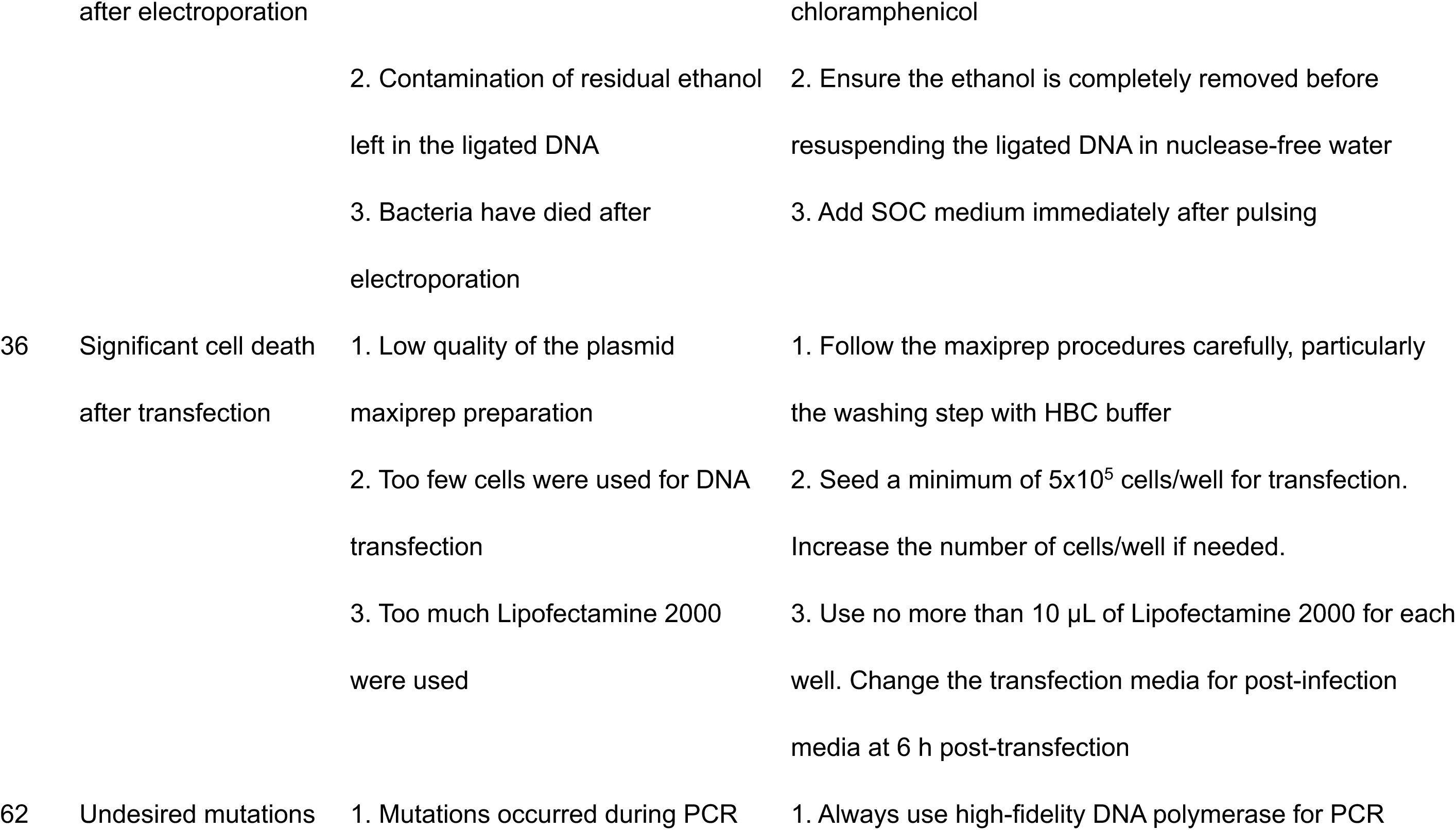
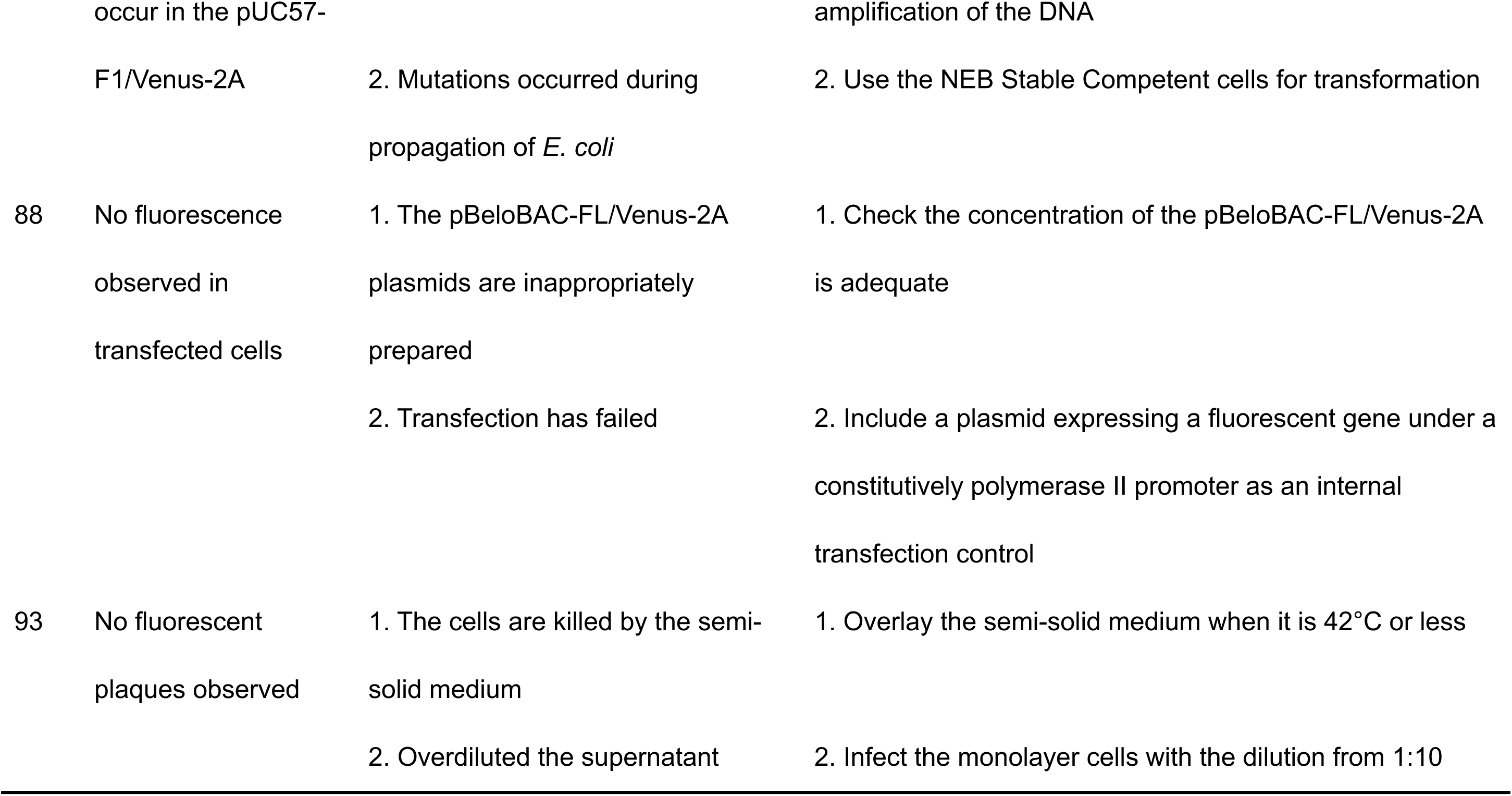

## ANTICIPATED RESULTS

This protocol can be used to successfully assemble the entire SARS-CoV-2 genome into the empty pBeloBAC, resulting in efficient recovery of rSARS-CoV-2/WT leading to apparent CPE on Vero E6 cells infected with cell culture supernatants from transfected cells (**Fig. 3b**). The rSARS-CoV-2/WT rescue can be further confirmed by IFA using a monoclonal antibody (1C7C7) against the viral N protein (**Fig. 3c**), and by Sanger sequencing the RT-PCR product of the M gene to demonstrate the recombinant nature of the rescued SARS-CoV-2 (**Fig. 3d**).

By using the pBeloBAC-FL as a backbone, this protocol efficiently introduces the reporter gene Venus into the pBeloBAC-FL and generates rSARS-CoV-2/Venus-2A expressing robust levels of Venus (**Fig. 6a**), allowing the virus to be readily detected as fluorescent plaques under a ChemiDoc imaging system (**Fig. 6b**).

The pBeloBAC-FL and pBeloBAC-FL/Venus-2A derivative for generation of rSARS-CoV-2/WT and rSARS-CoV-2/Venus-2A are available at this website: https://www.txbiomed.org/services-2/reverse-genetics-plasmids/. Other pBeloBAC constructs as well as their respective rSARS-CoV-2 are also available at the same website.

## AUTHOR CONTRIBUTIONS

C.Y and L. M.-S conceived the project, L. M.-S provided funding support, C.Y drafted the manuscript, and L. M.-S revised and finalized the manuscript.

## ACKNOWLEDGMENTS

We are thankful to Dr. Thomas Moran at the Icahn School of Medicine at Mount Sinai for providing the 1C7C7 monoclonal antibody. We are thankful also to Dr. Tracey Baas for criticizing reading and editing this manuscript.

