## Supplementary material for "Using a reverse genetics system to generate recombinant SARS-CoV-2 expressing robust levels of reporter genes"

**Fragment 1 (F1)**

CGTCTCTCATGGACATTGATTATTGACTAGTTATTAATAGTAATCAATTACGG
GGTCATTAGTTCATAGCCCATATATGGAGTTCCGCGTTACATAACTTACGGT
AAATGGCCCGCCTGGCTGACCGCCCAACGACCCCCGCCCATTGACGTC
AATAATGACGTATGTTCCCATAGTAACGCCAATAGGGACTTTCCATTGACG
TCAATGGGTGGAGTATTTACGGTAAACTGCCCACTTGGCAGTACATCAAG
TGTATCATATGCCAAGTACGCCCCCTATTGACGTCAATGACGGTAAATGGC
CCGCCTGGCATTATGCCCAGTACATGACCTTATGGGACTTTCCTACTTGG
CAGTACATCTACGTATTAGTCATCGCTATTACCATGGTGATGCGGTTTTGG
CAGTACATCAATGGGCGTGGATAGCGGTTTGACTCACGGGGATTTCCAAG
TCTCCACCCCATTGACGTCAATGGGAGTTTGTTTTGGCACCAAAATCAAC
GGGACTTTCCAAAATGTCGTAACAACTCCGCCCCATTGACGCAAATGGGC
GGTAGGCGTGTACGGTGGGAGGTCTATATAAGCAGAGCTCGTTTAGTGAA
CCGTATTAAAGGTTTATACCTTCCCAGGTAACAAACCAACCAACTTTCGAT
CTCTTGTAGATCTGTTCTCTAAACGAACTTTAAAATCTGTGTGGCTGTCAC
TCGGCTGCATGCTTAGTGCACTCACGCAGTATAATTAATAACTAATTACTGT
CGTTGACAGGACACGAGTAACTCGTCTATCTTCTGCAGGCTGCTTACGGT
TTCGTCCGTGTTGCAGCCGATCATCAGCACATCTAGGTTTCGTCCGGGTG
TGACCGAAAGGTAAGATGGAGAGCCTTGTCCCTGGTTTCAACGAGAAAA
CACACGTCCAACTCAGTTTGCCTGTTTTACAGGTTCGCGACGTGCTCGTA
CGTGGCTTTGGAGACTCCGTGGAGGAGGTCTTATCAGAGGCACGTCAAC
ATCTTAAAGATGGCACTTGTGGCTTAGTAGAAGTTGAAAAAGGCGTTTTG
CCTCAACTTGAACAGCCCTATGTGTTCATCAAACGTTCGGATGCTCGAAC
TGCACCTCATGGTCATGTTATGGTTGAGCTGGTAGCAGAACTCGAAGGCA
TTCAGTACGGTCGTAGTGGTGAGACACTTGGTGTCCTTGTCCCTCATGTG
GGCGAAATACCAGTGGCTTACCGCAAGGTTCTTCTTCGTAAGAACGGTAA
TAAAGGAGCTGGTGGCCATAGTTACGGCGCCATGTTAATTAACGCACGCG
TATATTCGAAATGGGATCCTGCTGCAAATTTGATGAAGACGACTCTGAGCC
AGTGCTCAAAGGAGTCAAATTACATTACACATAAACGAACTTATGGATTTG
TTTATGAGAATCTTCACAATTGGAACTGTAACTTTGAAGCAAGGTGAAATC
AAGGATGCTACTCCTTCAGATTTTGTTCGCGCTACTGCAACGATACCGATA
CAAGCCTCACTCCCTTTCGGATGGCTTATTGTTGGCGTTGCACTTCTTGC
TGTTTTTCAGAGCGCTTCCAAAATCATAACCCTCAAAAAGAGATGGCAACT
AGCACTCTCCAAGGGTGTTCACTTTGTTTGCAACTTGCTGTTGTTGTTTG
TAACAGTTTACTCACACCTTTTGCTCGTTGCTGCTGGCCTTGAAGCCCCT
TTTCTCTATCTTTATGCTTTAGTCTACTTCTTGCAGAGTATAAACTTTGTAAG
AATAATAATGAGGCTTTGGCTTTGCTGGAAATGCCGTTCCAAAAACCCATT
ACTTTATGATGCCAACTATTTTCTTTGCTGGCATACTAATTGTTACGACTATT
GTATACCTTACAATAGTGTAACTTCTTCAATTGTCATTACTTCAGGTGATGG
CACAACAAGTCCTATTTCTGAACATGACTACCAGATTGGTGGTTATACTGA
AAAATGGGAATCTGGAGTAAAAGACTGTGTTGTATTACACAGTTACTTCAC
TTCAGACTATTACCAGCTGTACTCAACTCAATTGAGTACAGACACTGGTGT
TGAACATGTTACCTTCTTCATCTACAATAAAATTGTTGATGAGCCTGAAGAA

CATGTCCAAATTCACACAATCGACGGTTCATCCGGAGTTGTTAATCCAGTA
ATGGAACCAATTTATGATGAACCGACGACGACTACTAGCGTGCCTTTGTAA
GCACAAGCTGATGAGTACGAACTTATGTACTCATTCGTTTCGGAAGAGAC
AGGTACGTTAATAGTTAATAGCGTACTTCTTTTTCTTGCTTTCGTGGTATTC
TTGCTAGTTACACTAGCCATCCTTACTGCGCTTCGATTGTGTGCGTACTGC
TGCAATATTGTTAACGTGAGTCTTGTAAAACCTTCTTTTTACGTTTACTCTC
GTGTTAAAAATCTGAATTCTTCTAGAGTTCCTGATCTTCTGGTCTAAACGA
ACTAAATATTATATTAGTTTTTCTGTTTGGAACTTTAATTTTAGCCATGGCAG
ATTCCAACGGTACTATTACCGTTGAAGAGCTTAAAAAGCTCCTTGAACAAT
GGAACCTAGTAATAGGTTTCCTATTCCTTACATGGATTTGTCTTCTACAATT
TGCCTATGCCAACAGGAATAGGTTTTTGTATATAATTAAGTTAATTTTCCTCT
GGCTGTTATGGCCAGTAACTTTAGCTTGTTTTGTGCTTGCTGCTGTTTACA
GAATAAATTGGATCACCGGTGGAATTGCTATCGCAATGGCTTGTCTTGTAG
GCTTGATGTGGCTCAGCTACTTCATTGCTTCTTTCAGACTGTTTGCGCGTA
CGCGATCCATGTGGTCATTCAATCCAGAAACTAACATTCTTCTCAACGTGC
CACTCCATGGCACTATTCTGACCAGACCGCTTCTAGAAAGTGAACTCGTA
ATCGGAGCTGTGATCCTTCGTGGACATCTTCGTATTGCTGGACACCATCT
AGGACGCTGTGACATCAAGGACCTGCCTAAAGAAATCACTGTTGCTACAT
CACGAACGCTTTCTTATTACAAATTGGGAGCTTCGCAGCGTGTAGCAGGT
GACTCAGGTTTTGCTGCATACAGTCGCTACAGGATTGGCAACTATAAATTA
AACACAGACCATTCCAGTAGCAGTGACAATATTGCTTTGCTTGTACAGTAA
GTGACAACAGATGTTTCATCTCGTTGACTTTCAGGTTACTATAGCAGAGAT
ATTACTAATTATTATGAGGACTTTTAAAGTTTCCATTTGGAATCTTGATTACA
TCATAAACCTCATAATTAAAAATTTATCTAAGTCACTAACTGAGAATAAATATT
CTCAATTAGATGAAGAGCAACCAATGGAGATTGATTAAACGAACATGAAAA
TTATTCTTTTCTTGGCACTGATAACACTCGCTACTTGTGAGCTTTATCACTA
CCAAGAGTGTGTTAGAGGTACAACAGTACTTTTAAAAGAACCTTGCTCTTC
TGGAACATACGAGGGCAATTCACCATTTCATCCTCTAGCTGATAACAAATT
TGCACTGACTTGCTTTAGCACTCAATTTGCTTTTGCTTGTCCTGACGGCG
TAAAACACGTCTATCAGTTACGTGCCAGATCAGTTTCACCTAAACTGTTCA
TCAGACAAGAGGAAGTTCAAGAACTTTACTCTCCAATTTTTCTTATTGTTG
CGGCAATAGTGTTTATAACACTTTGCTTCACACTCAAAAGAAAGACAGAAT
GATTGAACTTTCATTAATTGACTTCTATTTGTGCTTTTTAGCCTTTCTGCTAT
TCCTTGTTTTAATTATGCTTATTATCTTTTGGTTCTCACTTGAACTGCAAGAT
CATAATGAAACTTGTCACGCCTAAACGAACATGAAATTTCTTGTTTTCTTAG
GAATCATCACAACTGTAGCTGCATTTCACCAAGAATGTAGTTTACAGTCAT
GTACTCAACATCAACCATATGTAGTTGATGACCCGTGTCCTATTCACTTCTA
TTCTAAATGGTATATTAGAGTAGGAGCTAGAAAATCAGCACCTTTAATTGAA
TTGTGCGTGGATGAGGCTGGTTCTAAATCACCCATTCAGTACATCGATATC
GGTAATTATACAGTTTCCTGTTCACCTTTTACAATTAATTGCCAGGAACCTA
AATTGGGTAGTCTTGTAGTGCGTTGTTCGTTCTATGAAGACTTTTTAGAGT
ATCATGACGTTCGTGTTGTTTTAGATTTCATCTAAACGAACAAACTAAAATG
TCTGATAATGGACCCCAAAATCAGCGAAATGCACCCCGCATTACGTTTGG
TGGACCCTCAGATTCAACTGGCAGTAACCAGAATGGAGAACGCAGTGGG

GCGCGATCAAAACAACGTCGGCCCCAAGGTTTACCCAATAATACTGCGTC
TTGGTTCACCGCTCTCACTCAACATGGCAAGGAAGACCTTAAATTCCCTC
GAGGACAAGGCGTTCCAATTAACACCAATAGCAGTCCAGATGACCAAATT
GGCTACTACCGAAGAGCTACCAGACGAATTCGTGGTGGTGACGGTAAAAT
GAAAGATCTCAGTCCAAGATGGTATTTCTACTACCTAGGAACTGGGCCAG
AAGCTGGACTTCCCTATGGTGCTAACAAAGACGGCATCATATGGGTTGCA
ACTGAGGGAGCCTTGAATACACCAAAAGATCACATTGGCACCCGCAATCC
TGCTAACAATGCTGCAATCGTGCTACAACTTCCTCAAGGAACAACATTGC
CAAAAGGCTTCTACGCAGAAGGGAGCAGAGGCGGCAGTCAAGCCTCTTC
TCGTTCCTCATCACGTAGTCGCAACAGTTCAAGAAATTCAACTCCAGGCA
GCAGTAGGGGAACTTCTCCTGCTAGAATGGCTGGCAATGGCGGTGATGC
TGCTCTTGCTTTGCTGCTGCTTGACAGATTGAACCAGCTTGAGAGCAAAA
TGTCTGGTAAAGGCCAACAACAACAAGGCCAAACTGTCACTAAGAAATCT
GCTGCTGAGGCTTCTAAGAAGCCTCGGCAAAAACGTACTGCCACTAAAG
CATACAATGTAACACAAGCTTTCGGCAGACGTGGTCCAGAACAAACCCAA
GGAAATTTTGGGGACCAGGAACTAATCAGACAAGGAACTGATTACAAACA
TTGGCCGCAAATTGCACAATTTGCCCCCAGCGCTTCAGCGTTCTTCGGAA
TGTCGCGCATTGGCATGGAAGTCACACCTTCGGGAACGTGGTTGACCTA
CACAGGTGCCATCAAATTGGATGACAAAGATCCAAATTTCAAAGATCAAGT
CATTTTGCTGAATAAGCATATTGACGCATACAAAACATTCCCACCAACAGA
GCCTAAAAAGGACAAAAAGAAGAAGGCTGATGAAACTCAAGCCTTACCG
CAGAGACAGAAGAAACAGCAAACTGTGACTCTTCTTCCTGCTGCAGATTT
GGATGATTTCTCCAAACAATTGCAACAATCCATGAGCAGTGCTGACTCAA
CTCAGGCCTAAACTCATGCAGACCACACAAGGCAGATGGGCTATATAAAC
GTTTTCGCTTTTCCGTTTACGATATATAGTCTACTCTTGTGCAGAATGAATT
CTCGTAACTACATAGCACAAGTAGATGTAGTTAACTTTAATCTCACATAGCA
ATCTTTAATCAGTGTGTAACATTAGGGAGGACTTGAAAGAGCCACCACATT
TTCACCGAGGCCACGCGGAGTACGATCGAGTGTACAGTGAACAATGCTA
GGGAGAGCTGCCTATATGGAAGAGCCCTAATGTGTAAAATTAATTTTAGTA
GTGCTATCCCCATGTGATTTTAATAGCTTCTTAGGAGAATGACAAAAAAAA
AAAAAAAAAAAAAAAAAAAAAAAA*gggtcggcatggcatctccacctcctcgcggtccgac*
*ctgggcatccgaaggaggacgtcgtccactcggatggctaagggagagctcggatcgatccgcctcga*ct
gtgccttctagttgccagccatctgttgtttgcccctcccccgtgccttccttgaccctggaaggtgccactccca
ctgtcctttcctaataaaatgaggaaattgcatcgcattgtctgagtaggtgtcattctattctggggggtggggt
ggggcaggacagcaagggggaggattgggaagacaatagcaggcatgctggggaAGCTAGAGA
CG

- The underlined sequences in capital (CGTCTCT and AGAGACG) indicate the BsmBI restriction site used to generate overhangs that are compatible with PciI (ACATGT) and HindIII (AAGCTT) (**Figure 1**).
- The sequence highlighted in red represents the cytomegalovirus (CMV) promoter.
- The sequences highlighted in yellow represent unique KasI (GGCGCC), PacI (TTAATTAA), MluI (ACGCGT), BstBI (TTCGAA), and BamHI

(GGATCC) restriction sites use to clone F2-F5 (**Figure 1**).

- The single nucleotide highlighted in green indicates a silent mutation introduced to remove a MluI (ACGCGT) restriction site present in the viral membrane (M) gene to clone F3 (**Figure 1**). This single nucleotide was also used as a genetic tag to distinguish the rSARS-CoV-2 from the natural SARS-CoV-2 isolate.
- The sequence shown in lower case italic font indicates the hepatitis delta virus (HDV) ribozyme (Rz) sequence.
- The lower case underlined sequence represents the bovine growth hormone (bGH) polyadenylation signal.

**Fragment 2 (F2)**

GGCGCCGATCTAAAGTCATTTGACTTAGGCGACGAGCTTGGCACTGATCC
TTATGAAGATTTTCAAGAAAACTGGAACACTAAACATAGCAGTGGTGTTAC
CCGTGAACTCATGCGTGAGCTTAACGGAGGGGCATACACTCGCTATGTC
GATAACAACTTCTGTGGCCCTGATGGCTACCCTCTTGAGTGCATTAAAGA
CCTTCTAGCACGTGCTGGTAAAGCTTCATGCACTTTGTCCGAACAACTGG
ACTTTATTGACACTAAGAGGGGTGTATACTGCTGCCGTGAACATGAGCAT
GAAATTGCTTGGTACACGGAACGTTCTGAAAAGAGCTATGAATTGCAGAC
ACCTTTTGAAATTAAATTGGCAAAGAAATTTGACACCTTCAATGGGGAATG
TCCAAATTTTGTATTTCCCTTAAATTCCATAATCAAGACTATTCAACCAAGG
GTTGAAAAGAAAAAGCTTGATGGCTTTATGGGTAGAATTCGATCTGTCTAT
CCAGTTGCGTCACCAAATGAATGCAACCAAATGTGCCTTTCAACTCTCAT
GAAGTGTGATCATTGTGGTGAAACTTCATGGCAGACGGGCGATTTTGTTA
AAGCCACTTGCGAATTTTGTGGCACTGAGAATTTGACTAAAGAAGGTGCC
ACTACTTGTGGTTACTTACCCCAAAATGCTGTTGTTAAAATTTATTGTCCAG
CATGTCACAATTCAGAAGTAGGACCTGAGCATAGTCTTGCCGAATACCATA
ATGAATCTGGCTTGAAAACCATTCTTCGTAAGGGTGGTCGCACTATTGCC
TTTGGAGGCTGTGTGTTCTCTTATGTTGGTTGCCATAACAAGTGTGCCTAT
TGGGTTCCACGTGCTAGCGCTAACATAGGTTGTAACCATACAGGTGTTGT
TGGAGAAGGTTCCGAAGGTCTTAATGACAACCTTCTTGAAATACTCCAAA
AAGAGAAAGTCAACATCAATATTGTTGGTGACTTTAAACTTAATGAAGAGA
TCGCCATTATTTTGGCATCTTTTTCTGCTTCCACAAGTGCTTTTGTGGAAA
CTGTGAAAGGTTTGGATTATAAAGCATTCAAACAAATTGTTGAATCCTGTG
GTAATTTTAAAGTTACAAAAGGAAAAGCTAAAAAAGGTGCCTGGAATATTG
GTGAACAGAAATCAATACTGAGTCCTCTTTATGCATTTGCATCAGAGGCTG
CTCGTGTTGTACGATCAATTTTCTCCCGCACTCTTGAAACTGCTCAAAATT
CTGTGCGTGTTTTACAGAAGGCCGCTATAACAATACTAGATGGAATTTCAC
AGTATTCACTGAGACTCATTGATGCTATGATGTTCACATCTGATTTGGCTAC
TAACAATCTAGTTGTAATGGCCTACATTACAGGTGGTGTTGTTCAGTTGAC
TTCGCAGTGGCTAACTAACATCTTTGGCACTGTTTATGAAAAACTCAAACC
CGTCCTTGATTGGCTTGAAGAGAAGTTTAAGGAAGGTGTAGAGTTTCTTA
GAGACGGTTGGGAAATTGTTAAATTTATCTCAACCTGTGCTTGTGAAATTG
TCGGTGGACAAATTGTCACCTGTGCAAAGGAAATTAAGGAGAGTGTTCAG
ACATTCTTTAAGCTTGTAAATAAATTTTTGGCTTTGTGTGCTGACTCTATCA
TTATTGGTGGAGCTAAACTTAAAGCCTTGAATTTAGGTGAAACATTTGTCA
CGCACTCAAAGGGATTGTACAGAAAGTGTGTTAAATCCAGAGAAGAAACT
GGCCTACTCATGCCTCTAAAAGCCCCAAAAGAAATTATCTTCTTAGAGGGA
GAAACACTTCCCACAGAAGTGTTAACAGAGGAAGTTGTCTTGAAAACTGG
TGATTTACAACCATTAGAACAACCTACTAGTGAAGCTGTTGAAGCTCCATT
GGTTGGTACACCAGTTTGTATTAACGGGCTTATGTTGCTCGAAATCAAAGA
CACAGAAAAGTACTGTGCCCTTGCACCTAATATGATGGTAACAAACAATAC
CTTCACACTCAAAGGCGGTGCACCAACAAAGGTTACTTTTGGTGATGACA
CTGTGATAGAAGTGCAAGGTTACAAGAGTGTGAATATCACTTTTGAACTTG

ATGAAAGGATTGATAAAGTACTTAATGAGAAGTGCTCTGCCTATACAGTTG
AACTCGGTACAGAAGTAAATGAGTTCGCCTGTGTTGTGGCAGATGCTGTC
ATAAAAACTTTGCAACCAGTATCTGAATTACTTACACCACTGGGCATTGATT
TAGATGAGTGGAGTATGGCTACATACTACTTATTTGATGAGTCTGGTGAGT
TTAAATTGGCTTCACATATGTATTGTTCTTTCTACCCTCCAGATGAGGATGA
AGAAGAAGGTGATTGTGAAGAAGAAGAGTTTGAGCCATCAACTCAATATG
AGTATGGTACTGAAGATGATTACCAAGGTAAACCTTTGGAATTTGGTGCCA
CTTCTGCTGCTCTTCAACCTGAAGAAGAGCAAGAAGAAGATTGGTTAGAT
GATGATAGTCAACAAACTGTTGGTCAACAAGACGGCAGTGAGGACAATCA
GACAACTACTATTCAAACAATTGTTGAGGTTCAACCTCAATTAGAGATGGA
ACTTACACCAGTTGTTCAGACTATTGAAGTGAATAGTTTTAGTGGTTATTTA
AAACTTACTGACAATGTATACATTAAAAATGCAGACATTGTGGAAGAAGCTA
AAAAGGTAAAACCAACAGTGGTTGTTAATGCAGCCAATGTTTACCTTAAAC
ATGGAGGAGGTGTTGCAGGAGCCTTAAATAAGGCTACTAACAATGCCATG
CAAGTTGAATCTGATGATTACATAGCTACTAATGGACCACTTAAAGTGGGT
GGTAGTTGTGTTTTAAGCGGACACAATCTTGCTAAACACTGTCTTCATGTT
GTCGGCCCAAATGTTAACAAAGGTGAAGACATTCAACTTCTTAAGAGTGC
TTATGAAAATTTTAATCAGCACGAAGTTCTACTTGCACCATTATTATCAGCT
GGTATTTTTGGTGCTGACCCTATACATTCTTTAAGAGTTTGTGTAGATACTG
TTCGCACAAATGTCTACTTAGCTGTCTTTGATAAAAATCTCTATGACAAACT
TGTTTCAAGCTTTTTGGAAATGAAGAGTGAAAAGCAAGTTGAACAAAAGA
TCGCTGAGATTCCTAAAGAGGAAGTTAAGCCATTTATAACTGAAAGTAAAC
CTTCAGTTGAACAGAGAAAACAAGATGATAAGAAAATCAAAGCTTGTGTTG
AAGAAGTTACAACAACTCTGGAAGAAACTAAGTTCCTCACAGAAAACTTG
TTACTTTATATTGACATTAATGGCAATCTTCATCCAGATTCTGCCACTCTTG
TTAGTGACATTGACATCACTTTCTTAAAGAAAGATGCTCCATATATAGTGGG
TGATGTTGTTCAAGAGGGTGTTTTAACTGCTGTGGTTATACCTACTAAAAA
GGCTGGTGGCACTACTGAAATGCTAGCGAAAGCTTTGAGAAAAGTGCCA
ACAGACAATTATATAACCACTTACCCGGGTCAGGGTTTAAATGGTTACACT
GTAGAGGAGGCAAAGACAGTGCTTAAAAAGTGTAAAAGTGCCTTTTACAT
TCTACCATCTATTATCTCTAATGAGAAGCAAGAAATTCTTGGAACTGTTTCT
TGGAATTTGCGAGAAATGCTTGCACATGCAGAAGAAACACGCAAATTAAT
GCCTGTCTGTGTGGAAACTAAAGCCATAGTTTCAACTATACAGCGTAAATA
TAAGGGTATTAAAATACAAGAGGGTGTGGTTGATTATGGTGCTAGATTTTA
CTTTTACACCAGTAAAACAACTGTAGCGTCACTTATCAACACACTTAACGA
TCTAAATGAAACTCTTGTTACAATGCCACTTGGCTATGTAACACATGGCTTA
AATTTGGAAGAAGCTGCTCGGTATATGAGATCTCTCAAAGTGCCAGCTAC
AGTTTCTGTTTCTTCACCTGATGCTGTTACAGCGTATAATGGTTATCTTACT
TCTTCTTCTAAAACACCTGAAGAACATTTTATTGAAACCATCTCACTTGCTG
GTTCCTATAAAGATTGGTCCTATTCTGGACAATCTACACAACTAGGTATAGA
ATTTCTTAAGAGAGGTGATAAAAGTGTATATTACACTAGTAATCCTACCACA
TTCCACCTAGATGGTGAAGTTATCACCTTTGACAATCTTAAGACACTTCTT
TCTTTGAGAGAAGTGAGGACTATTAAGGTGTTTACAACAGTAGACAACATT
AACCTCCACACGCAAGTTGTGGACATGTCAATGACATATGGACAACAGTT

TGGTCCAACTTATTTGGATGGAGCTGATGTTACTAAAATAAAACCTCATAAT
TCACATGAAGGTAAAACATTTTATGTTTTACCTAATGATGACACTCTACGTG
TTGAGGCTTTTGAGTACTACCACACAACTGATCCTAGTTTTCTGGGTAGGT
ACATGTCAGCATTAAATCACACTAAAAAGTGGAAATACCCACAAGTTAATG
GTTTAACTTCTATTAAATGGGCAGATAACAACTGTTATCTTGCCACTGCATT
GTTAACACTCCAACAAATAGAGTTGAAGTTTAATCCACCTGCTCTACAAGA
TGCTTATTACAGAGCAAGGGCTGGTGAAGCTGCTAACTTTTGTGCACTTAT
CTTAGCCTACTGTAATAAGACAGTAGGTGAGTTAGGTGATGTTAGAGAAAC
AATGAGTTACTTGTTTCAACATGCCAATTTAGATTCTTGCAAAAGAGTCTT
GAACGTGGTGTGTAAAACTTGTGGACAACAGCAGACAACCCTTAAGGGT
GTAGAAGCTGTTATGTACATGGGCACACTTTCTTATGAACAATTTAAGAAA
GGTGTTCAGATACCTTGTACGTGTGGTAAACAAGCTACAAAATATCTAGTA
CAACAGGAGTCACCTTTTGTTATGATGTCAGCACCACCTGCTCAGTATGA
ACTTAAGCATGGTACATTTACTTGTGCTAGTGAGTACACTGGTAATTACCA
GTGTGGTCACTATAAACATATAACTTCTAAAGAAACTTTGTATTGCATAGAC
GGTGCTTTACTTACAAAGTCCTCAGAATACAAAGGTCCTATTACGGATGTT
TTCTACAAAGAAAACAGTTACACAACAACCATAAAACCAGTTACTTATAAAT
TGGATGGTGTTGTTTGTACAGAAATTGACCCTAAGTTGGACAATTATTATAA
GAAAGACAATTCTTATTTCACAGAGCAACCAATTGATCTTGTACCAAACCA
ACCATATCCAAACGCAAGCTTCGATAATTTTAAGTTTGTATGTGATAATATCA
AATTTGCTGATGATTTAAACCAGTTAACTGGTTATAAGAAACCTGCTTCAAG
AGAGCTTAAAGTTACATTTTTCCCTGACTTAAATGGTGATGTGGTGGCTAT
TGATTATAAACACTACACACCCTCTTTTAAGAAAGGAGCTAAATTGTTACAT
AAACCTATTGTTTGGCATGTTAACAATGCAACTAATAAAGCCACGTATAAAC
CAAATACCTGGTGTATACGTTGTCTTTGGAGCACAAAACCAGTTGAAACAT
CAAATTCGTTTGATGTACTGAAGTCAGAGGACGCGCAGGGAATGGATAAT
CTTGCCTGCGAAGATCTAAAACCAGTCTCTGAAGAAGTAGTGGAAAATCC
TACCATACAGAAAGACGTTCTTGAGTGTAATGTGAAAACTACCGAAGTTGT
AGGAGACATTATACTTAAACCAGCAAATAATAGTTTAAAAATTACAGAAGAG
GTTGGCCACACAGATCTAATGGCTGCTTATGTAGACAATTCTAGTCTTACT
ATTAAGAAACCTAATGAATTATCTAGAGTATTAGGTTTGAAAACCCTTGCTA
CTCATGGTTTAGCTGCTGTTAATAGTGTCCCTTGGGATACTATAGCTAATTA
TGCTAAGCCTTTTCTTAACAAAGTTGTTAGTACAACTACTAACATAGTTACA
CGGTGTTTAAACCGTGTTTGTACTAATTATATGCCTTATTTCTTTACTTTATT
GCTACAATTGTGTACTTTTACTAGAAGTACAAATTCTAGAATTAAAGCATCT
ATGCCGACTACTATAGCAAAGAATACTGTTAAGAGTGTCGGTAAATTTTGT
CTAGAGGCTTCATTTAATTATTTGAAGTCACCTAATTTTTCTAAACTGATAAA
TATTATAATTTGGTTTTTACTATTAAGTGTTTGCCTAGGTTCTTTAATCTACTC
AACCGCTGCTTTAGGTGTTTTAATGTCTAATTTAGGCATGCCTTCTTACTGT
ACTGGTTACAGAGAAGGCTATTTGAACTCTACTAATGTCACTATTGCAACC
TACTGTACTGGTTCTATACCTTGTAGTGTTTGTCTTAGTGGTTTAGATTCTT
TAGACACCTATCCTTCTTTAGAAACTATACAAATTACCATTTCATCTTTTAAA
TGGGATTTAACTGCTTTTGGCTTAGTTGCAGAGTGGTTTTTGGCATATATT
CTTTTCACTAGGTTTTTCTATGTACTTGGATTGGCTGCAATCATGCAATTGT

TTTTCAGCTATTTTGCAGTACATTTTATTAGTAATTCTTGGCTTATGTGGTTA
ATAATTAATCTTGTACAAATGGCCCCGATTTCAGCTATGGTTAGAATGTACA
TCTTCTTTGCATCATTTTATTATGTATGGAAAAGTTATGTGCATGTTGTAGAC
GGTTGTAATTCATCAACTTGTATGATGTGTTACAAACGTAATAGAGCAACAA
GAGTCGAATGTACAACTATTGTTAATGGTGTTAGAAGGTCCTTTTATGTCTA
TGCTAATGGAGGTAAAGGCTTTTGCAAACTACACAATTGGAATTGTGTTAA
TTGTGATACATTCTGTGCTGGTAGTACATTTATTAGTGATGAAGTTGCGAG
AGACTTGTCACTACAGTTTAAAAGACCAATAAATCCTACTGACCAGTCTTC
TTACATCGTTGATAGTGTTACAGTGAAGAATGGTTCCATCCATCTTTACTTT
GATAAAGCTGGTCAAAAGACTTATGAAAGACATTCTCTCTCTCATTTTGTTA
ACTTAGACAACCTGAGAGCTAATAACACTAAAGGTTCATTGCCTATTAATGT
TATAGTTTTTGATGGTAAATCAAAATGTGAAGAATCATCTGCAAAATCAGCG
TCTGTTTACTACAGTCAGCTTATGTGTCAACCTATACTGTTACTAGATCAGG
CATTAGTGTCTGATGTTGGTGATAGTGCGGAAGTTGCAGTTAAAATGTTTG
ATGCTTACGTTAATACGTTTTCATCAACTTTTAACGTACCAATGGAAAAACT
CAAAACACTAGTTGCAACTGCAGAAGCTGAACTTGCAAAGAATGTGTCCT
TAGACAATGTCTTATCTACTTTTATTTCAGCAGCTCGGCAAGGGTTTGTTG
ATTCAGATGTAGAAACTAAAGATGTTGTTGAATGTCTTAAATTGTCACATCA
ATCTGACATAGAAGTTACTGGCGATAGTTGTAATAACTATATGCTCACCTAT
AACAAAGTTGAAAACATGACACCCCGTGACCTTGGTGCTTGTATTGACTG
TAGTGCGCGTCATATTAATGCGCAGGTAGCAAAAAGTCACAACATTGCTTT
GATATGGAACGTTAAAGATTTCATGTCATTGTCTGAACAACTACGAAAACA
AATACGTAGTGCTGCTAAAAAGAATAACTTACCTTTTAAGTTGACATGTGCA
ACTACTAGACAAGTTGTTAATGTTGTAACAACAAAGATAGCACTTAAGGGT
GGTAAAATTGTTAATAATTGGTTGAAGCAGTTAATTAA

- The sequences highlighted in yellow indicate unique Kas I (GGCGCC) and PacI (TTAATTAA) restriction sites used to clone F2 (**Figure 1**).

**Fragment 3 (F3)**

TTAATTAAAGTTACACTTGTGTTCCTTTTTGTTGCTGCTATTTTCTATTTAAT
AACACCTGTTCATGTCATGTCTAAACATACTGACTTTTCAAGTGAAATCATA
GGATACAAGGCTATTGATGGTGGTGTCACTCGTGACATAGCATCTACAGAT
ACTTGTTTTGCTAACAAACATGCTGATTTTGACACATGGTTTAGTCAGCGT
GGTGGTAGTTATACTAATGACAAAGCTTGCCCATTGATTGCTGCAGTCATA
ACAAGAGAAGTGGGTTTTGTCGTGCCTGGTTTGCCTGGCACGATATTACG
CACAACTAATGGTGACTTTTTGCATTTCTTACCTAGAGTTTTTAGTGCAGTT
GGTAACATCTGTTACACACCATCAAAACTTATAGAGTACACTGACTTTGCA
ACATCAGCTTGTGTTTTGGCTGCTGAATGTACAATTTTTAAAGATGCTTCT
GGTAAGCCAGTACCATATTGTTATGATACCAATGTACTAGAAGGTTCTGTT
GCTTATGAAAGTTTACGCCCTGACACACGTTATGTGCTCATGGATGGCTCT
ATTATTCAATTTCCTAACACCTACCTTGAAGGTTCTGTTAGAGTGGTAACAA
CTTTTGATTCTGAGTACTGTAGGCACGGCACTTGTGAAAGATCAGAAGCT
GGTGTTTGTGTATCTACTAGTGGTAGATGGGTACTTAACAATGATTATTACA
GATCTTTACCAGGAGTTTTCTGTGGTGTAGATGCTGTAAATTTACTTACTAA
TATGTTTACACCACTAATTCAACCTATTGGTGCTTTGGACATATCAGCATCT
ATAGTAGCTGGTGGTATTGTAGCTATCGTAGTAACATGCCTTGCCTACTATT
TTATGAGGTTTAGAAGAGCTTTTGGTGAATACAGTCATGTAGTTGCCTTTA
ATACTTTACTATTCCTTATGTCATTCACTGTACTCTGTTTAACACCAGTTTAC
TCATTCTTACCTGGTGTTTATTCTGTTATTTACTTGTACTTGACATTTTATCT
TACTAATGATGTTTCTTTTTTAGCACATATTCAGTGGATGGTTATGTTCACA
CCTTTAGTACCTTTCTGGATAACAATTGCTTATATCATTTGTATTTCCACAAA
GCATTTCTATTGGTTCTTTAGTAATTACCTAAAGAGACGTGTAGTCTTTAAT
GGTGTTTCCTTTAGTACTTTTGAAGAAGCTGCGCTGTGCACCTTTTTGTTA
AATAAAGAAATGTATCTAAAGTTGCGTAGTGATGTGCTATTACCTCTTACGC
AATATAATAGATACTTAGCTCTTTATAATAAGTACAAGTATTTTAGTGGAGCA
ATGGATACAACTAGCTACAGAGAAGCTGCTTGTTGTCATCTCGCAAAGGC
TCTCAATGACTTCAGTAACTCAGGTTCTGATGTTCTTTACCAACCACCACA
AACCTCTATCACCTCAGCTGTTTTGCAGAGTGGTTTTAGAAAAATGGCATT
CCCATCTGGTAAAGTTGAGGGTTGTATGGTACAAGTAACTTGTGGTACAA
CTACACTTAACGGTCTTTGGCTTGATGACGTAGTTTACTGTCCAAGACATG
TGATCTGCACCTCTGAAGACATGCTTAACCCTAATTATGAAGATTTACTCAT
TCGTAAGTCTAATCATAATTTCTTGGTACAGGCTGGTAATGTTCAACTCAG
GGTTATTGGACATTCTATGCAAAATTGTGTACTTAAGCTTAAGGTTGATACA
GCCAATCCTAAGACACCTAAGTATAAGTTTGTTCGCATTCAACCAGGACAG
ACTTTTTCAGTGTTAGCTTGTTACAATGGTTCACCATCTGGTGTTTACCAAT
GTGCTATGAGGCCCAATTTCACTATTAAGGGTTCATTCCTTAATGGTTCAT
GTGGTAGTGTTGGTTTTAACATAGATTATGACTGTGTCTCTTTTTGTTACAT
GCACCATATGGAATTACCAACTGGAGTTCATGCTGGCACAGACTTAGAAG
GTAACTTTTATGGACCTTTTGTTGACAGGCAAACAGCACAAGCAGCTGGT
ACGGACACAACTATTACAGTTAATGTTTTAGCTTGGTTGTACGCTGCTGTT
ATAAATGGAGACAGGTGGTTTCTCAATCGATTTACCACAACTCTTAATGAC

TTTAACCTTGTGGCTATGAAGTACAATTATGAACCTCTAACACAAGACCAT
GTTGACATACTAGGACCTCTTTCTGCTCAAACTGGAATTGCCGTTTTAGAT
ATGTGTGCTTCATTAAAAGAATTACTGCAAAATGGTATGAATGGACGTACC
ATATTGGGTAGTGCTTTATTAGAAGATGAATTTACACCTTTTGATGTTGTTA
GACAATGCTCAGGTGTTACTTTCCAAAGTGCAGTGAAAAGAACAATCAAG
GGTACACACCACTGGTTGTTACTCACAATTTTGACTTCACTTTTAGTTTTA
GTCCAGAGTACTCAATGGTCTTTGTTCTTTTTTTTGTATGAAAATGCCTTTT
TACCTTTTGCTATGGGTATTATTGCTATGTCTGCTTTTGCAATGATGTTTGT
CAAACATAAGCATGCATTTCTCTGTTTGTTTTTGTTACCTTCTCTTGCCACT
GTAGCTTATTTTAATATGGTCTATATGCCTGCTAGTTGGGTGATGCGTATTAT
GACATGGTTGGATATGGTTGATACTAGTTTGTCTGGTTTTAAGCTAAAAGA
CTGTGTTATGTATGCATCAGCTGTAGTGTTACTAATCCTTATGACAGCAAGA
ACTGTGTATGATGATGGTGCTAGGAGAGTGTGGACACTTATGAATGTCTT
GACACTCGTTTATAAAGTTTATTATGGTAATGCTTTAGATCAAGCCATTTCC
ATGTGGGCTCTTATAATCTCTGTTACTTCTAACTACTCAGGTGTAGTTACAA
CTGTCATGTTTTTGGCCAGAGGTATTGTTTTTATGTGTGTTGAGTATTGCC
CTATTTTCTTCATAACTGGTAATACACTTCAGTGTATAATGCTAGTTTATTGT
TTCTTAGGCTATTTTTGTACTTGTTACTTTGGCCTCTTTTGTTTACTCAACC
GCTACTTTAGACTGACTCTTGGTGTTTATGATTACTTAGTTTCTACACAGGA
GTTTAGATATATGAATTCACAGGGACTACTCCCACCCAAGAATAGCATAGAT
GCCTTCAAACTCAACATTAAATTGTTGGGTGTTGGTGGCAAACCTTGTATC
AAAGTAGCCACTGTACAGTCTAAAATGTCAGATGTAAAGTGCACATCAGTA
GTCTTACTCTCAGTTTTGCAACAACTCAGAGTAGAATCATCATCTAAATTGT
GGGCTCAATGTGTCCAGTTACACAATGACATTCTCTTAGCTAAAGATACTA
CTGAAGCCTTTGAAAAAATGGTTTCACTACTTTCTGTTTTGCTTTCCATGC
AGGGTGCTGTAGACATAAACAAGCTTTGTGAAGAAATGCTGGACAACAGG
GCAACCTTACAAGCTATAGCCTCAGAGTTTAGTTCCCTTCCATCATATGCA
GCTTTTGCTACTGCTCAAGAAGCTTATGAGCAGGCTGTTGCTAATGGTGA
TTCTGAAGTTGTTCTTAAAAAGTTGAAGAAGTCTTTGAATGTGGCTAAATC
TGAATTTGACCGTGATGCAGCCATGCAACGTAAGTTGGAAAAGATGGCTG
ATCAAGCTATGACCCAAATGTATAAACAGGCTAGATCTGAGGACAAGAGG
GCAAAAGTTACTAGTGCTATGCAGACAATGCTTTTCACTATGCTTAGAAAG
TTGGATAATGATGCACTCAACAACATTATCAACAATGCAAGAGATGGTTGT
GTTCCCTTGAACATAATACCTCTTACAACAGCAGCCAAACTAATGGTTGTC
ATACCAGACTATAACACATATAAAAATACGTGTGATGGTACAACATTTACTTA
TGCATCAGCATTGTGGGAAATCCAACAGGTTGTAGATGCAGATAGTAAAAT
TGTTCAACTTAGTGAAATTAGTATGGACAATTCACCTAATTTAGCATGGCCT
CTTATTGTAACAGCTTTAAGGGCCAATTCTGCTGTCAAATTACAGAATAATG
AGCTTAGTCCTGTTGCACTACGACAGATGTCTTGTGCTGCCGGTACTACA
CAAACTGCTTGCACTGATGACAATGCGTTAGCTTACTACAACACAACAAA
GGGAGGTAGGTTTGTACTTGCACTGTTATCCGATTTACAGGATTTGAAATG
GGCTAGATTCCCTAAGAGTGATGGAACTGGTACTATCTATACAGAACTGGA
ACCACCTTGTAGGTTTGTTACAGACACACCTAAAGGTCCTAAAGTGAAGT
ATTTATACTTTATTAAAGGATTAAACAACCTAAATAGAGGTATGGTACTTGGT

AGTTTAGCTGCCACAGTACGTCTACAAGCTGGTAATGCAACAGAAGTGCC
TGCCAATTCAACTGTATTATCTTTCTGTGCTTTTGCTGTAGATGCTGCTAAA
GCTTACAAAGATTATCTAGCTAGTGGGGGACAACCAATCACTAATTGTGTT
AAGATGTTGTGTACACACACTGGTACTGGTCAGGCAATAACAGTTACACC
GGAAGCCAATATGGATCAAGAATCCTTTGGTGGTGCATCGTGTTGTCTGT
ACTGCCGTTGCCACATAGATCATCCAAATCCTAAAGGATTTTGTGACTTAA
AAGGTAAGTATGTACAAATACCTACAACTTGTGCTAATGACCCTGTGGGTT
TTACACTTAAAAACACAGTCTGTACCGTCTGCGGTATGTGGAAAGGTTATG
GCTGTAGTTGTGATCAACTCCGCGAACCCATGCTTCAGTCAGCTGATGCA
CAATCGTTTTTAAACGGGTTTGCGGTGTAAGTGCAGCCCGTCTTACACCG
TGCGGCACAGGCACTAGTACTGATGTCGTATACAGGGCTTTTGACATCTA
CAATGATAAAGTAGCTGGTTTTGCTAAATTCCTAAAAACTAATTGTTGTCGC
TTCCAAGAAAAGGACGAAGATGACAATTTAATTGATTCTTACTTTGTAGTTA
AGAGACACACTTTCTCTAACTACCAACATGAAGAAACAATTTATAATTTACT
TAAGGATTGTCCAGCTGTTGCTAAACATGACTTCTTTAAGTTTAGAATAGA
CGGTGACATGGTACCACATATATCACGTCAACGTCTTACTAAATACACAAT
GGCAGACCTCGTCTATGCTTTAAGGCATTTTGATGAAGGTAATTGTGACAC
ATTAAAAGAAATACTTGTCACATACAATTGTTGTGATGATGATTATTTCAATA
AAAAGGACTGGTATGATTTTGTAGAAAACCCAGATATATTACGCGT

- The sequences highlighted in yellow indicate unique PacI (TTAATTAA) and MluI (ACGCGT) restriction sites used to clone F3 (**Figure 1**).

**Fragment 4 (F4)**

ACGCGTATACGCCAACTTAGGTGAACGTGTACGCCAAGCTTTGTTAAAAA
CAGTACAATTCTGTGATGCCATGCGAAATGCTGGTATTGTTGGTGTACTGA
CATTAGATAATCAAGATCTCAATGGTAACTGGTATGATTTCGGTGATTTCAT
ACAAACCACGCCAGGTAGTGGAGTTCCTGTTGTAGATTCTTATTATTCATT
GTTAATGCCTATATTAACCTTGACCAGGGCTTTAACTGCAGAGTCACATGT
TGACACTGACTTAACAAAGCCTTACATTAAGTGGGATTTGTTAAAATATGAC
TTCACGGAAGAGAGGTTAAAACTCTTTGACCGTTATTTTAAATATTGGGAT
CAGACATACCACCCAAATTGTGTTAACTGTTTGGATGACAGATGCATTCTG
CATTGTGCAAACTTTAATGTTTTATTCTCTACAGTGTTCCCACCTACAAGTT
TTGGACCACTAGTGAGAAAAATATTTGTTGATGGTGTTCCATTTGTAGTTT
CAACTGGATACCACTTCAGAGAGCTAGGTGTTGTACATAATCAGGATGTAA
ACTTACATAGCTCTAGACTTAGTTTTAAGGAATTACTTGTGTATGCTGCTGA
CCCTGCTATGCACGCTGCTTCTGGTAATCTATTACTAGATAAACGCACTAC
GTGCTTTTCAGTAGCTGCACTTACTAACAATGTTGCTTTTCAAACTGTCAA
ACCCGGTAATTTTAACAAAGACTTCTATGACTTTGCTGTGTCTAAGGGTTT
CTTTAAGGAAGGAAGTTCTGTTGAATTAAAACACTTCTTCTTTGCTCAGGA
TGGTAATGCTGCTATCAGCGATTATGACTACTATCGTTATAATCTACCAACA
ATGTGTGATATCAGACAACTACTATTTGTAGTTGAAGTTGTTGATAAGTACT
TTGATTGTTACGATGGTGGCTGTATTAATGCTAACCAAGTCATCGTCAACA
ACCTAGACAAATCAGCTGGTTTTCCATTTAATAAATGGGGTAAGGCTAGAC
TTTATTATGATTCAATGAGTTATGAGGATCAAGATGCACTTTTCGCATATACA
AAACGTAATGTCATCCCTACTATAACTCAAATGAATCTTAAGTATGCCATTA
GTGCAAAGAATAGAGCTCGCACCGTAGCTGGTGTCTCTATCTGTAGTACT
ATGACCAATAGACAGTTTCATCAAAAATTATTGAAATCAATAGCCGCCACTA
GAGGAGCTACTGTAGTAATTGGAACAAGCAAATTCTATGGTGGTTGGCAC
AACATGTTAAAAACTGTTTATAGTGATGTAGAAAACCCTCACCTTATGGGTT
GGGATTATCCTAAATGTGATAGAGCCATGCCTAACATGCTTAGAATTATGG
CCTCACTTGTTCTTGCTCGCAAACATACAACGTGTTGTAGCTTGTCACAC
CGTTTCTATAGATTAGCTAATGAGTGTGCTCAAGTATTGAGTGAAATGGTC
ATGTGTGGCGGTTCACTATATGTTAAACCAGGTGGAACCTCATCAGGAGA
TGCCACAACTGCTTATGCTAATAGTGTTTTTAACATTTGTCAAGCTGTCAC
GGCCAATGTTAATGCACTTTTATCTACTGATGGTAACAAAATTGCCGATAAG
TATGTCCGCAATTTACAACACAGACTTTATGAGTGTCTCTATAGAAATAGAG
ATGTTGACACAGACTTTGTGAATGAGTTTTACGCATATTTGCGTAAACATTT
CTCAATGATGATACTCTCTGACGATGCTGTTGTGTGTTTCAATAGCACTTAT
GCATCTCAAGGTCTAGTGGCTAGCATAAAGAACTTTAAGTCAGTTCTTTAT
TATCAAAACAATGTTTTTATGTCTGAAGCAAAATGTTGGACTGAGACTGAC
CTTACTAAAGGACCTCATGAATTTTGCTCTCAACATACAATGCTAGTTAAAC
AGGGTGATGATTATGTGTACCTTCCTTACCCAGATCCATCAAGAATCCTAG
GGGCCGGCTGTTTTGTAGATGATATCGTAAAAACAGATGGTACACTTATGA
TTGAACGGTTCGTGTCTTTAGCTATAGATGCTTACCCACTTACTAAACATC
CTAATCAGGAGTATGCTGATGTCTTTCATTTGTACTTACAATACATAAGAAA

GCTACATGATGAGTTAACAGGACACATGTTAGACATGTATTCTGTTATGCTT
ACTAATGATAACACTTCAAGGTATTGGGAACCTGAGTTTTATGAGGCTATG
TACACACCGCATACAGTCTTACAGGCTGTTGGGGCTTGTGTTCTTTGCAA
TTCACAGACTTCATTAAGATGTGGTGCTTGCATACGTAGACCATTCTTATG
TTGTAAATGCTGTTACGACCATGTCATATCAACATCACATAAATTAGTCTTG
TCTGTTAATCCGTATGTTTGCAATGCTCCAGGTTGTGATGTCACAGATGTG
ACTCAACTTTACTTAGGAGGTATGAGCTATTATTGTAAATCACATAAACCAC
CCATTAGTTTTCCATTGTGTGCTAATGGACAAGTTTTTGGTTTATATAAAAAT
ACATGTGTTGGTAGCGATAATGTTACTGACTTTAATGCAATTGCAACATGT
GACTGGACAAATGCTGGTGATTACATTTTAGCTAACACCTGTACTGAAAGA
CTCAAGCTTTTTGCAGCAGAAACGCTCAAAGCTACTGAGGAGACATTTAA
ACTGTCTTATGGTATTGCTACTGTACGTGAAGTGCTGTCTGACAGAGAATT
ACATCTTTCATGGGAAGTTGGTAAACCTAGACCACCACTTAACCGAAATTA
TGTCTTTACTGGTTATCGTGTAACTAAAAACAGTAAAGTACAAATAGGAGA
GTACACCTTTGAAAAAGGTGACTATGGTGATGCTGTTGTTTACCGAGGTA
CAACAACTTACAAATTAAATGTTGGTGATTATTTTGTGCTGACATCACATAC
AGTAATGCCATTAAGTGCACCTACACTAGTGCCACAAGAGCACTATGTTAG
AATTACTGGCTTATACCCAACACTCAATATCTCAGATGAGTTTTCTAGCAAT
GTTGCAAATTATCAAAAGGTTGGTATGCAAAAGTATTCTACACTCCAGGGA
CCACCTGGTACTGGTAAGAGTCATTTTGCTATTGGCCTAGCTCTCTACTAC
CCTTCTGCTCGCATAGTGTATACAGCTTGCTCTCATGCCGCTGTTGATGCA
CTATGTGAGAAGGCATTAAAATATTTGCCTATAGATAAATGTAGTAGAATTAT
ACCTGCACGTGCTCGTGTAGAGTGTTTTGATAAATTCAAAGTGAATTCAAC
ATTAGAACAGTATGTCTTTTGTACTGTAAATGCATTGCCTGAGACGACAGC
AGATATAGTTGTCTTTGATGAAATTTCAATGGCCACAAATTATGATTTGAGT
GTTGTCAATGCCAGATTACGTGCTAAGCACTATGTGTACATTGGCGACCCT
GCTCAATTACCTGCACCACGCACATTGCTAACTAAGGGCACACTAGAACC
AGAATATTTCAATTCAGTGTGTAGACTTATGAAAACTATAGGTCCAGACATG
TTCCTCGGAACTTGTCGGCGTTGTCCTGCTGAAATTGTTGACACTGTGAG
TGCTTTGGTTTATGATAATAAGCTTAAAGCACATAAAGACAAATCAGCTCAA
TGCTTTAAAATGTTTTATAAGGGTGTTATCACGCATGATGTTTCATCTGCAA
TTAACAGGCCACAAATAGGCGTGGTAAGAGAATTCCTTACACGTAACCCT
GCTTGGAGAAAAGCTGTCTTTATTTCACCTTATAATTCACAGAATGCTGTA
GCCTCAAAGATTTTGGGACTACCAACTCAAACTGTTGATTCATCACAGGG
CTCAGAATATGACTATGTCATATTCACTCAAACCACTGAAACAGCTCACTC
TTGTAATGTAAACAGATTTAATGTTGCTATTACCAGAGCAAAAGTAGGCATA
CTTTGCATAATGTCTGATAGAGACCTTTATGACAAGTTGCAATTTACAAGTC
TTGAAATTCCACGTAGGAATGTGGCAACTTTACAAGCTGAAAATGTAACAG
GACTTTTTAAAGATTGTAGTAAGGTAATCACTGGGTTACATCCTACACAGG
CACCTACACACCTCAGTGTTGACACTAAATTCAAAACTGAAGGTTTATGTG
TTGACATACCTGGCATACCTAAGGACATGACCTATAGAAGACTCATCTCTA
TGATGGGTTTTAAAATGAATTATCAAGTTAATGGTTACCCTAACATGTTTATC
ACCCGCGAAGAAGCTATAAGACATGTACGTGCATGGATTGGCTTCGATGT
CGAGGGGTGTCATGCTACTAGAGAAGCTGTTGGTACCAATTTACCTTTAC

AGCTAGGTTTTTCTACAGGTGTTAACCTAGTTGCTGTACCTACAGGTTATG
TTGATACACCTAATAATACAGATTTTTCCAGAGTTAGTGCTAAACCACCGCC
TGGAGATCAATTTAAACACCTCATACCACTTATGTACAAAGGACTTCCTTG
GAATGTAGTGCGTATAAAGATTGTACAAATGTTAAGTGACACACTTAAAAAT
CTCTCTGACAGAGTCGTATTTGTCTTATGGGCACATGGCTTTGAGTTGAC
ATCTATGAAGTATTTTGTGAAAATAGGACCTGAGCGCACCTGTTGTCTATG
TGATAGACGTGCCACATGCTTTTCCACTGCTTCAGACACTTATGCCTGTTG
GCATCATTCTATTGGATTTGATTACGTCTATAATCCGTTTATGATTGATGTTC
AACAATGGGGTTTTACAGGTAACCTACAAAGCAACCATGATCTGTATTGTC
AAGTCCATGGTAATGCACATGTAGCTAGTTGTGATGCAATCATGACTAGGT
GTCTAGCTGTCCACGAGTGCTTTGTTAAGCGTGTTGACTGGACTATTGAA
TATCCTATAATTGGTGATGAACTGAAGATTAATGCGGCTTGTAGAAAGGTT
CAACACATGGTTGTTAAAGCTGCATTATTAGCAGACAAATTCCCAGTTCTT
CACGCTATTGCAAACCCTAAAGCTATTAAGTGTGTACCTCAAGCTGATGTA
GAATGGAAGTTCTATGATGCACAGCCTTGTAGTGACAAAGCTTATAAAATA
GAAGAATTATTCTATTCTTATGCCACACATTCTGACAAATTCACAGATGGTG
TATGCCTATTTTGGAATTGCAATGTCGATAGATATCCTGCTAATTCCATTGTT
TGTAGATTTGACACTAGAGTGCTATCTAACCTTAACTTGCCTGGTTGTGAT
GGTGGCAGTTTGTATGTAAATAAACATGCATTCCACACACCAGCTTTTGAT
AAAAGTGCTTTTGTTAATTTAAAACAATTACCATTTTTCTATTACTCTGACAG
TCCATGTGAGTCTCATGGAAAACAAGTAGTGTCAGATATAGATTATGTACC
ACTAAAGTCTGCTACGTGTATAACACGTTGCAATTTAGGTGGTGCTGTCTG
TAGACATCATGCTAATGAGTACAGATTGTATCTCGATGCTTATAACATGATG
ATCTCAGCTGGCTTTAGCTTGTGGGTTTACAAACAATTTGATACTTATAACC
TCTGGAACACTTTTACAAGACTTCAGAGTTTAGAAAATGTGGCTTTTAATG
TTGTAAATAAGGGACACTTTGATGGACAACAGGGTGAAGTACCAGTTTCT
ATCATTAATAACACTGTTTACACAAAAGTTGATGGTGTTGATGTAGAATTGT
TTGAAAATAAAACAACATTACCTGTTAATGTAGCATTTGAGCTTTGGGCTAA
GCGCAACATTAAACCAGTACCAGAGGTGAAAATACTCAATAATTTGGGTGT
GGACATTGCTGCTAATACTGTGATCTGGGACTACAAAAGAGATGCTCCAG
CACATATATCTACTATTGGTGTTTGTTCTATGACTGACATAGCCAAGAAACC
AACTGAAACGATTTGTGCACCACTCACTGTCTTTTTGATGGTAGAGTTGA
TGGTCAAGTAGACTTATTTAGAAATGCCCGTAATGGTGTTCTTATTACAGAA
GGTAGTGTTAAAGGTTTACAACCATCTGTAGGTCCCAAACAAGCTAGTCTT
AATGGAGTCACATTAATTGGAGAAGCCGTAAAAACACAGTTCAATTATTATA
AGAAAGTTGATGGTGTTGTCCAACAATTACCTGAAACTTACTTTACTCAGA
GTAGAAATTTACAAGAATTTAAACCCAGGAGTCAAATGGAAATTGATTTCTT
AGAATTAGCTATGGATGAATTCATTGAACGGTATAAATTAGAAGGCTATGCC
TTCGAA

- The sequences highlighted in yellow indicate unique MluI (ACGCGT) and BstBI (TTCGAA) restriction sites used to clone F4 (**Figure 1**).

**Fragment 5 (F5)**

TTCGAACATATCGTTTATGGAGATTTTAGTCATAGTCAGTTAGGTGGTTTAC
ATCTACTGATTGGACTAGCTAAACGTTTTAAGGAATCACCTTTTGAATTAGA
AGATTTTATTCCTATGGACAGTACAGTTAAAAACTATTTCATAACAGATGCG
CAAACAGGTTCATCTAAGTGTGTGTGTTCTGTTATTGATTTATTACTTGATG
ATTTTGTTGAAATAATAAAATCCCAAGATTTATCTGTAGTTTCTAAGGTTGTC
AAAGTGACTATTGACTATACAGAAATTTCATTTATGCTTTGGTGTAAAGATG
GCCATGTAGAAACATTTTACCCAAAATTACAATCTAGTCAAGCGTGGCAAC
CGGGTGTTGCTATGCCTAATCTTTACAAAATGCAAAGAATGCTATTAGAAA
AGTGTGACCTTCAAAATTATGGTGATAGTGCAACATTACCTAAAGGCATAAT
GATGAATGTCGCAAAATATACTCAACTGTGTCAATATTTAAACACATTAACA
TTAGCTGTACCCTATAATATGAGAGTTATACATTTTGGTGCTGGTTCTGATA
AAGGAGTTGCACCAGGTACAGCTGTTTTAAGACAGTGGTTGCCTACGGG
TACGCTGCTTGTCGATTCAGATCTTAATGACTTTGTCTCTGATGCAGATTC
AACTTTGATTGGTGATTGTGCAACTGTACATACAGCTAATAAATGGGATCT
CATTATTAGTGATATGTACGACCCTAAGACTAAAAATGTTACAAAAGAAAAT
GACTCTAAAGAGGGTTTTTTCACTTACATTTGTGGGTTTATACAACAAAAG
CTAGCTCTTGGAGGTTCCGTGGCTATAAAGATAACAGAACATTCTTGGAAT
GCTGATCTTTATAAGCTCATGGGACACTTCGCATGGTGGACAGCCTTTGT
TACTAATGTGAATGCGTCATCATCTGAAGCATTTTTAATTGGATGTAATTATC
TTGGCAAACCACGCGAACAAATAGATGGTTATGTCATGCATGCAAATTACA
TATTTTGGAGGAATACAAATCCAATTCAGTTGTCTTCCTATTCTTTATTTGAC
ATGAGTAAATTTCCCCTTAAATTAAGGGGTACTGCTGTTATGTCTTTAAAAG
AAGGTCAAATCAATGATATGATTTTATCTCTTCTTAGTAAAGGTAGACTTATA
ATTAGAGAAAACAACAGAGTTGTTATTTCTAGTGATGTTCTTGTTAACAACT
AAACGAACAATGTTTGTTTTTCTTGTTTTATTGCCACTAGTCTCTAGTCAGT
GTGTTAATCTTACAACCAGAACTCAATTACCCCCTGCATACACTAATTCTTT
CACACGTGGTGTTTATTACCCTGACAAAGTTTTCAGATCCTCAGTTTTACA
TTCAACTCAGGACTTGTTCTTACCTTTCTTTTCCAATGTTACTTGGTTCCAT
GCTATACATGTCTCTGGGACCAATGGTACTAAGAGGTTTGATAACCCTGTC
CTACCATTTAATGATGGTGTTTATTTTGCTTCCACTGAGAAGTCTAACATAA
TAAGAGGCTGGATTTTTGGTACTACTTTAGACTCGAAGACCCAGTCCCTA
CTTATTGTTAATAACGCTACTAATGTTGTTATTAAAGTCTGTGAATTTCAATT
TTGTAATGATCCATTTTTGGGTGTTTATTACCACAAAAACAACAAAAGTTGG
ATGGAAAGTGAGTTCAGAGTTTATTCTAGTGCGAATAATTGCACTTTTGAA
TATGTCTCTCAGCCTTTTCTTATGGACCTTGAAGGAAAACAGGGTAATTTC
AAAAATCTTAGGGAATTTGTGTTTAAGAATATTGATGGTTATTTTAAAATATAT
TCTAAGCACACGCCTATTAATTTAGTGCGTGATCTCCCTCAGGGTTTTTCG
GCTTTAGAACCATTGGTAGATTTGCCAATAGGTATTAACATCACTAGGTTTC
AAACTTTACTTGCTTTACATAGAAGTTATTTGACTCCTGGTGATTCTTCTTC
AGGTTGGACAGCTGGTGCTGCAGCTTATTATGTGGGTTATCTTCAACCTA
GGACTTTTCTATTAAAATATAATGAAAATGGAACCATTACAGATGCTGTAGA
CTGTGCACTTGACCCTCTCTCAGAAACAAAGTGTACGTTGAAATCCTTCA

CTGTAGAAAAAGGAATCTATCAAACTTCTAACTTTAGAGTCCAACCAACAG
AATCTATTGTTAGATTTCCTAATATTACAAACTTGTGCCCTTTTGGTGAAGT
TTTTAACGCCACCAGATTTGCATCTGTTTATGCTTGGAACAGGAAGAGAAT
CAGCAACTGTGTTGCTGATTATTCTGTCCTATATAATTCCGCATCATTTTCC
ACTTTTAAGTGTTATGGAGTGTCTCCTACTAAATTAAATGATCTCTGCTTTA
CTAATGTCTATGCAGATTCATTTGTAATTAGAGGTGATGAAGTCAGACAAAT
CGCTCCAGGGCAAACTGGAAAGATTGCTGATTATAATTATAAATTACCAGA
TGATTTTACAGGCTGCGTTATAGCTTGGAATTCTAACAATCTTGATTCTAAG
GTTGGTGGTAATTATAATTACCTGTATAGATTGTTTAGGAAGTCTAATCTCA
AACCTTTTGAGAGAGATATTTCAACTGAAATCTATCAGGCCGGTAGCACAC
CTTGTAATGGTGTTGAAGGTTTTAATTGTTACTTTCCTTTACAATCATATGG
TTTCCAACCCACTAATGGTGTTGGTTACCAACCATACAGAGTAGTAGTACT
TTCTTTTGAACTTCTACATGCACCAGCAACTGTTTGTGGACCTAAAAAGTC
TACTAATTTGGTTAAAAACAAATGTGTCAATTTCAACTTCAATGGTTTAACA
GGCACAGGTGTTCTTACTGAGTCTAACAAAAAGTTTCTGCCTTTCCAACA
ATTTGGCAGAGACATTGCTGACACTACTGATGCTGTCCGTGATCCACAGA
CACTTGAGATTCTTGACATTACACCATGTTCTTTTGGTGGTGTCAGTGTTA
TAACACCAGGAACAAATACTTCTAACCAGGTTGCTGTTCTTTATCAGGATG
TTAACTGCACAGAAGTCCCTGTTGCTATTCATGCAGATCAACTTACTCCTA
CTTGGCGTGTTTATTCTACAGGTTCTAATGTTTTCAAACACGTGCAGGCT
GTTTAATAGGGGCTGAACATGTCAACAACTCATATGAGTGTGACATACCCA
TTGGTGCAGGTATATGCGCTAGTTATCAGACTCAGACTAATTCTCCTCGGC
GGGCACGTAGTGTAGCTAGTCAATCCATCATTGCCTACACTATGTCACTTG
GTGCAGAAAATTCAGTTGCTTACTCTAATAACTCTATTGCCATACCCACAAA
TTTTACTATTAGTGTTACCACAGAAATTCTACCAGTGTCTATGACCAAGACA
TCAGTAGATTGTACAATGTACATTTGTGGTGATTCAACTGAATGCAGCAAT
CTTTTGTTGCAATATGGCAGTTTTTGTACACAATTAAACCGTGCTTTAACTG
GAATAGCTGTTGAACAAGACAAAAACACCCAAGAAGTTTTTGCACAAGTC
AAACAAATTTACAAAACACCACCAATTAAAGATTTTGGTGGTTTTAATTTTT
CACAAATATTACCAGATCCATCAAAACCAAGCAAGAGGTCATTTATTGAAG
ATCTACTTTTCAACAAAGTGACACTTGCAGATGCTGGCTTCATCAAACAAT
ATGGTGATTGCCTTGGTGATATTGCTGCTAGAGACCTCATTTGTGCACAAA
AGTTTAACGGCCTTACTGTTTTGCCACCTTTGCTCACAGATGAAATGATTG
CTCAATACACTTCTGCACTGTTAGCGGGTACAATCACTTCTGGTTGGACC
TTTGGTGCAGGTGCTGCATTACAAATACCATTTGCTATGCAAATGGCTTATA
GGTTTAATGGTATTGGAGTTACACAGAATGTTCTCTATGAGAACCAAAAAT
TGATTGCCAACCAATTTAATAGTGCTATTGGCAAAATTCAAGACTCACTTTC
TTCCACAGCAAGTGCACTTGGAAAACTTCAAGATGTGGTCAACCAAAATG
CACAAGCTTTAAACACGCTTGTTAAACAACTTAGCTCCAATTTTGGTGCAA
TTTCAAGTGTTTTAAATGATATCCTTTCACGTCTTGACAAAGTTGAGGCTG
AAGTGCAAATTGATAGGTTGATCACAGGCAGACTTCAAAGTTTGCAGACA
TATGTGACTCAACAATTAATTAGAGCTGCAGAAATCAGAGCTTCTGCTAAT
CTTGCTGCTACTAAAATGTCAGAGTGTGTACTTGGACAATCAAAAAGAGTT
GATTTTTGTGGAAAGGGCTATCATCTTATGTCCTTCCCTCAGTCAGCACCT

CATGGTGTAGTCTTCTTGCATGTGACTTATGTCCCTGCACAAGAAAAGAA
CTTCACAACTGCTCCTGCCATTTGTCATGATGGAAAAGCACACTTTCCTC
GTGAAGGTGTCTTTGTTTCAAATGGCACACACTGGTTTGTAACACAAAGG
AATTTTTATGAACCACAAATCATTACTACAGACAACACATTTGTGTCTGGTA
ACTGTGATGTTGTAATAGGAATTGTCAACAACACAGTTTATGATCCTTTGC
AACCTGAATTAGACTCATTCAAGGAGGAGTTAGATAAATATTTTAAGAATCA
TACATCACCAGATGTTGATTTAGGTGACATCTCTGGCATTAATGCTTCAGT
TGTAAACATTCAAAAAGAAATTGACCGCCTCAATGAGGTTGCCAAGAATTT
AAATGAATCTCTCATCGATCTCCAAGAACTTGGAAAGTATGAGCAGTATATA
AAATGGCCATGGTACATTTGGCTAGGTTTTATAGCTGGCTTGATTGCCATA
GTAATGGTGACAATTATGCTTTGCTGTATGACCAGTTGCTGTAGTTGTCTC
AAGGGCTGTTGTTCTTGTGGATCC

- Sequences highlighted in yellow represent unique BstBI (TTCGAA) and BamHI (GGATCC) restriction sites used to clone F5 (**Figure 1**).

- The single nucleotide highlighted in green represents a silent mutation introduced to remove a BstBI (TTCGAA) restriction site present in the viral spike (S) gene to clone F5 (**Figure 1**). This single nucleotide silent mutation was also used as a genetic tag to distinguish the rSARS-CoV-2 from the natural SARS-CoV-2 isolate.

**Venus-2A**

ATGGTGAGCAAGGGCGAGGAGCTGTTCACCGGGGTGGTGCCCATCCTG
GTCGAGCTGGACGGCGACGTAAACGGCCACAAGTTCAGCGTGTCCGGC
GAGGGCGAGGGCGATGCCACCTACGGCAAGCTGACCCTGAAGCTGATC
TGCACCACCGGCAAGCTGCCCGTGCCCTGGCCCACCCTCGTGACCACC
CTGGGCTACGGCCTGCAGTGCTTCGCCCGCTACCCCGACCACATGAAGC
AGCACGACTTCTTCAAGTCCGCCATGCCCGAAGGCTACGTCCAGGAGCG
CACCATCTTCTTCAAGGACGACGGCAACTACAAGACCCGCGCCGAGGTG
AAGTTCGAGGGCGACACCCTGGTGAACCGCATCGAGCTGAAGGGCATC
GACTTCAAGGAGGACGGCAACATCCTGGGGCACAAGCTGGAGTACAACT
ACAACAGCCACAACGTCTATATCACCGCCGACAAGCAGAAGAACGGCAT
CAAGGCCAACTTCAAGATCCGCCACAACATCGAGGACGGCGGCGTGCA
GCTCGCCGACCACTACCAGCAGAACACCCCCATCGGCGACGGCCCCGT
GCTGCTGCCCGACAACCACTACCTGAGCTACCAGTCCAAGCTGAGCAAA
GACCCCAACGAGAAGCGCGATCACATGGTCCTGCTGGAGTTCGTGACCG
CCGCCGGGATCACTCTCGGCATGGACGAGCTGTACAAAGGTCCGGAG
CCACGAACTTCTCTCTGTTAAAGCAAGCAGGGGACGTGGAAGAAAACCC
CGGTCCT

- The underlined sequence indicates the porcine Teschovirus-1 (PTV-1) 2A self-cleaving peptide.
